# Perilous coexistence: *Chilli Leaf Curl Virus* and *Candidatus Phytoplasma trifolii* infecting *Capsicum annuum*, India

**DOI:** 10.1101/2022.12.17.520842

**Authors:** Vineeta Pandey, Aarshi Srivastava, Smriti Mall, Abdullah M. Al-Sadi, Muhammad Shafiq Shahid, R. K. Gaur

## Abstract

Molecular computing was used to investigate the possible causal agents of chilli crop samples showing mixed symptoms of yellow leaf curl and little leaf type diseases in the Uttar Pradesh province, India. Total genomic DNA was extracted from twenty-five samples and amplified by PCR using a universal primer pair for begomovirus and phytoplasma. Mixed infection samples show positive amplified products for begomovirus (DNA-A and betasatellite) and phytoplasma (16S rRNA and Sec A). The identified begomovirus from chilli samples was identified as a strain isolate of the previously described Chilli Leaf Curl Virus (94.2% nucleotide sequence identity), which is known to infect *Solanum lycopersicon*, in Oman, whereas the 16S rRNA was identified from the source *Candidatus Phytoplasma trifolii* (99.04% nucleotide sequence identity), which is known to infect Helichrysum flowering plants in India. Subsequently, molecular computing research based on phylogenetic interweaves, putative recombination, amino acid selection, and genetic diversity were investigated, revealing divergent evolutionary patterns with significant variation and recombination events. The majority of the sequence variations observed in begomovirus and phytoplasma were caused via inter- and intra-specific recombination. These findings could be the first *in silico* combined infection analysis of ChiLCV and *Ca.P.trifolii* in a chilli crop in India, revealing the potential adaption and evolution of begomovirus and phytoplasma to a new geographic range and crop.

## Introduction

*Solanaceae*, a family of flowering and fruiting plants, comprising crops of regional significance, are severely affected by many plant viruses; begomoviruses are identified as the primary limiting pathogen (Pandey et. al., 2021). *Capsicum annuum L*. (family *Solanaceae*), is grown in tropical and sub-tropical countries. India is one of the topmost cultivators of chilli crops globally, based on the FAOSTAT statistical report for the year 2020, it is grown on 0.683 million hectares annually (https://www.fao.org/faostat/en/#data/QCL) and yielding 2.05 million tonnes, according to the APEDA 2020-21 data (Agricultural and Processed Food Products Export Development Authority) (https://agriexchange.apeda.gov.in/India%20Production/_India_Productions.aspx?_hscode=1098). The cultivars of chilli are susceptible to a wide array of pathogens and diseases. Among the important biotic factors that limit the productivity of chilli crops in various regions of the globe are begomoviruses and phytoplasma (Pandey et al., 2022; Mishra et al., 2022; Lavanya and Arun, 2021; Pandey et al., 2021; Mishra et al., 2020; Kumari et al., 2019, Ganefianti et al., 2017)‥ In the chilli plant, infection of begomovirus has been described in distinct states of India and was revealed to cause severe yielding loss (Khan and Khan, 2017). Symptoms of varying severity were observed in plants, ranging from vein yellowing of leaves, mosaic, mottling, stunting, curling, flower abortion, distortion, and small-unvendible fruits reported in Rojas et al., 2018 study. Plant pests and pathogens are some of the foremost factors that confined the production rate of crops. Begomovirus is a hazardous plant virus of the largest family of *Geminiviridae* that constitutes extremely devastating universal phytopathogens (Gibbs and Ohshima, 2010; Scholthof et al., 2011).

Begomoviruses have a circular single-stranded DNA genetic constituent and either a (DNA-A) monopartite genome component that encodes for several important proteins involved in virus long-distance and cell-to-cell movement, virus replication, encapsidation, transcription, and suppression of host silencing genes or a (DNA-A and DNA-B) bipartite genome component that plays an important role in enlarging host range, virus movement, and symptom expression in the plant (Marwal et al., 2021; Srivastava et al., 2022). As a consequence of the rapid spread of the vector whitefly (*Bemisia tabaci*), economically significant crops are infected (Moffat, 1999; Gutierrez 2000). In general, betasatellites are the most commonly associated satellite with monopartite begomoviruses, followed by alphasatellites and, in very few cases, deltasatellites. (Briddon et al., 2018; Mubin et al., 2020; Fiallo-Olivé et al., 2016). Betasatellite melloencodes for βC1 protein on the complementary sense strand and has a significant task in the induction of symptoms, and post-transcriptional and transcriptional gene silencing (Zhou et al., 2013; Patil et al., 2010). Its size is half the size of the DNA-A component. The Geminiviridae family, which included fourteen genera (Fiallo-Olive et al., 2021) and ~520 species, was identified in 1978 by Goodman, 1981. Chilli leaf curl virus (ChiLCV) is a prevalent monopartite begomovirus that causes significant damage, including solanaceous and non-solanaceous hosts when combined with various betasatellites (Malathi et al., 2017). The ChiLCV infection has resulted in 14–100% chilli yield losses in Rajasthan (India) (Kumar et al., 2015; Senanayake et al 2012), causing substantial economic losses across India’s tropical and subtropical regions.

On the other scenario, Phytoplasmas are phloem-limited pleomorphic plant-pathogenic prokaryotes which are housed in the class *Mollicutes* and cannot be cultured under axenic conditions. The occurrence of phytoplasmas and their related infections is an emerging danger to crop productivity, resulting in considerable yield losses globally. They are transmitted in natural conditions mainly by phloem-sucking vectors such as leafhoppers and could also be transmitted through seeds, plant propagation materials, and grafting (Amaral-Mello et al., 2006). Plants exhibit various levels of symptoms, which are characterised by little leaves, big buds, flower virescence, phyllody, and witches’ brooms (Ermacora and Osler, 2019). And over 45 plant species have been related to phytoplasma infections in India., including vegetables, fruit, ornamental and other agriculturally important crop species (Chaturvedi et al., 2010; Rao et al., 2011). Among them, the little leaf is the utmost widespread disease, causing 100 % yield loss and is reported when conditions are favourable for phytoplasmas (Rao et al., 2021). Based on 16S rRNA sequence analysis, ten different groupings such as 16SrI, 16SrII, 16SrV, 16SrVI, 16SrIX, 16SrXI, and 16SrXIV are primarily associated with various types of infected vegetable crops identified throughout India. Among them, the most prevalent phytoplasma groups are Aster yellows (16SrI), followed by the ‘peanut witches’ broom (16SrII), which have been linked with more than 31 plant diseases (Mall et al., 2011; Rao et al., 2017) of different cultivated and non-cultivated plants, and most of the groups were reported from the northeastern regions of India. The clover proliferation group’s ‘Ca. Phytoplasma trifolii’ and closely related phytoplasma strains typically damage dicots and monocots plants. Leguminous, solanaceous, and brassica crops are among the most usually afflicted dicotyledonous hosts (Lee et al., 2004; Girsova et al., 2017; Kumari et al., 2019).

Plant viruses are currently a major constraint in tropical and subtropical nations and have greatly limited the production of many fibre and vegetable crops (Mishra et al., 2020; Sohrab et al., 2016c; Sohrab et al., 2016b). Agriculture crop expansions and overlaps enhance the host ranges of insect vectors and begomoviruses, resulting in a laborious effort in disease management due to limited availability of eco-friendly chemical control techniques and the overlap availability of resistance sources. The previous literature has well documented the natural co-existence of phytoplasma and viruses in the identical hosts of many crop plants (Lebsky et al., 2011; Aljanabi et al., 2008; Kaminska et al., 2005). Therefore, to access the disease incidence in chilli crops, a survey was conducted during 2020-21 across the Gorakhpur region in the Uttar Pradesh state of India. The infected chilli plant showed mosaic, little leaf and leaf curl symptoms, which were identified predominately along with the combination of vectors population such as whitefly and leaf hoppers. The characteristic disease symptoms and the presence of vectors on chilli plants encouraged us about a mixed infection of phytoplasma and begomovirus. Understanding the molecular mechanism of begomovirus-phytoplasma association, epidemiology, interaction, and other factors invoking growth and progression is crucial. The accessibility of phytoplasma detection molecular approaches at the strain level has significantly facilitated the development and application of integrated disease management programmes (Kumari et al., 2019). Therefore, the current study was carried out to validate the natural co-existence of begomovirus and phytoplasma. Moreover, we also compare the evolutionary dynamics among them.

## Materials and Methods

### Survey and collection of diseased chilli leaf

A Sample survey was conducted in 34 different farmer’s fields in Gorakhpur, Uttar Pradesh, India during 2020–21 with combined symptoms of leaf curl and little leaf symptoms on chilli. 25 samples (IC-1 to IC-25) of chilli with both little leaf and leaf curl symptoms were collected from various farmer fields in the Gorakhpur region. The chilli plants permitted for sample collection were those with the typical mixed symptoms of the little leaf along with leaf curling for phytoplasma and begomovirus, respectively. Besides the symptomatic sample, four asymptomatic chilli plants (HC-1 to HC-4), devoid of any symptoms, were also taken in this study.

### DNA isolation and begomovirus detection

From asymptomatic and symptomatic chilli leaf samples, total genomic DNA was isolated employing the CTAB (Cetyl Trimethyl Ammonium Bromide) procedure (Sahu et al., 2018). The occurrence of a begomovirus in chilli plant samples was detected using PCR amplification using coat protein (CP) gene, DNA-B, and associated satellite molecules specific primers such as PAL1v1978 /PAR1c496 for coat protein (CP) (Rojas, 1993), DNABLC1/DNABLV2 for DNA-B (Green et al., 2001), Beta01/Beta02 for betasatellite (Bull et al., 2003) and DNA 101/DNA102 (Briddon et al., 2002) for alphasatellite (Srivastava et al., 2020a). The absence of DNA-B and alphasatellites was confirmed by PCR using their respective primers. For amplifying the circular DNA-A, the PCR amplified eleven positive virus genomes were amplified by RCA (rolling circle amplification) using the RCA-based TempliPhi DNA amplification kit (GE Healthcare) following the manufacturer’s instructions. The RCA-positive products were restriction digested, cloned, purified, and sequenced at an ABI automated sequencer at The Centre for Genomic Application in New Delhi by using Sanger’s Method before being submitted to NCBI, GenBank.

### Detection and sequence characterization of a phytoplasma

The same DNA samples used to test for begomoviruses were also used to screen for phytoplasma in the chilli plant. The presence of phytoplasma was done by the first round of PCR, the universal primer set P1/P7 (Deng and Hiruki, 1991; Smart et al., 1996) and then was followed by a second round of nested PCR amplification using the primer pair R16F2n/R16R2n (Gundersen and Lee, 1996). The PCR reaction mixture was set up with purified DNA in a thermocycler (Eppendorf, Germany) under the following conditions with a total of 35 cycles: 94 °C for 30 s, 53 °C for 60 s, and 72 °C for 60 s, with a final extension step of 72 °C for a 10 min run programme. The first round of PCR’s positive amplicon, which contained 1 μL, serves as the template for the second round of PCR. Additionally, the SecA gene’s first and second rounds of PCRs used identical reaction conditions with primer pairs SecAfor1/SecArev3 (Hodgetts et al., 2008) and SecAfor2/SecArev3 (Lee et al., 2010b). The PCR amplified products of the chilli plant phytoplasma 16SrDNA (1.25 kb) and SecA gene (0.48 kb) were excised from the gel, cleaned, and purified using the Wizard SV Gel and PCR Clean-R19 purification systems following the instructions by the manufacturer (Promega, Madison, USA). The positive PCR amplicons were bidirectionally sequenced and gathered using BioEdit software (Hall, 1999). The CLUSTAL W approach from the MEGA X programme was applied to align 16S rRNA and SecA sequences with closely similar sequences already in NCBI databases, and the consensus sequences of both independent gene sequences were deposited to NCBI, GenBank.

### *In-silico* enzyme digestion of phytoplasma

Sequences of the 16S rRNA gene related to *Ca.P.trifolii* strains were exposed to virtual RFLP restriction enzyme analysis employing the iPhyClassifier tool (Zhao et al., 2009). The F2nR2 segment was digested by seventeen restriction enzymes, that are used to classify phytoplasmas into various categories and subgroups (Wei et al., 2008). Following in-silico restriction digestion, a computerized gel electrophoresis image (3.0% agarose) was generated. The PCR-RFLP pattern with the key enzymes is used to differentiate the pattern from formerly recognised group/subgroup patterns, which helps for fine-scale differentiation of chilli plant strains within the clover proliferation group (16SrVI).

### Sequence analysis

The newly obtained sequences for Chilli Leaf Curl Virus-DNA-A (ChiLCV-A), all 6 ORFs and Chilli Leaf Curl Betasatellite (ChLCuB) was evaluated for sequence homology with all the sequences accessible utilising the BLASTn technique in the GenBank database (http://www.ncbi.nlm.nih.gov) (Altschul et al., 1990). Furthermore, NCBI database sequences with the maximum similarity to ChiLCV-A and ChLCuB were retrieved. Furthermore, using the SDT (Sequence Demarcation Tool) (v1.2) and the MUSCLE approach, all datasets for begomovirus and all six ORFs were used to determine pairwise sequence identities between amino acid sequences (Muhire et al., 2014).

By using the BLASTn program in the NCBI database, the same highest similarity sequence retrieval approach was employed for phytoplasma 16S rRNA and Sec A genes (Altschul et al., 1990), and the SDT v1.2 with MUSCLE technique has been used for pairwise sequence identities among amino acid sequences (Muhire et al., 2014).

### Phylogenetic Analysis

Subsequently, the best substitution model for each of the sequence datasets of begomovirus was found based on the lowest Bayesian Information Criterion (BIC) score using MEGA X (Kumar et al., 2018) and for phylogenetic analysis of all 6 ORFs using each ORF sequence for every isolates taken for DNA-A and betasatellite phylogenetic tree construction **(Table S1 and S2)**, we used the Maximum Likelihood Method with 1000 bootstrap replications employing MEGA X. Whereas the MCC (maximum clade credibility) phylogenetic tree was constructed using the Bayesian method and Tree annotator tool accessible in BEAST v.1.10 (Suchard et al., 2018) and the resulting tree was edited and Interactive Tree Of Life (iTOL) v.6.5 was used to develop a figure (https://itol.embl.de/#) (Letunic et al., 2021) for ChiLCV-A and ChLCuB. Besides that, the transition/transversion rate and its bias were also evaluated using the MEGA X programme for each of the sequence datasets of begomovirus (Kumar et al., 2018).

Afterwards, the optimum substitution model for the sequence dataset of phytoplasma, i.e., 16S rRNA and Sec A genes, was found using MEGA X based on the lowest BIC score (Kumar et al., 2018). The MCC phylogenetic tree was constructed using the Bayesian approach and the Tree annotator tool, both of which are accessible in BEAST v.1.10 (Suchard et al., 2018). iTOL v.6.5 was used to view and edit the resulting tree (Letunic et al., 2021).

### Recombination Analysis

The neighbour net method of Splits Tree v.4 (Huson et al., 2006) software was used to characterize the reticulate phylogenetic network for both begomovirus components (ChiLCV-A and ChLCuB) and phytoplasma datasets that provide evidence for recombination event. Furthermore, recombination analysis was carried out with the default setting and the highest permissible Bonferroni corrected p-value of 0.05 using seven different algorithms implemented in RDP4.1, including RDP, BOOTSCAN, GENECOV, CHIMARA MaxChi, SISCAN, and 3SEQ (Martin et al., 2015). Among the seven approaches, events with at least three algorithms were utilised to avoid false-positive results and explore proper recombination (Mishra et al., 2020) among begomovirus sequence datasets for DNA-A, 6 ORFs, and betasatellite. The same approach was used to examine recombination in two phytoplasma datasets, 16S rRNA and Sec A gene.

### Computation of nucleotide substitution rate

The Bayesian Markov Chain Monte Carlo (MCMC) of BEAST (v.1.10) (Suchard et al., 2018) was used to assess the mean substitution rate and rate of mutation at 3 distinct codon positions for both the ChiLCV-A and ChLCuB sequence datasets of begomovirus. This uses coalescent constant demographic models and best-fit clock to effectively achieve sample size through the Tracer programme (v.1.5) (Rambaut et al., 2018). The MCMC chain analysis was run for 10^7^ with 10% burn-in to obtain a 95% highest probability density (HPD) value. A similar procedure was used to determine the nucleotide substitution rate across all phytoplasma datasets.

### Population Demography Analysis

A population demographic analysis was carried out to inspect the nucleotide polymorphism based on different parameters of genetic diversity such as the total number of mutations (η), the number of polymorphic sites (S), nucleotide diversity (π), the average of nucleotide differences (k), the total number of mutations (θ – η), Watterson’s estimate of the population mutation rate based on the total number of segregation sites (θ - w), number of haplotypes (h) and haplotype diversity (Hd) by using DnaSP v. 6.0 (Rozas et al., 2017). Different types of Neutrality tests such as Tajima’s D (nucleotide diversity with the proportion of polymorphic sites), Fu and Li’s F* (difference between the number of singletons and the average number of nucleotide differences between paired sequences), and Fu and Li’s D* (difference between the number of singletons and the total number of mutations) tests were also performed and offered in DnaSP v.6.0 for a sequence of both begomovirus components (DNA-A, 6 ORFs and betasatellite)

Analysis of population demographics for 16S rRNA and Sec A was estimated for all the above attributes of genetic diversity and three neutrality tests by using DnaSP v. 6.0 (Rozas et al., 2017).

### Selection analysis

To comprehend the potential impact of positive and negative selection pressure on the unequal distribution of variation seen between ChiLCV-A, 6 ORFs, and ChLCuB, the ratio of non-synonymous to synonymous (dN/dS) substitutions was determined using standard MEGA X (Kumar et al., 2018) parameters. Negative (purifying), positive (diversifying), and neutral selection pressures are indicated by the ratios dN/dS < 1, dN/dS > 1, and dN/dS = 1, respectively. Using three methods, datamonkey (www.datamonkey.org) was utilised to detect the sites evolved under negative (Purifying) and positive (Diversifying) selection: fast unbiased Bayesian approximation (FUBAR), single-likelihood ancestor counting (SLAC), and fixed-effects likelihood (FEL) (Weaver et al., 2018).These selection pressure assessments were also undertaken on components of phytoplasma using MEGA X (Kumar et al., 2018) and Datamonkey (www.datamonkey.org) (Weaver et al., 2018).

## Result

### Identification, sequencing and Species distinction

During the years 2020–2021, twenty-five chilli leaf samples exhibiting little leaf and curling infection symptoms were taken from thirty-four different locations in Gorakhpur, Uttar Pradesh, India. To explore the causes of symptoms and infection in the chilli plant, whole DNA was isolated from both symptomatic and asymptomatic leaves **(Figure. 1)**. Additionally, begomovirus-specific PCR amplification was done to confirm the virus’s presence using a universal degenerate PAL1v1978/PAR1c496 primer (Rojas et al., 1993). From infected leaf samples, specific DNA segments of 1.1 kb were amplified. We also tested for the possible DNA-B component and an alphasatellite molecule, but no evidenced was found. In order to obtain the whole genome, eleven PCR-positive samples were treated with RCA (Haible et al., 2006). The restriction enzymes were then used to digest the RCA products *Bam*HI/*Hind*III (ChiLCV-A) and *Kpn*I/*Sac*I (ChLCuB), yielding 2.7 and 1.3 kb DNA fragments only from symptomatic samples, not from healthy plants. Simultaneously, the positive RCA products were cloned further into pUC19 vector to produce the recombinant plasmids pUCCLV-A and pUCCLV-β, which are known as ChiLCV-A/GKP and ChLCuB /GKP.

**Figure 1.**
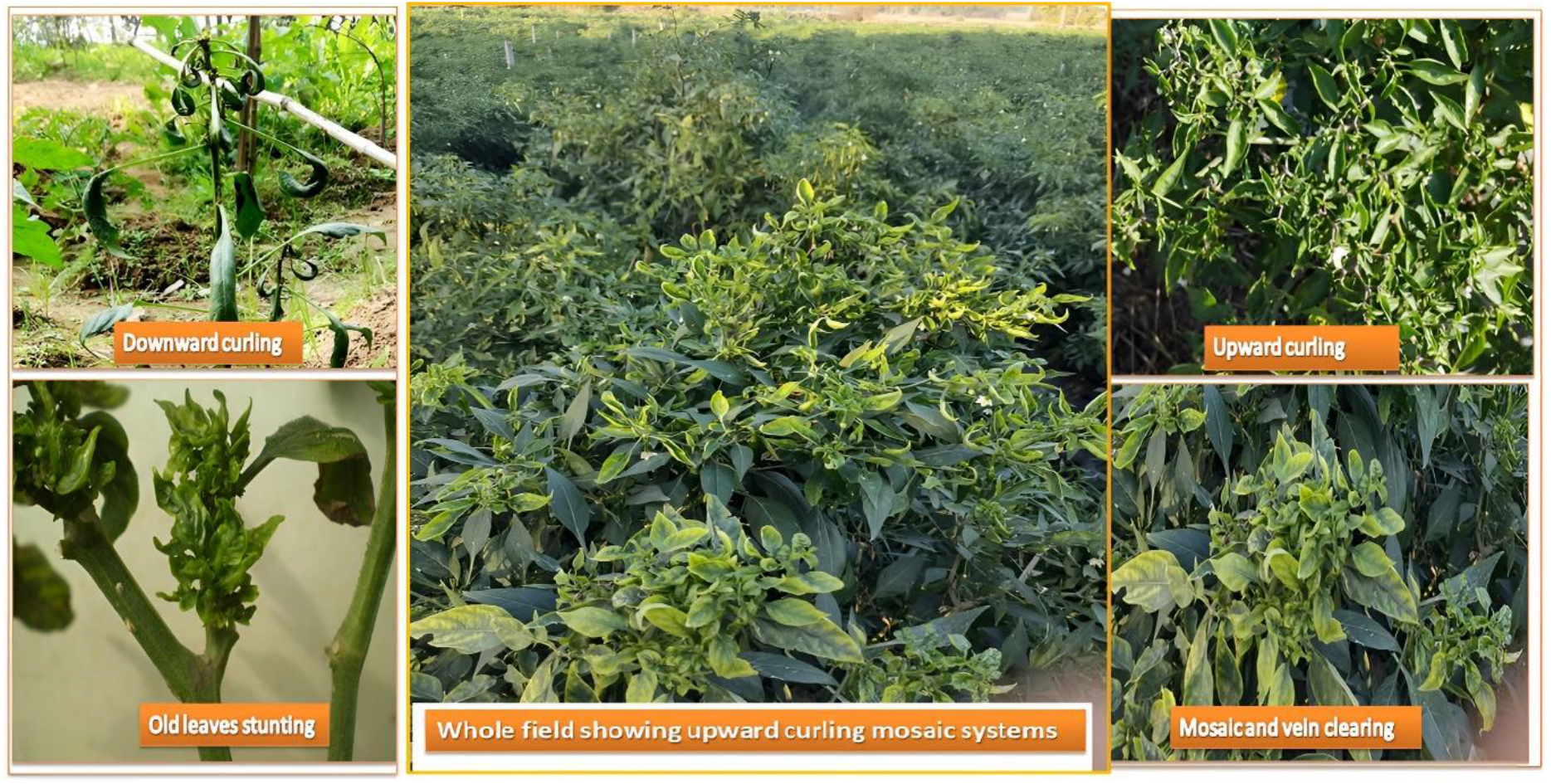
Chilli leaf samples exhibiting begomoviral symptoms including both downward and upward curling, mosaic and vein clearing, and some sharing the common symptoms of begomovirus and phytoplasma such as leaf miniature/stunting.

Following the study, we identified little leaf symptoms like those reported earlier during Phytoplasma infections. As a result, all collected samples were tested for phytoplasmas. Subsequently, for 16S rRNA, P1/P7 primer pair was employed for the first-round amplification, and we obtained a 1.8kb amplified PCR product that was treated to a specific set of primers, i.e., R16F2n/R16R2 for the second round of nested PCR amplification, and we obtained a final 1.25kb amplified product. Similarly, the Sec A gene was amplified using SecAfor1/SecArev3 for the first round and SecAfor1/SecArev3 for the second round of nested PCR, yielding 0.83 kb and 0.48 kb of the amplified product, respectively. Only one leaf sample (IC-21) showed association of begomovirus with phytoplasma from Belwar, Gorakhpur, Uttar Pradesh, India.

The positive cloned products for begomovirus (DNA-A and betasatellite) and phytoplasma (16S rRNA and Sec A) were sequenced using Sanger’s Method at ABI automated sequencer at the Centre for Genomic Application in New Delhi. To assess nucleotide similarity with other isolates, BLASTn was used on full genome sequences of DNA-A and betasatellite. The DNA-A had the highest nucleotide identity of 94.2% with *Chilli leaf curl India virus-*India (Acc. No. MK757213), whereas the betasatellite had the maximum sequence identity of 97.801% with *Chilli leaf curl betasatellite* (Acc. No. MT385295) (**Table S1**).

Furthermore, the ORF Finder programme was used to identify open reading frames (ORFs) in both DNA-A and betasatellite molecules, revealing the characteristics of genomic sequences more towards the monopartite genome of isolated begomovirus (**Figure. 2**) (**Table S1**). As a result, the genomic sequence of a monopartite GKP isolate (ChiLCV) and related satellite DNA were submitted to the NCBI GenBank database under the accession numbers MZ540908 (DNA-A) and MZ540909 (betasatellite). The NCBI database yielded the most nucleotide alignment matches, 61 for DNA-A and 42 for betasatellite (**Table S2 and Table S3**). The current study’s sequences demonstrated that there could be substantial recombination and mutation depending on the highest identity index of alignment. The MUSCLE technique of SDT v.1.0 software (Muhire et al., 2014) was used to calculate the percentage pairwise identity at the transcript level between the sequences of selected begomovirus and ORFs sequences of isolates ChiLCV-GKP and ChLCuB-GKP. This investigation at the amino acid level also indicates 87.8 % sequence identity for ChiLCV (Acc. No. MK757213) and 50.6 % amino acid sequence identity with ChLCuB (Acc. No. DQ343289) (**Table S4 and Table S5**) (**Figure S1**).

**Figure 2.**
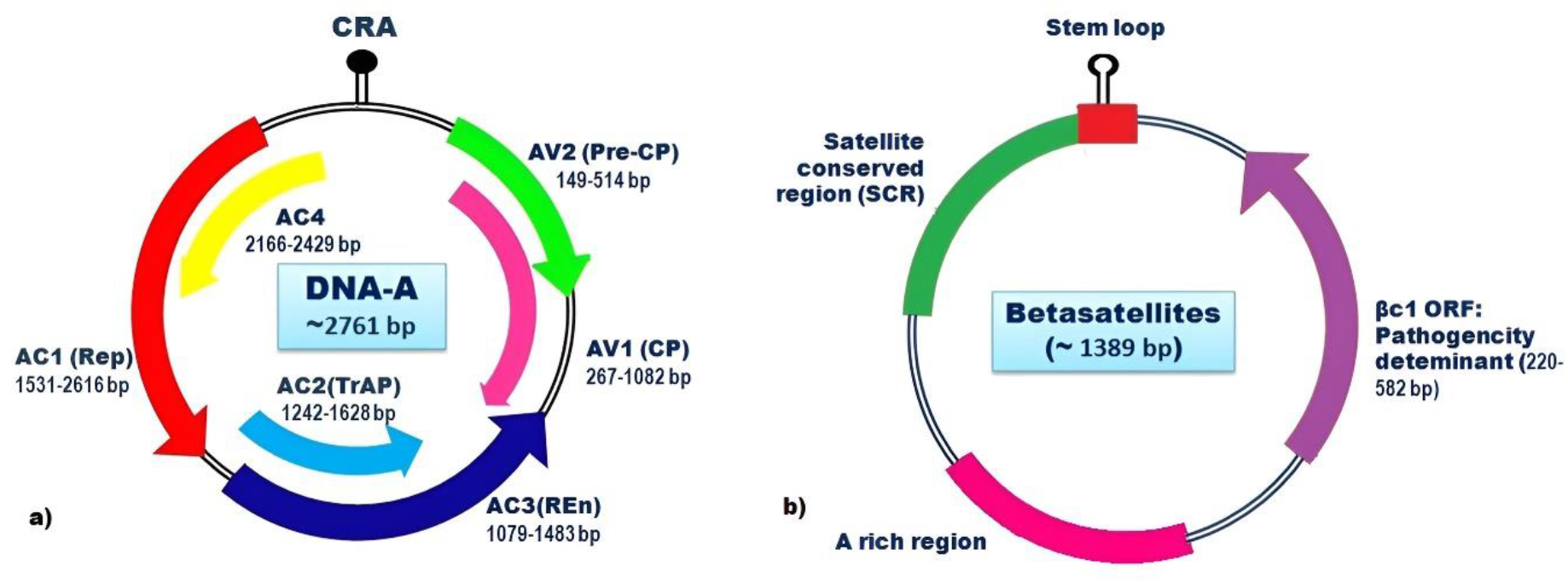
The genome organisation of the identified *Chilli Leaf Curl Virus* DNA-A molecule and associated betasatellite; a) indicating the range of all 6 ORFs of the ChiLCV-A GKP isolate AV1 (Coat protein: 267–1082 bp), AV2 (Pre-Coat protein: 149–514 bp), AC1 (Replication initiator protein: 1531–2616 bp), AC2 (Transcription activator protein:1242-1628bp), AC3 (Replication enhancer protein: 1079–148bp), AC4 (2166–2429bp) and Common Region (CR); b) organisation of the related betasatellite C1 (Pathogenicity determinant: 220–582bp), Adenine-rich area, and Satellite conserved region.

Similarly, BLASTn was employed for phytoplasma sequences to find entire genome sequence identities of 16S rRNA and the Sec A gene. The nucleotide identity of the 16S rRNA gene with the *Candidatus Phytoplasma trifolii* clone HSD2 16S ribosomal RNA gene (Acc. No. MN861370) was 99.04 per cent, but the Sec A gene has 98.3 per cent sequence identity with the Brinjal little leaf phytoplasma isolate MS-1 (Sec A) gene (Acc. No. KR906730) (**Table S6 and Table S7**). As a result, the sequences for the 16S rRNA and the Sec A gene have been submitted to the NCBI GenBank database, with accession numbers MZ557805 (16S rRNA) and MZ620707 (Sec A gene) (Sec A gene). Furthermore, the MUSCLE technique of SDT v.1.0 (Muhire et al., 2014) was applied to evaluate the maximum pairwise identity between the amino acid sequences of selected 16S rRNA phytoplasma sequences, exhibiting 97.4% maximum identity with *Ca.P. trifolii/HSD2/16VI-A:19* (Acc. No. MN861370), whereas Sec A genes show the highest amino acid identity with exceptionally different sequences, i.e. Brinjal little leaf/secA:15 (Acc. No. KT335271) **(Table S8 and Table S9) (Figure S1)**.

### *In-silico* enzyme digestion classification of 16S group/subgroup

In silico enzyme digestion was created from the 16S rDNA F2nR2 fragment of a *Ca. P. trifolii* clone, HSD2 (Acc. No. MZ557805), which was unique from all previously known 16Sr groups and subgroups. The 16Sr group VI, subgroup D reference pattern with the highest level of similarity (GenBank accession: X83431). Within 16Sr group VI, this strain represents a novel subgroup. Because of differences in the *Mse I* restriction profile, this RFLP pattern identified the phytoplasma strain that infects chilli as a novel strain in the subgroup 16SrVI-D (**Figure 3**).

**Figure 3.**
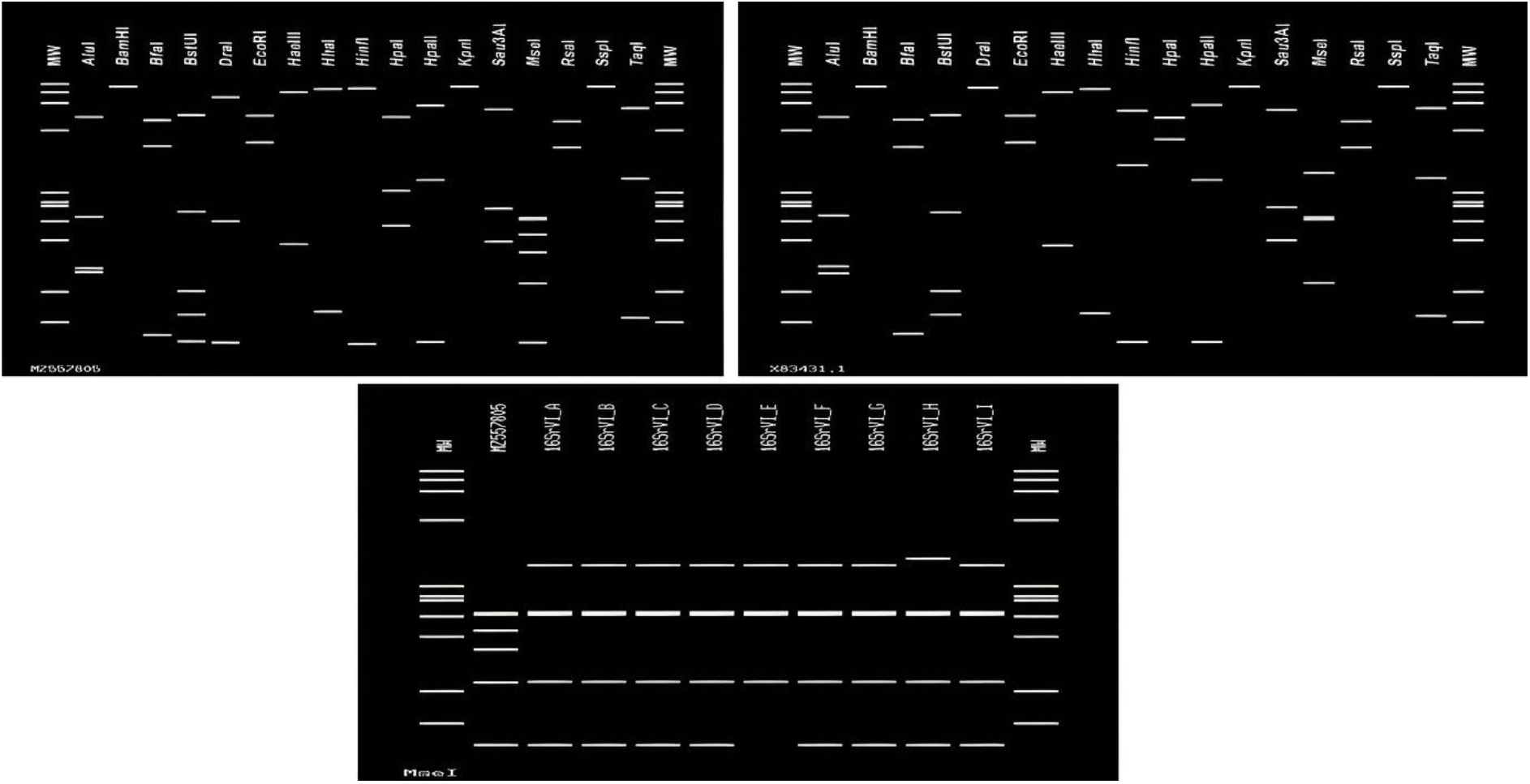
*In silico* RFLP was done on F2n/R2 segments of the *Ca. P.trifolii* clone HSD2 (Acc. No. MZ557805) with the following restriction endonuclease enzymes (left to right): AluI, BamHI, BfaI, BstUI, DraI, EcoRI, HaeIII, HhaI, HinfI, HpaI, HpaII, KpnI, Sau3AI, Mse1 RsaI, SspI Taq I). Mse1 virtual RFLP patterns from *Ca. P.trifolii* were used to distinguish strains of diverse phytoplasma strains from India belonging to strains in group 16SrVI. The reference pattern of the 16Sr group VI, subgroup D (GenBank accession: X83431) was the most similar. In-silico electrophoresis through 3% agarose gel was employed to separate the restriction fragments.

### Phylogenetic and transition/transversion bias (R) analysis

Meanwhile, using the MEGA X programme, evolutionary divergence was evaluated for ChiLCV datasets based on multiple sequence alignment and the most appropriate nucleotide substitution model, TN93+G for ChiLCV-A and GTR+G for ChLCuB, was opted because of the lowest BIC score. We also assessed the optimum nucleotide substitution model for all ORFs of DNA-A and betasatellite, namely T92+G+I for AV1, JC+G for AV2, T92+G for AC1, K2+G for AC2, JC+G for AC3, K2+G for AC4, and T92+G for βC1 (**Table 1**). This was then applied to generate maximum likelihood phylogenetic trees for all ORFs using MEGA X with bootstraps of 1000. This contributes to our understanding of divergence among individual ORFs, and we observed that all ORFs were clustered with different isolates. For instance, AV1, AV2, AC1, AC2, AC3 and AC4 show grouping with ChiLCV-[IN:MM1:17] (Acc. No. MF737343), ChiLCV-[IN:ND:chi:16] (Acc. No. KX533940), ChiLCV-[IN:Noi:Pap:09] (Acc. No. HM140371), ChiLCV-[Pak:CapAS2:Chi:13] (Acc. No. KM023148), ChiLCV-[IN:MM1:17] (Acc. No. MF737343) and ChiLCV-[IN:Noi:Pap:09] (Acc. No. HM140371), whereas for βC1 it was ChLCuB/[IN:KAN-05:sal spl:20 (Acc. No. MT385295) (**Figure S2**). Although MCC phylogenetic analysis using the BEAST v.1.10 software package (Suchard et al., 2018) and the (iTOL) v.6.5 (https://itol.embl.de/#) (Letunic et al., 2021) visualisation tool for DNA-A and betasatellite datasets, we observed ChiLCV-[IN: MM1:17] (Acc. No. MF737343) clustered with our ChiLCV-A (Acc. No MZ540908) (**Figure 4a**), whereas the ChLCuB isolates grouped with ChLCuB/[IN: KAN-05:sal spl:20] (Acc. No. MT385295) isolate from Kanpur (**Figure 4b**) which was different from βC1 clustered isolate. In addition, the transition/transversion bias (R) of each ChiLCV dataset was evaluated. The range of transitional and transversional substitution rates for ChiLCV-A was 10.58–15.95 and 5.04– 7.29, respectively, with a calculated transition/transversion bias (R) for DNA-A of 0.96 (Table 10). Similarly, the transitional and transversional substitution rates for ChLCuB were 7.22– 14.07 and 5.43–9.69, respectively, with a calculated transition/transversion bias (R) of 0.85 (**Table 1**). Analogously, among ORFs, the highest transitional substitution rate was observed in the Rep gene, which ranged from 11.34–15.59, and the highest transversional substitution rate was assessed in the REn gene, which ranged from 6.9–12.18, with the maximum transition/transversion bias (R) observed in the CP gene, which was 1.20. Furthermore, the transition/transversion bias (R) for the betasatellite was 0.85 (**Table 1**).

**Figure 4.**
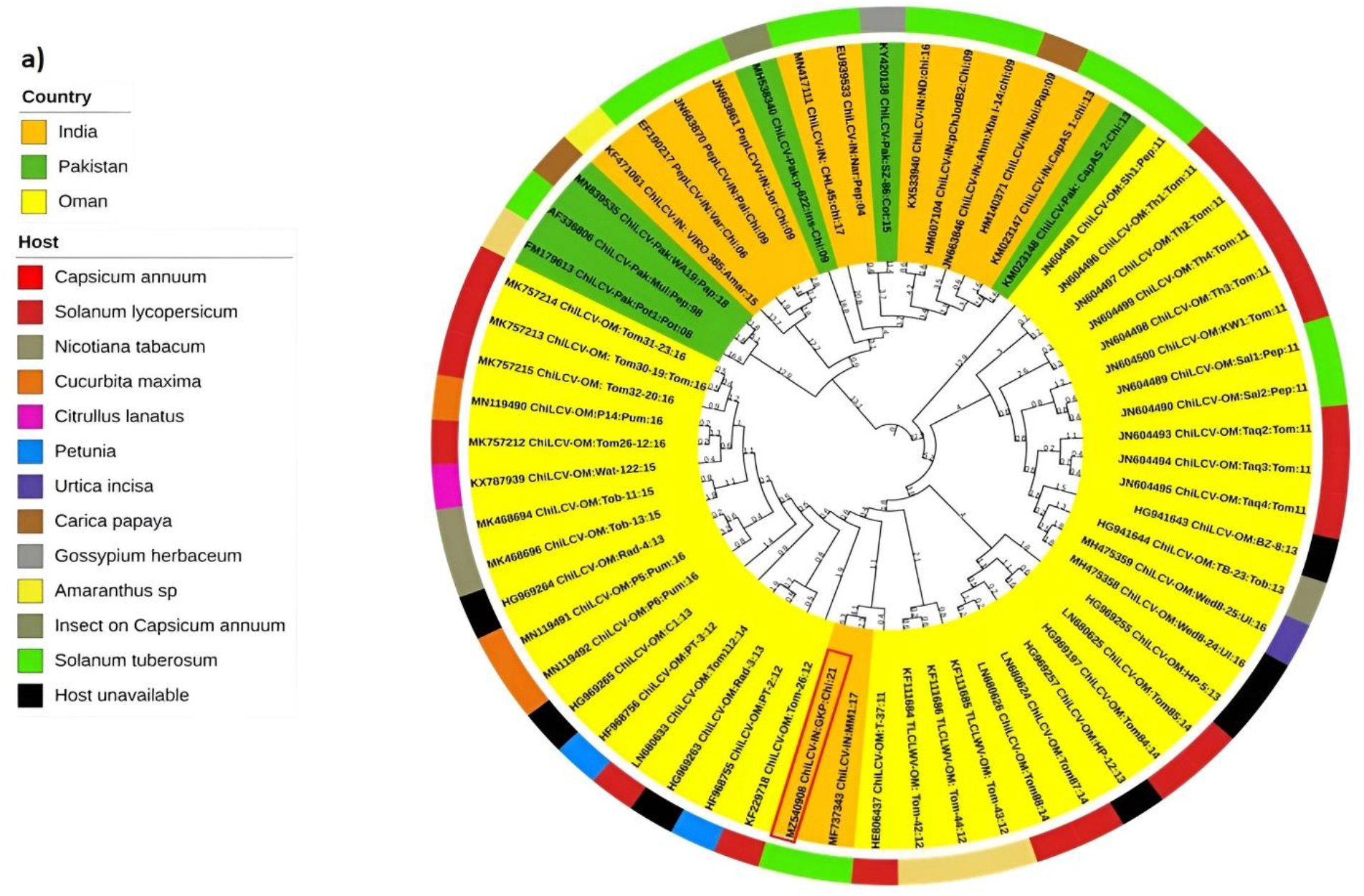
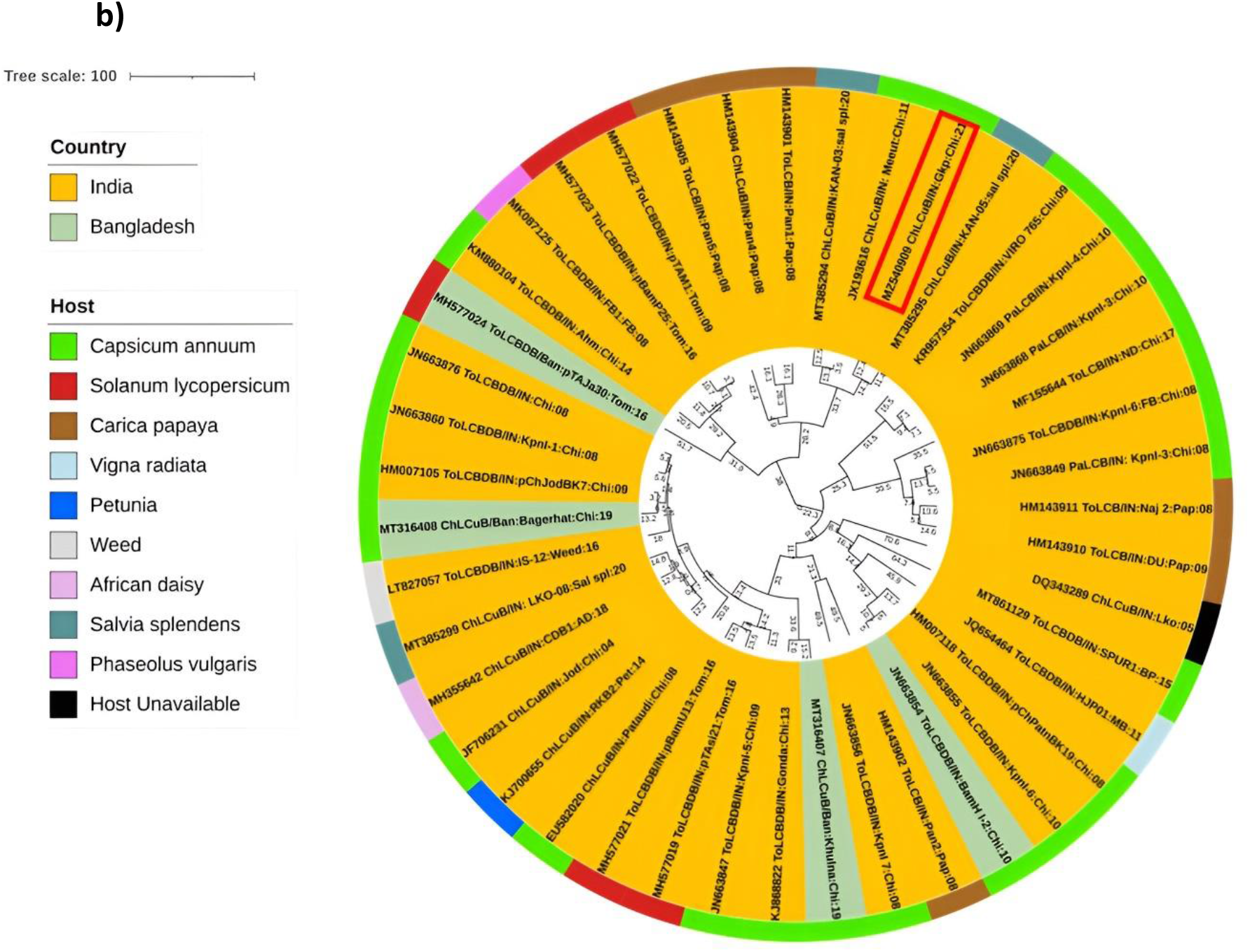
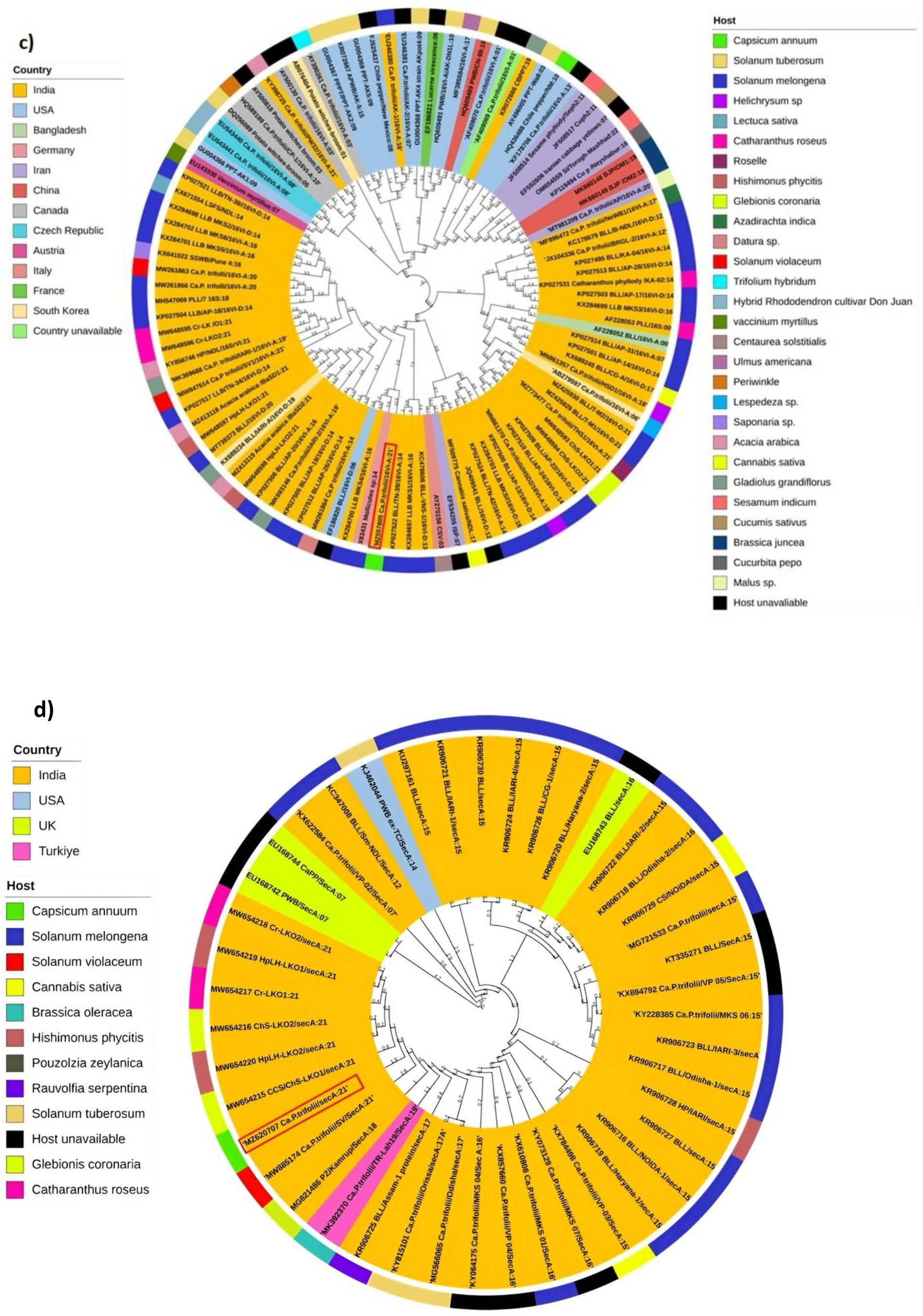
The Bayesian technique as well as the Tree annotator tool in the BEAST v.1.10 package (Suchard et al., 2018) were applied to Maximum clade credibility (MCC) phylogenetic evaluations derived from full-length nucleotide sequences of all identified begomovirus GKP isolates and Ca. P. trifolii 16S rRNA clone HSD2 and Sec A gene with a rooted tree mid-point and the tree was created using the web application Interactive Tree Of Life (iTOL) (for each begomovirus GKP isolate and Ca. P trifolii 16S rRNA clone HSD2 and Sec A gene, the numbers show the height median). The phylogenetic trees demonstrate that the viral populations and phytoplasma must therefore be examined separately: a) ChiLCV-A-GKP (Acc. No. MZ540908); b) ChLCuB-GKP (Acc. No. MZ540909); c) *Ca. P trifolii* 16S rRNA (Acc. No. MZ557805); and d) Ca. P trifolii Sec A gene (Acc. No. MZ620707). The outermost ring represents the host, while the innermost ring represents the country of origin.

**Table. 1.** The best fit model and transitional substitution rate of ChiLCV-A-GKP (Acc. No. MZ540908), six ORFs, and ChLCuB-GKP (Acc. No. MZ540909) with its βC1 gene of identified ChiLCV.

| Virus component | Best Model | Transitional Substitution Rate | Transversional Substitutions Rate | Transition/Transversion Bias (R) |
| --- | --- | --- | --- | --- |
| DNA-A | TN93+G | 10.58-15.95 | 5.04-7.29 | 0.96 |
| CP | T92+G+I | 10.41-16.67 | 4.79-6.97 | 1.20 |
| Pre-CP | JC+G | 9.96-16.6 | 5.43-6.41 | 0.86 |
| Rep | T92+G | 11.34-15.59 | 5.39-6.9 | 1.02 |
| TrAP | K2+G | 8.38-18.15 | 5.37-6.64 | 1.05 |
| REn | JC+G | 0.97-16.16 | 6.9-12.18 | 0.78 |
| C4 | K2+G | 10.13-17.95 | 5.13-7.05 | 1.03 |
| ChLCuB | GTR+G | 7.22-14.07 | 5.43-9.69 | 0.85 |
| βC1 | T92+G | 4.33-21.18 | 4.1-7.97 | 1.13 |

The same BEAST v.1.10 and the iTOL visualising software were used to analyse phytoplasma 16S rRNA and Sec-A gene datasets and observed that 16S rRNA shows clustering with Brinjal little leaf/TN-39/16VI-A: 14 (Acc. No. KP027522) reported from eggplant from India, whereas Sec-A with *Chrysanthemum coronarium* stunting/ChS-LKO1/secA:21 (Acc. No. MW654215) from India (**Figure 4c and 4d**). In addition to this, the transversional and transitional substitution rates for 16S rRNA were 2.58–4.31 and 9.68–26.1, respectively, with a bias (R) of 1.62 (Table 2). Whereas for the Sec-A gene, the bias (R) was 1.59 and the transversional and transitional substitution rates were 1.71-6.86 and 7.41-28.71, respectively (**Table 2**), and in comparison, to Sec A, 16S r RNA showed a significant high transitional/transversional bias.

**Table. 2.** The best fit model and transitional substitution rate *Ca. P. trifolii* 16S rRNA (Acc. No. MZ557805) and *Ca.P.trifolii* Sec A gene (Acc. No. MZ620707)

| Virus component | Best Model | Transitional Substitution Rate | Transversional Substitutions Rate | Transition/Transversion Bias (R) |
| --- | --- | --- | --- | --- |
| 16S rRNA | HKY+G+I | 9.68-26.1 | 2.58-4.31 | 1.62 |
| Sec-A | T92+G | 7.41-28.71 | 1.71-6.86 | 1.59 |

### Recombination estimation

Phylogenetic interweave networks containing reticulation were generated with the ChiLCV-A (**Figure 5a**) and ChLCuB **(Figure 5b)**, giving exceedingly clear evidence for recombination events by utilising the neighbour net method of Splits Tree v.4 (Huson et al., 2006) tool. Furthermore, for DNA-A with all of its ORFs and betasatellite, subsequent recombination analysis was achieved by the RDP v.4.2 package (Martin et al., 2015), which included identifying potential recombination breakpoints together with major and minor viral parents. This study discovered numerous and diverse recombination events, and a relevant recombination event supported by at least three approaches was chosen to reduce erroneous results. During this examination, twenty-one breakpoints were identified in DNA-A, with the highest incidence of breakpoints in the V1, C1, and C3 genes (**Table 3a**) (**Figure 6**). Secondly, recombination breakpoint analysis of 6 ORFs of DNA-A was used to validate our prior findings. CP was shown to contribute significantly to the most intra-species recombination (AV1) with five breakpoints. However, with only three breakpoints, the REP (AC1) and REn (AC3) genes may be minor recombination contributors (**Table 3b**). However, TrAP (AC2), AC4, and pre-CP (AV2) did not exhibit any recombination events. Concurrently, recombination breakpoints for datasets of ChLCuB were examined (**Figure 6**), and eleven putative breakpoints were identified among them, showing recombination evidence **(Table 3a)**, noting that very few recombination breakpoints in betasatellite around the βC1 ORF.

**Table. 3a.**
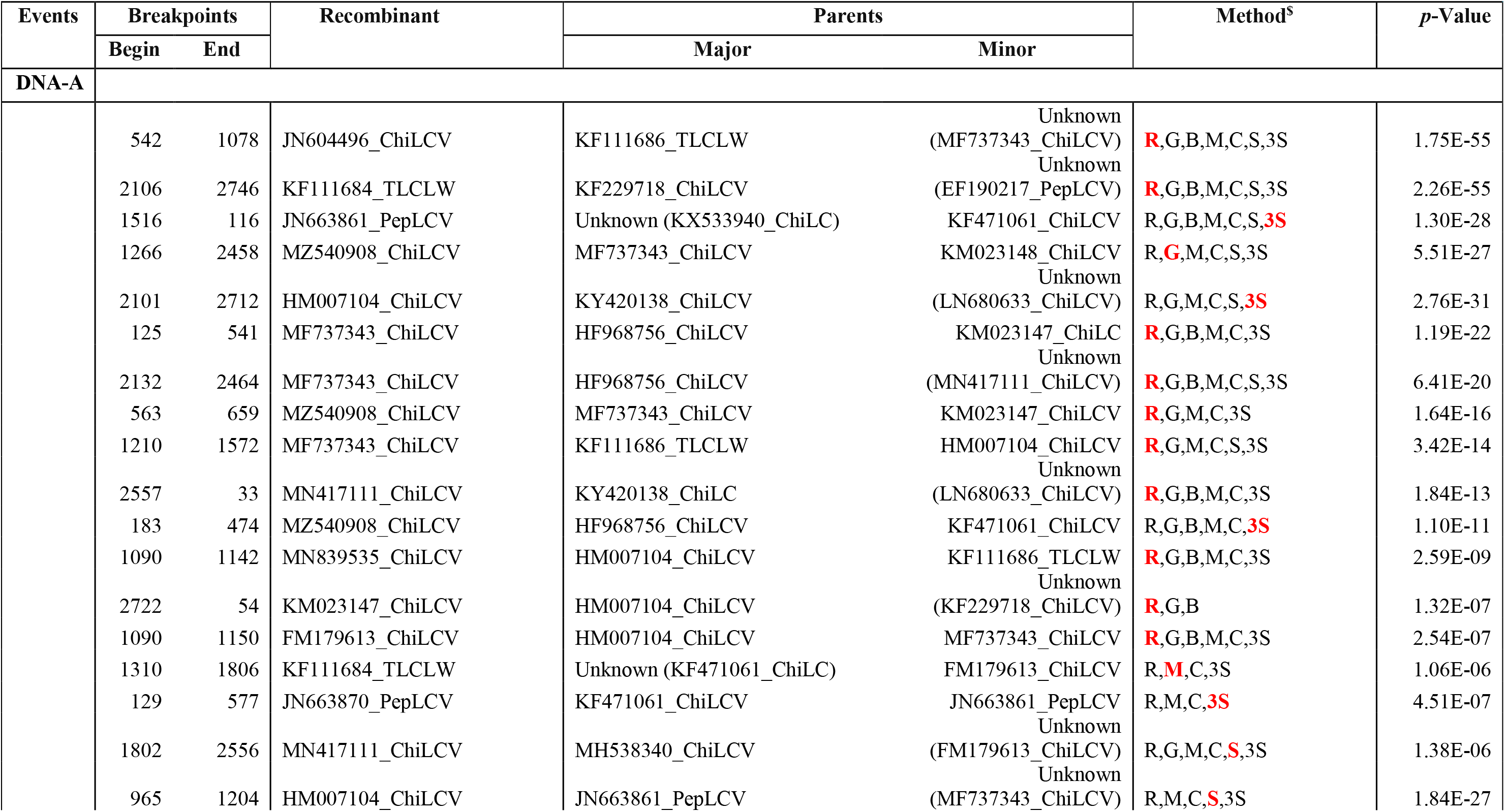
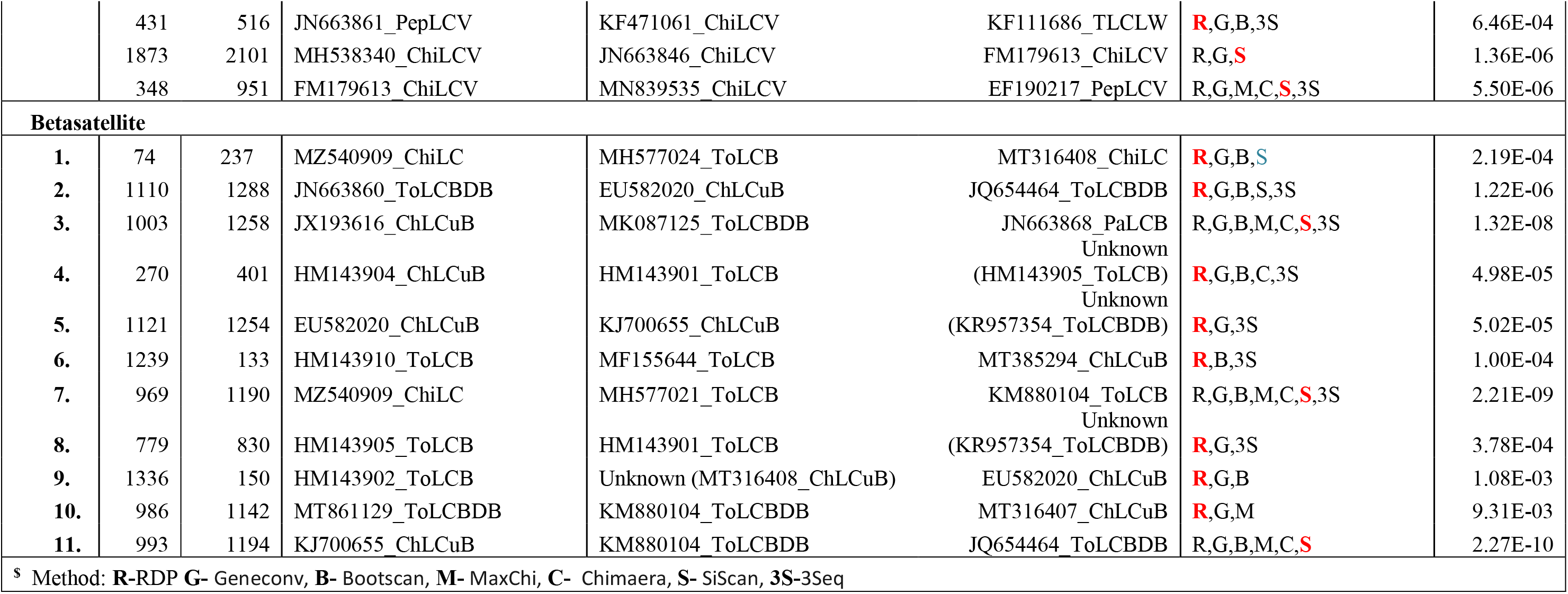
Recombination analysis for identified ChiLCV-A-GKP (Acc. No. MZ540908) and ChLCuB-GKP (Acc. No. MZ540909) isolates with major/minor parents and recombination events calculated by different algorithms.

**Table. 3b.** Putative breakpoints identification of all 6 ORFs of ChiLCV-A-GKP (Acc. No. MZ540908) isolates with major/minor parents and recombination events calculated by different algorithms.

| Events | | | Recombinant | Breakpoints | | Method<sup>\$</sup> | *p*-Value |
| --- | --- | --- | --- | --- | --- | --- | --- |
|  | Begin | End |  | Major | Minor |  |  |
| **Rep ( AC1)** |  |  |  |  |  |  |  |
| 1. | 544 | 1035 | KF111686_TLCLW | Unknown (MN839535_ChiLCV) | MF737343_ChiLCV | R,G,B,M,C,S,**3S** | 1.43E-24 |
| 2. | 808 | 234 | MZ540908_ChiLCV | MH538340_ChiLCV | JN604493_ChiLCV | **R**,G,B,M,C,3S | 3.25E-08 |
| 3. | 484 | 1070 | JN663870_PepLCV | JN604497_ChiLCV | KM023148_ChiLCV | R,G,B,M,C,**S**,3S | 3.34E-14 |
| **TrAP (AC2)** No Recombination Found |  |  |  |  |  |  |  |
| **REn (AC3)** |  |  |  |  |  |  |  |
| 1. | 275 | 405 | MF737343_ChiLCV | HM007104_ChiLCV | KF111686_TLCLW | G,B,**M**,3S | 2.52E-07 |
| 2. | 405 | 218 | MZ540908_ChiLCV | Unknown (JN663870_PepLC) | KM023147_ChiLCV | G,B,M,C,S,**3S** | 7.52E-07 |
| 3. | 348 | 71 | MN839535_ChiLCV | HM007104_ChiLCV | Unknown (JN663870_PepLCV) | B,M,**S** | 8.44E-07 |
| **C4** No Recombination Found |  |  |  |  |  |  |  |
| **CP (AV1)** |  |  |  |  |  |  |  |
| 1. | 296 | 393 | MZ540908_ChiLCV | MF737343_ChiLCV | MN417111_ChiLCV | R,**G**,B,M,C,3S | 2.16E-14 |
| 2. | 749 | 202 | MF737343_ChiLCV | LN680633_ChiLCV | Unknown (JN604491_ChiLCV) | R,**G**,B,M,C,S,3S | 6.25E-07 |
| 3. | 686 | 202 | JN604491_ChiLCV | KM023147_ChiLCV | LN680633_ChiLCV | R,M,**C**,3S | 7.20E-05 |
| 4. | 653 | 144 | KF471061_ChiLCV | JN604489_ChiLCV | Unknown (LN680633_ChiLCV) | **R**,G,M,C,S,3S | 6.69E-09 |
| 5. | 722 | 216 | MH538340_ChiLCV | JN604489_ChiLCV | Unknown (HG969263_ChiLCV) | **R**,G,M,C,3S | 3.18E-07 |
| **Pre-CP ( AV2)** No Recombination Found |  |  |  |  |  |  |  |
| <sup>\$</sup> Method : **R**-RDP **G**- Geneconv, **B**- Bootscan, **M**- MaxChi, **C**- Chimaera, **S**- SiScan, **3S**-3Seq | | | | | | | |

**Figure 5.**
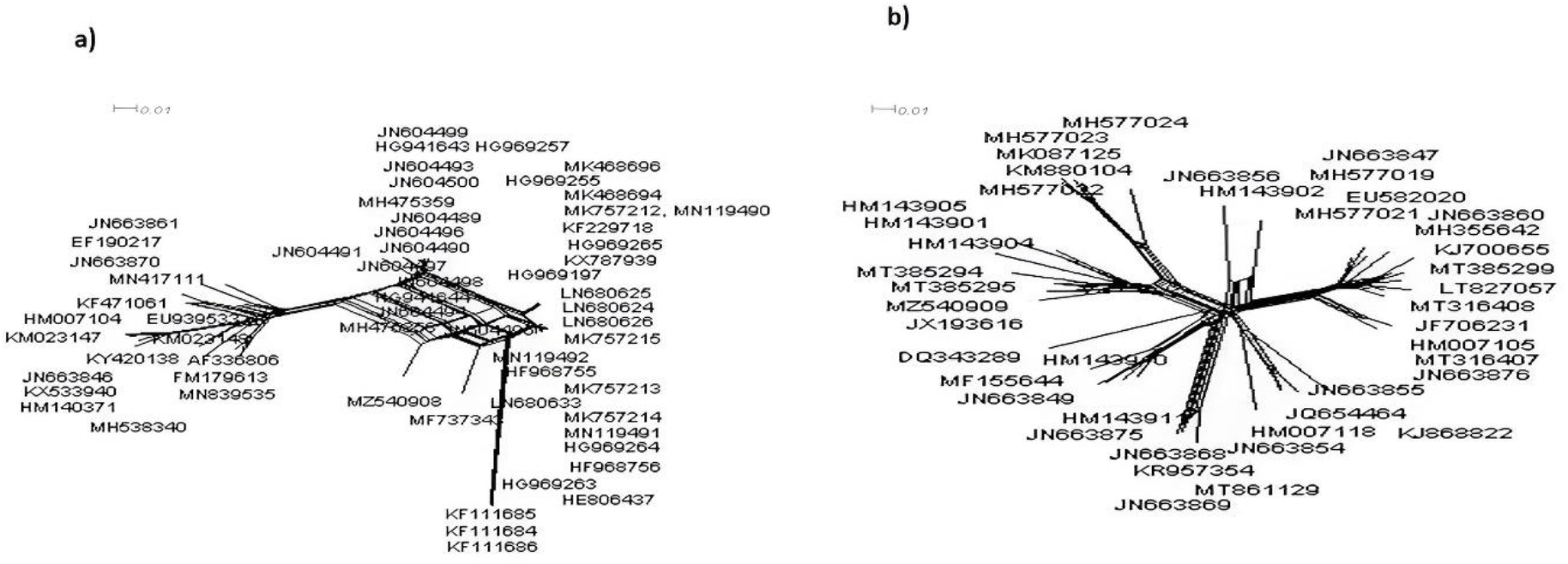
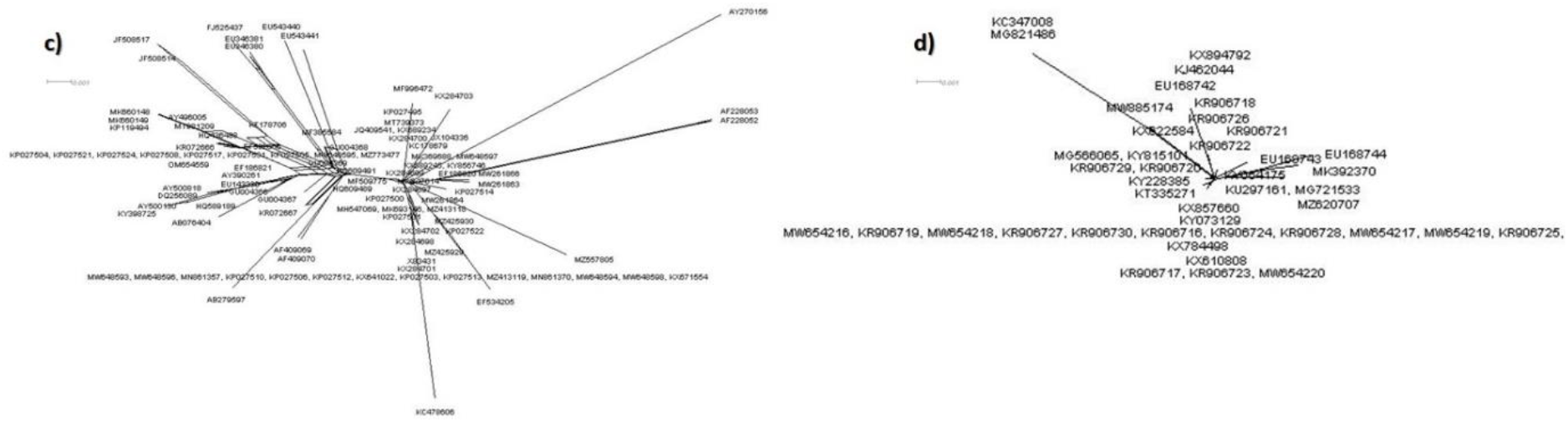
Phylogenetic dendrogram analysis with neighbour-net based on the alignment of full-length nucleotide sequences (neighbour net method of Splits Tree v.4 (Huson et al., 2006) software) and the presence of recombination events was reflected by reticulated networks; a) ChiLCV-A-GKP (Acc. No. MZ540908); b) ChLCuB-GKP (Acc. No. MZ540909); c) Ca. P. trifolii 16S rRNA (Acc. No. MZ557805); and d) Ca.P.trifolii Sec A gene (Acc. No. MZ620707).

**Figure 6.**
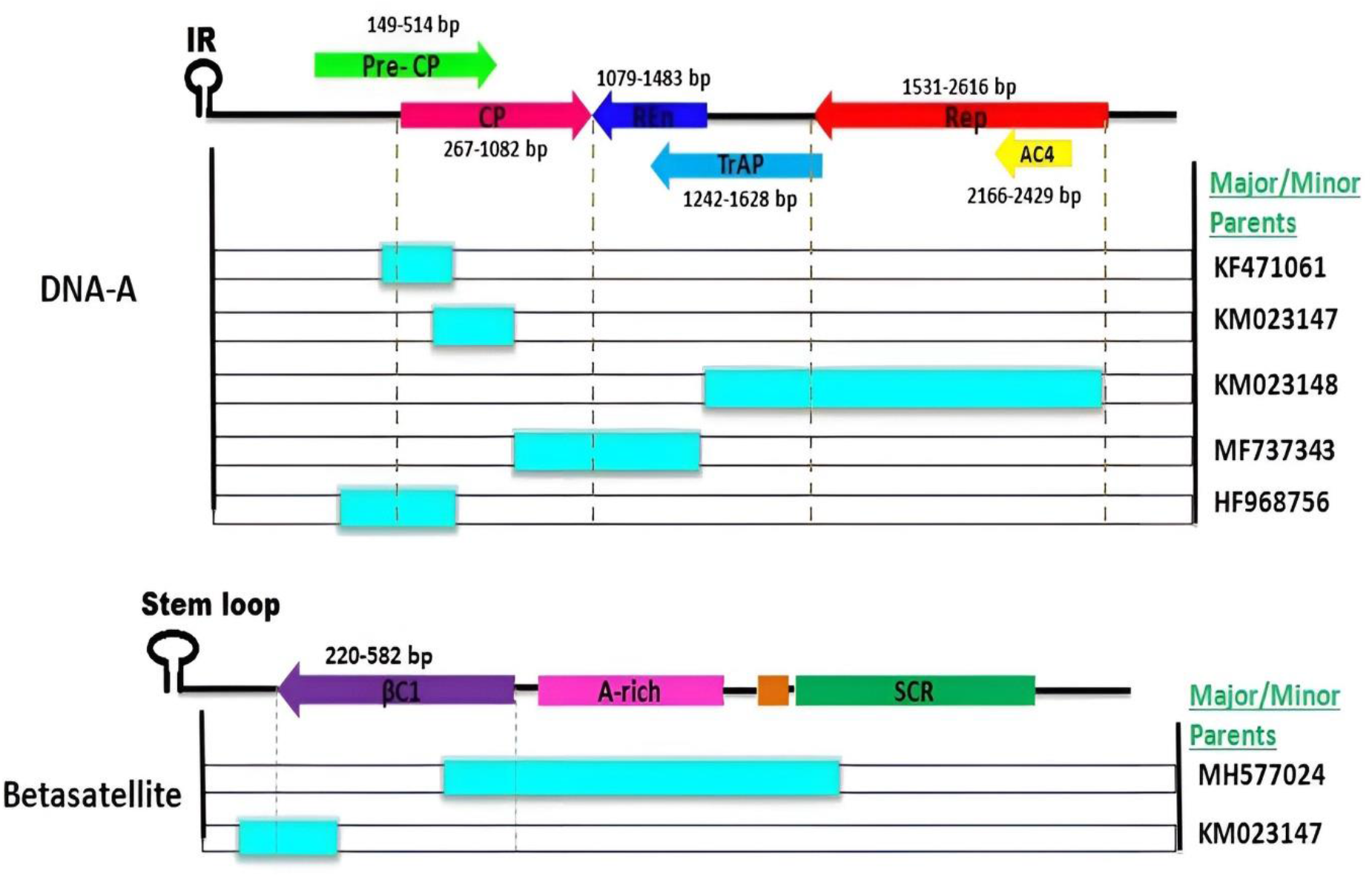
Recombination events have been observed in the ChiLCV-A-GKP (Acc. No. MZ540908) and ChLCuB-GKP (Acc. No. MZ540909) isolates; the genome layout at the top of the ORFs graph matches to the sequences’ schematic representations above, with each ORF, coloured differently. The blue sections relate to the donated (major/minor parent) parts (see Table 3a and 3b for further information on recombination occurrences)

Simultaneously, to support our previous reticulated results for 16S rRNA (**Figure 5c**) and Sec A (**Figure 5d**), we performed recombination analysis and found just two recombinant breakpoints in 16S rRNA datasets but none in Sec A datasets. This implies recombination was much less common among them (**Table 4**).

**Table. 4.** Recombination analysis for identified *Ca. P. trifolii* 16S rRNA (Acc. No. MZ557805) and *Ca.P.trifolii* Sec A gene (Acc. No. MZ620707). calculated by different algorithms.

| Events | Breakpoints | | Recombinant | Parents | | Method<sup>\$</sup> | *p*-Value |
| --- | --- | --- | --- | --- | --- | --- | --- |
|  | Begin | End |  | Major | Minor |  |  |
| 16S rRNA |  |  |  |  |  |  |  |
| 1. | 1676 | 1767 | AB279597_Ca.P. | KP119494_Cucur | Unknown (AY270156_Centa) | R,G,M,C,S,**3S** | 1.06E-09 |
| 2. | 27 | 1625 | AF228053_Periw | Unknown (KP119494_Cucur) | KX284703_Brinj | R,G,M,C,**S**,3S | 6.31E-09 |
| Sec-A No Recombination Found |  |  |  |  |  |  |  |
| \$ Method: **R**-RDP **G**- Geneconv, **B**- Bootscan, **M**- MaxChi, **C**- Chimaera, **S**- SiScan, **3S**-3Seq | | | | | | | |

### Coalescent analysis

Recombination and nucleotide substitution both have an impact on the genetic diversity and evolution of begomovirus. To validate this statement, we calculate the mean substitution rate with a 95% HPD interval at both relaxed and strict clock for all ChiLCV components (DNA-A, all 6 ORFs and betasatellite) by BEAST v.1.10. Furthermore, a relaxed molecular clock was identified to explain the significant positive selection within mean nucleotide substitution (Duffy and Holmes, 2009) (**Table 5**). The average rate of nucleotide substitution in ChiLCV-A was 2.445×10^−3^ substitutions/site/year with a 95% HPD interval of 6.23×10^−4^ to 5.15×10^−3^, while for betasatellite we observed a substitution rate of 7.27×10^−4^ substitutions/site/year with a 95% HPD interval of 3.86×10^−4^ to 1.07×10^−3^, indicating rapid evolution as it falls within the range of RNA virus nucleotide substitution rate (**Table 5**). Consequently, the overall nucleotide substitution rate for ChLCuB was 9.62×10-4 substitutions/site/year (within the 95% HPD interval of 5.64×10-5 to 2.28×10-3). The substitution rate for all ORFs of DNA-A was also determined to be in the high range of 7.15×10-4 substitutions/site/year for the Rep gene, this could be related to the rapid mutation rate **(Table 5)**. Although this rate of mutation amongst plant viruses is crucial in genetic variation and evolution, it was computed for every component of ChiLCV at three locations of codon and observed significant high-rate mutation at wobble codon positions 3 for DNA-A and position 2 for ChLCuB. The rate of mutation in ORFs was highest at codon position 3 for the CP gene and secondly at codon position 1 for the pre-CP gene (**Table 5**). Interestingly, the mutation rate for all genes except pre-CP was highest at codon position 3.

**Table. 5.** Estimation of mean substitution rate (at 95% HPD) and codon position mutation rate (CP1, CP2, CP3) for ChiLCV-A-GKP (Acc. No. MZ540908), six ORFs, and ChLCuB-GKP (Acc. No. MZ540909) with discovered ChiLCV C1 gene

| Viral component | Mean substitution rate<br>(at 95% HPD interval) |  | Mutation at various codon positions<br>(CP1, CP2, CP3 respectively) |  |
| --- | --- | --- | --- | --- |
|  | Relaxed clock | Strict clock | Relaxed clock | Strict clock |
| **DNA-A** | 2.445E-3 [6.2326E-4, 5.1519E-3] | 3.772E-4 [2.2925E-4, 5.254E-4] | 1.127,0.672,1.201 | 1.0924,0.679,1.227 |
| **CP** | 8.5067E-5 [1.3442E-5, 1.7361E-4] | 1.3397E-5 [6.6824E-6, 2.0255E-5] | 0.341,0.439, 2.19 | 0.376,0.401,2.223 |
| **Pre-CP** | 6.0558E-5 [9.8569E-9, 1.5974E-4] | 8.3002E-8 [5.0137E-21, 4.9458E-7] | 1.468,0.333,1.199 | 1.446,0.333,1.221 |
| **Rep** | 7.1531E-4 [1.7734E-6, 1.76E-3] | 6.152E-5 [1.2417E-11, 2.0294E-4] | 0.762,0.459,1.779 | 0.710,0.444,1.846 |
| **TrAP** | 4.6471E-4 [2.3552E-4, 7.1808E-4] | 4.077E-4 [2.1793E-4, 6.1867E-4] | 0.788,0.803,1.1412 | 0.787,0.808,1.408 |
| **REn** | 3.4299E-4 [9.5192E-5, 6.1357E-4] | 4.1119E-4 [1.5651E-4, 6.9087E-4] | 1.079,0.745,1.176 | 1.076,0.741,1.183 |
| **C4** | 4.7956E-4 [7.2879E-6, 1.1285E-3] | 1.0374E-4 [8.3516E-9, 2.4964E-4] | 0.719,0.538,0.977 | 0.554,1.477,0.969 |
| **Betasatellite** | 7.2679E-4 [3.8551E-4, 1.069E-3] | 6.2016E-4 [3.8937E-4, 8.4136E-4] | 0.921,1.155,0.924 | 0.921,1.157,0.922 |
| **βC1** | 7.0983E-4 [1.7269E-4, 1.2736E-3] | 6.6045E-4 [2.6013E-4, 1.1159E-3] | 0.769,0.567,1.664 | 0.758,0.581,1.661 |

The mean substitution rate and mutation rate of phytoplasma 16S rRNA and Sec-A genes were also analysed using the same method as for ChiLCV components. The average nucleotide substitution rate for 16S rRNA was 1.64×10^−4^ substitutions/site/year, with a 95% HPD interval of 6.21×10^−5^ to 2.61×10^−4^, while Sec-A had a substitution rate of 7.02×10^−4^ substitutions/site/year (with a 95% HPD interval of 2.42×10^−3^ to 1.14×10^−2^) (**Table 6**). Surprisingly, we observed that the mean substitution rate for both the 16S rRNA and Sec-A genes was remarkably comparable to that of an RNA virus (10^−2^-10^−5^ nucleotide substitution per site, each year) (Duffy et al., 2008; Pagán and Garca-Arenal, 2018). Whereas both have quite a high rate of mutation at codon position 1, with 1.119 and 1.566, which was similar to ChiLCV (**Table 6**).

**Table. 6.**
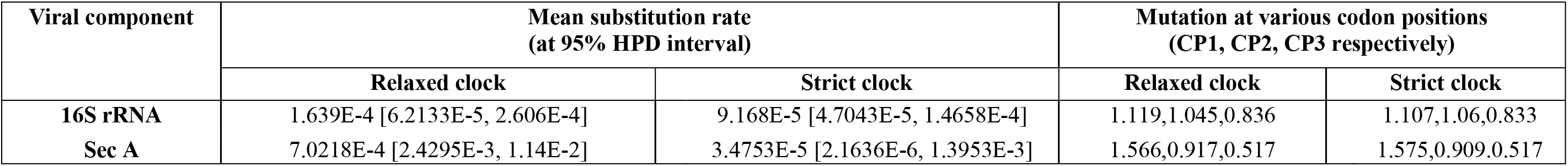
Estimation of mean substitution rate (at 95% HPD) and codon position mutation rate (CP1, CP2, CP3) for identified *Ca. P. trifolii* 16S rRNA (Acc. No. MZ557805); and *Ca.P.trifolii* Sec A gene (Acc. No. MZ620707).

### Population Demographic estimation

To strengthen the evaluation of chilli DNA-A and betasatellite isolates, genetic variability was analyzed using DnaSP v. 6.0. (Rozas et al., 2017). However, the number of polymorphic sites (s) for ChiLCV-A and ChLCuB was 911 and 554, respectively, while the total number of mutations (η) for DNA-A was 1092 and 746 for betasatellites (**Table 7**). Likewise, nucleotide diversity (π) for DNA-A and associated betsatellite was 0.07792 and 0.08597, respectively. Furthermore, to verify the dispersion of anomalous mutations in the viral genome, we assessed the nucleotide diversity (π) for every ORF on DNA-A, viz., CP, pre-CP, Ren, Rep, Trap, and C4 (>0.08) genetic variability that was especially high in CP and Rep, which was 0.13235 and 0.06850 respectively, demonstrating that they contributed majorly to DNA-A genetic diversification. The estimated value of nucleotide differences (k) for DNA-A and betasatellites sequences was 209.91 and 113.31, respectively (**Table 7**). Overall, the diversity of ChiLCV-A sequences was greater than that of ChLCuB sequences as per the outcomes.

**Table. 7.**
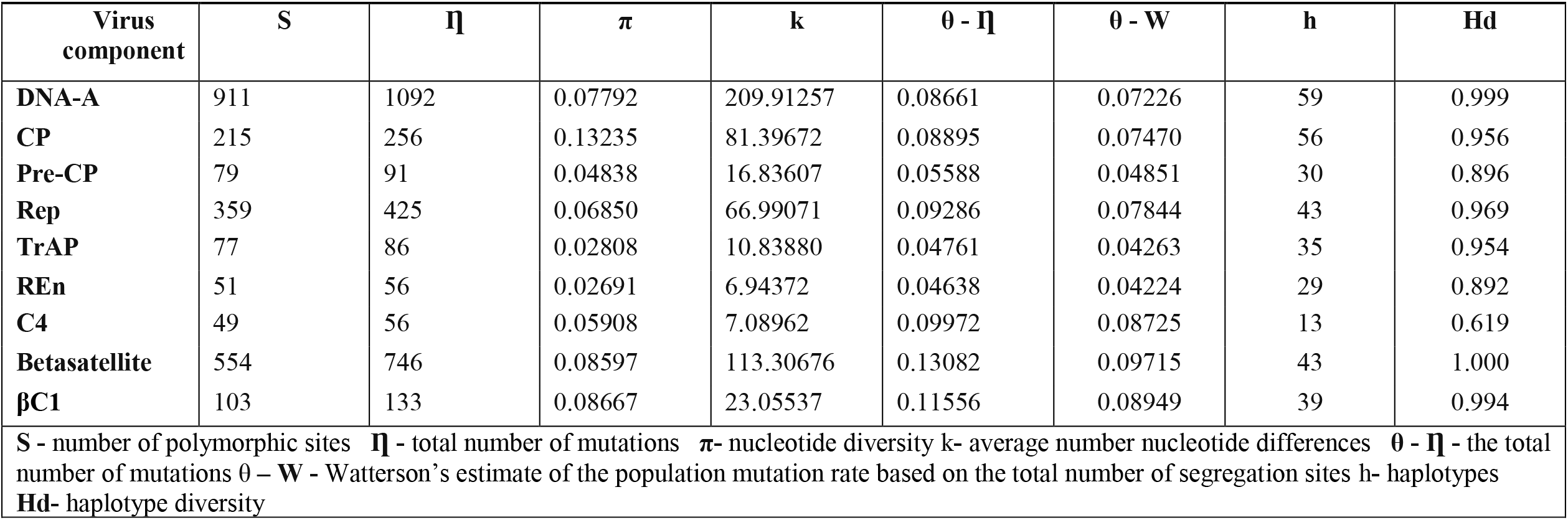
Population demographic estimation of ChiLCV-A-GKP (Acc. No. MZ540908), six ORFs, and ChLCuB-GKP (Acc. No. MZ540909) with βC1 gene of identified ChiLCV.

| Virus component | S | $\Pi$ | $\pi$ | k | $\theta - \Pi$ | $\theta - W$ | h | Hd |
| --- | --- | --- | --- | --- | --- | --- | --- | --- |
| DNA-A | 911 | 1092 | 0.07792 | 209.91257 | 0.08661 | 0.07226 | 59 | 0.999 |
| CP | 215 | 256 | 0.13235 | 81.39672 | 0.08895 | 0.07470 | 56 | 0.956 |
| Pre-CP | 79 | 91 | 0.04838 | 16.83607 | 0.05588 | 0.04851 | 30 | 0.896 |
| Rep | 359 | 425 | 0.06850 | 66.99071 | 0.09286 | 0.07844 | 43 | 0.969 |
| TrAP | 77 | 86 | 0.02808 | 10.83880 | 0.04761 | 0.04263 | 35 | 0.954 |
| REn | 51 | 56 | 0.02691 | 6.94372 | 0.04638 | 0.04224 | 29 | 0.892 |
| C4 | 49 | 56 | 0.05908 | 7.08962 | 0.09972 | 0.08725 | 13 | 0.619 |
| Betasatellite | 554 | 746 | 0.08597 | 113.30676 | 0.13082 | 0.09715 | 43 | 1.000 |
| βC1 | 103 | 133 | 0.08667 | 23.05537 | 0.11556 | 0.08949 | 39 | 0.994 |
| **S** - number of polymorphic sites **$\Pi$** - total number of mutations **π**- nucleotide diversity k- average number nucleotide differences **$\theta - \Pi$** - the total number of mutations $\theta - W$ - Watterson's estimate of the population mutation rate based on the total number of segregation sites h- haplotypes<br>**Hd**- haplotype diversity | | | | | | | | |

Furthermore, the average of nucleotide differences across ChiLCV-A coding areas was computed, and it was found that the gene AV1/CP had the best k value of 81.40 **(Table 7).** Furthermore, 59 haplotypes for DNA-A were found in addition to its ORFs, with the highest haplotypes for the CP (56nt) and Rep (43 nt) genes **(Figure 7)**, but only 43 and 39 for betasatellite and βC1 genes, respectively. The haplotype diversity (Hd) of isolates with ORFs of DNA-A and betasatellite that were nearly identical or close to one another was assessed. According to this study, the viruses we isolated were diverse (**Table 7**), and among ORFs, the CP and AC4 genes had the maximum levels of genetic variation (**Figure 7**).

**Figure 7.**
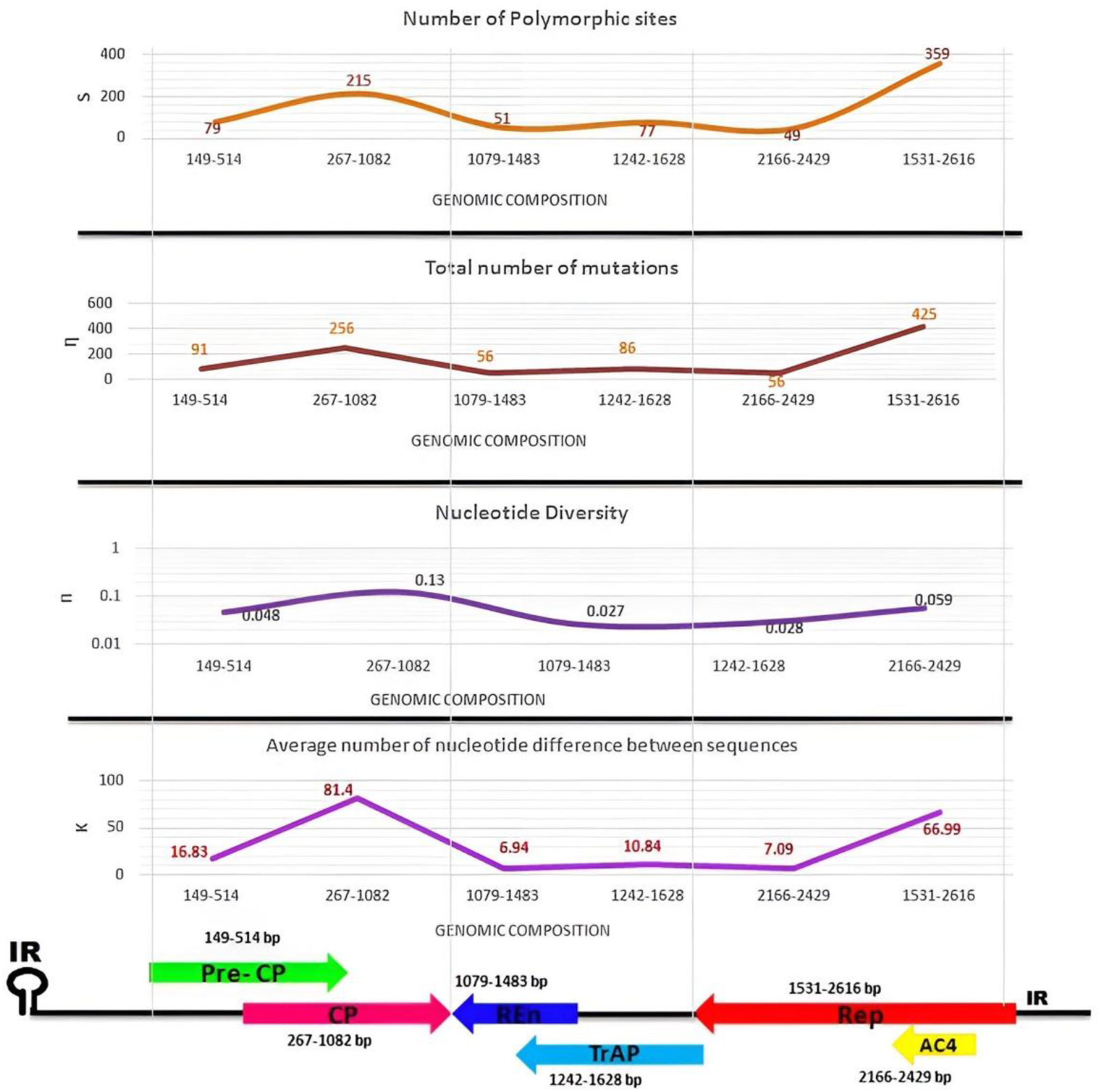
The genetic diversity of ChiLCV-A-GKP (Acc. No. MZ540908) was calculated using DnaSP v. 6.0 (Rozas et al., 2017)., and the major factor involved in variability (the number of polymorphic sites (S), total number of mutations (η), nucleotide diversity (π), and the average of nucleotide differences (k)) was graphically compared between all 6 ORFs of DNA-A-GKP, we noticed the involvement of CP and AC4 REP genes.

Neutrality tests, i.e., Tajima’s D, Fu & Li’s D, and Fu & Li’s F were also performed to analyse the several forms of selecting acts on ChiLCV-A, its coding regions, and the betasatellites. Significant statistical discrepancies were identified using the neutrality test across each component based on the population under consideration (**Table S10**). Conversely, the maximal datasets had highly negative statistical values (**Table S10**), indicating that the variability at the genetic level in the population was quite significant and specific to the sequences themselves. Secondly, we noticed that these nucleotide diversity values were obtained when the population’s evident variability was not continuous and swiftly vanished due to purifying selection either through nucleotide-level activities or population proliferation at the nucleotide level. Furthermore, the negative results (**Figure 8)** and (**Table S10**) of all neutrality tests indicate that our ChiLCV isolate and its associated betasatellite were under purifying selection, implying that the gene’s nature seems conserved

**Figure 8.**
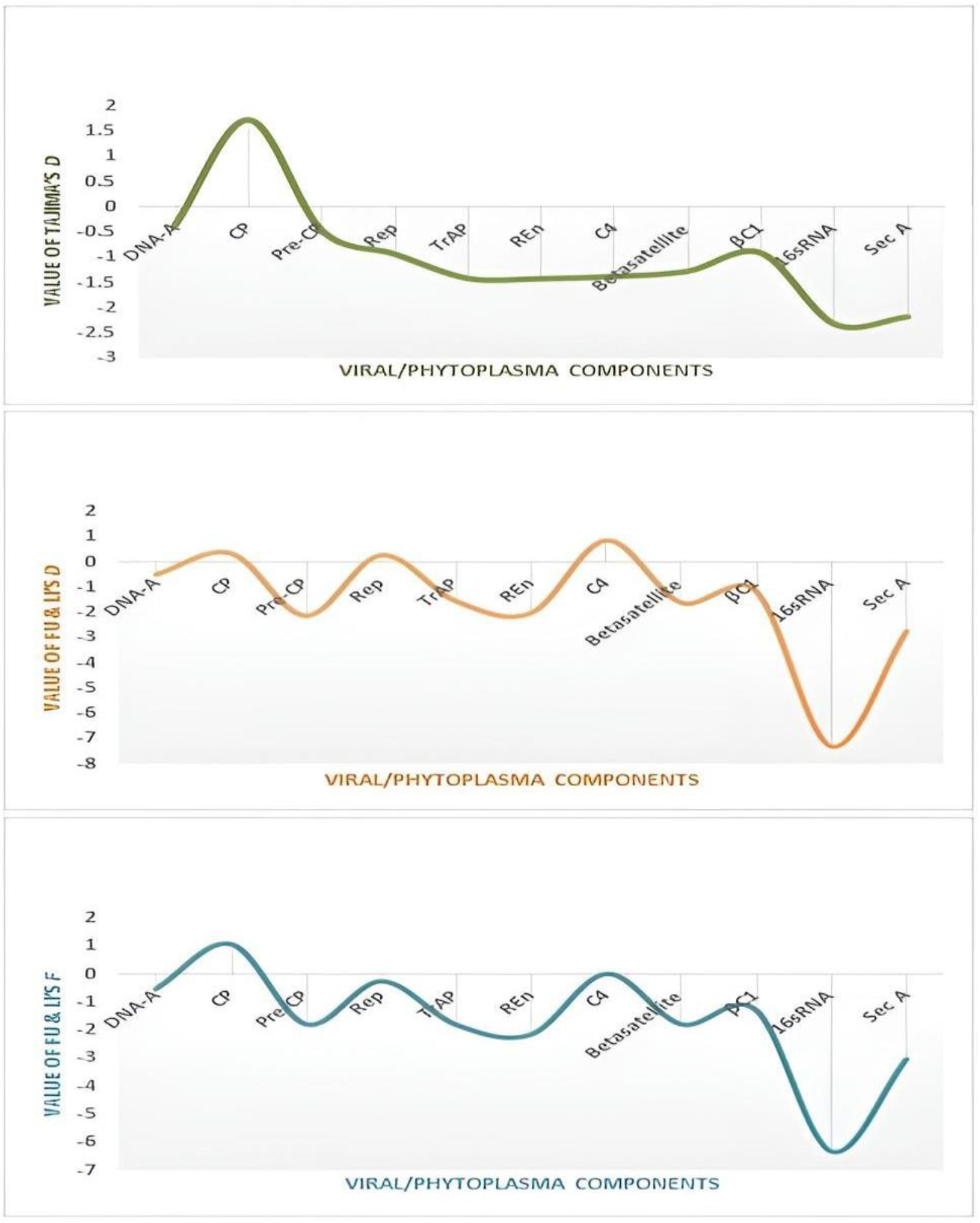
Graphical comparison of all three neutrality tests for ChiLCV-A-GKP (Acc. No. MZ540908) along with its six ORFs; ChLCuB-GKP (Acc. No. MZ540909); *Ca. P. trifolii*16S rRNA (Acc. No. MZ557805); and *Ca. P. trifolii* Sec A gene (Acc. No. MZ620707); while examined to begomoviral and phytoplasma components, both 16S rRNA and Sec A were showing extremely negative value indicates its population under purifying selection

Population structural analysis was executed for phytoplasma components 16S rRNA and Sec-A and found that our sequences performed adequately. For instance, the number of polymorphic sites (s) reported more for 16S rRNA was 62, whereas the nucleotide diversity (π) found for the Sec-A gene was 0.00352 (**Table 8**). Meanwhile, we observed that 16S rRNA was the dominating one in terms of the total number of mutations (η), nucleotide differences (k), haplotypes (h), and haplotype diversity (Hd) with a score of 63, 3.41, 30, and 0.760 respectively. However, the neutrality test by 16S rRNA and Sec-A yielded an unexpectedly negative result and signifies its population in the purifying selection while comparing with outcomes of ChiLCV (**Figure 8**) (**Table S11**).

**Table 8.**
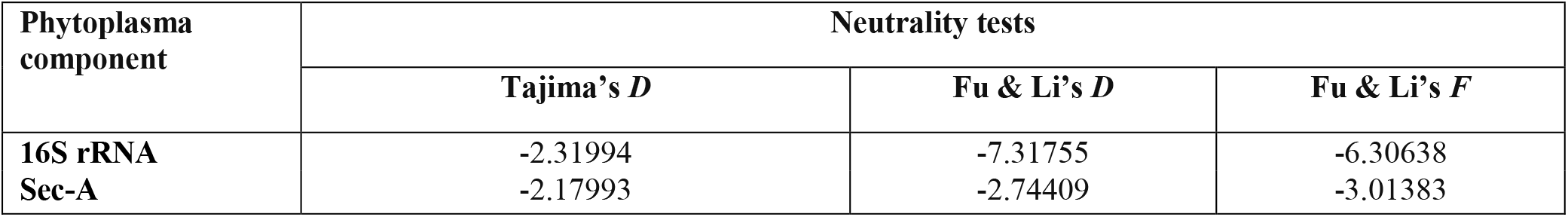
Results of different neutrality tests analysis *Ca.P. trifolii* 16s rRNA (Acc. No. MZ557805) and *Ca.P.trifolii* Sec A gene (Acc. No. MZ620707).

### Selection analysis

The dN/dS were evaluated for every gene as well as the genomic sequence to gain a better understanding of the impact of selection pressure upon the non-uniform distribution of variation observed between DNA-A, all ORFs, and betasatellite. Except for REn and AC4, practically all genes had dN/dS ratios <1, revealing that negative selection was operating in both regions **(Table 9**). The dN/dS ratio of the REn and AC4 genes was >1, with values of 1.48 and 1.06, respectively, while it was 0.49 for DNA-A and ranged from 0.67 to 0.93 for other ORFs (CP, pre-CP, Rep, and TrAP). This shows that each site in the genome was subjected to a varied level of selection pressure (**Table 9**) (**Figure 9a**). Despite this, betasatellite was under positive selection with dN/dS >1, i.e., 1.21 (**Table 9**) (**Figure 9b**). In keeping with the high dN/dS ratio of the REn and AC4 genes (**Table 9**), we obtained proof of positive selection by FEL (p≤0.1) and FUBER. FEL detected 2 and 4 positive sites, while FUBER detected 3 and 6 positively selected sites for REn and AC4 respectively (**Table 10**). Additionally, we identified 15, 36, and 30 positively selected sites for ChLCuB identified by SLAC, FEL, and FUBER, respectively (**Table 10**).

**Table 9.**
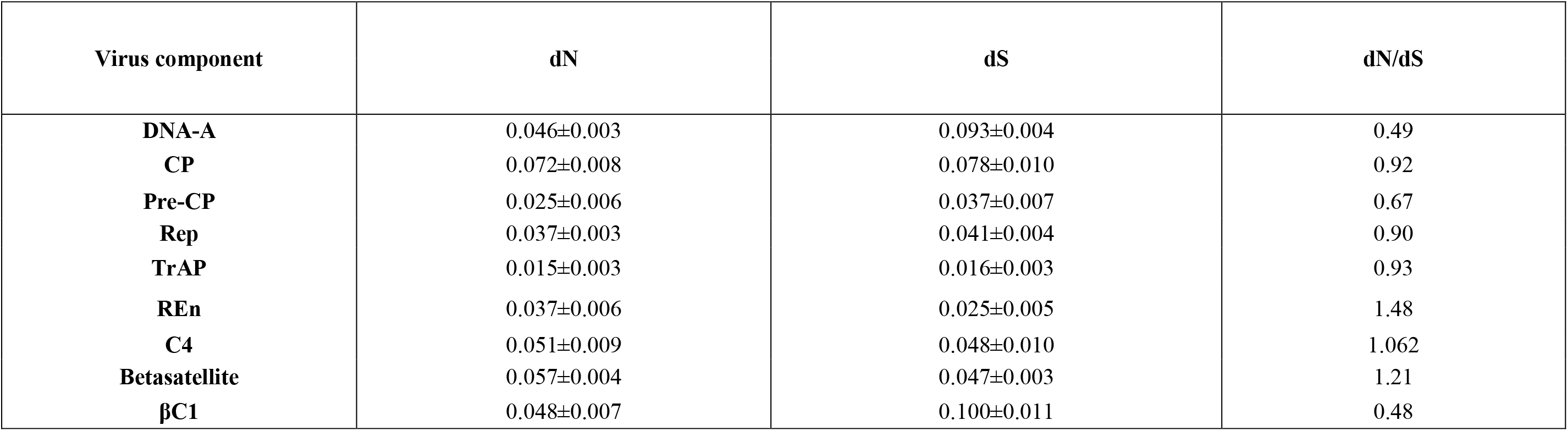
Selection pressure analysis for ChiLCV-A-GKP (Acc. No. MZ540908), six ORFs, and ChLCuB-GKP (Acc. No. MZ540909) with βC1 gene of identified ChiLCV.

| Virus component | dN | dS | dN/dS |
| --- | --- | --- | --- |
| **DNA-A** | 0.046±0.003 | 0.093±0.004 | 0.49 |
| **CP** | 0.072±0.008 | 0.078±0.010 | 0.92 |
| **Pre-CP** | 0.025±0.006 | 0.037±0.007 | 0.67 |
| **Rep** | 0.037±0.003 | 0.041±0.004 | 0.90 |
| **TrAP** | 0.015±0.003 | 0.016±0.003 | 0.93 |
| **REn** | 0.037±0.006 | 0.025±0.005 | 1.48 |
| **C4** | 0.051±0.009 | 0.048±0.010 | 1.062 |
| **Betasatellite** | 0.057±0.004 | 0.047±0.003 | 1.21 |
| **βC1** | 0.048±0.007 | 0.100±0.011 | 0.48 |

**Figure 9.**
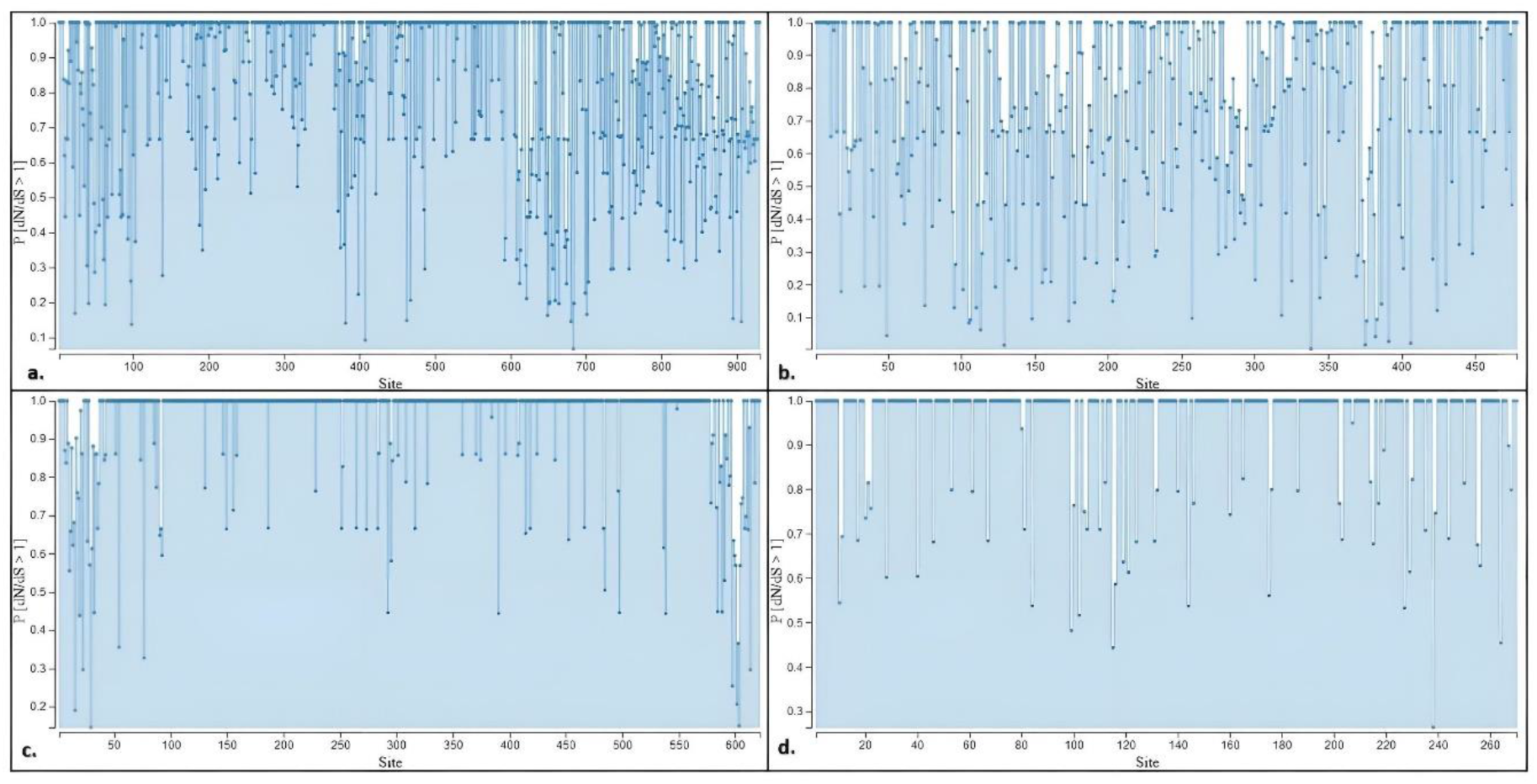
Graph generated by DataMonkey (www.datamonkey.org) (Weaver et al., 2018) showing the P[dN/dS>1] ratio for all coding sites of a) ChiLCV-A-GKP (Acc. No. MZ540908) with >900 amino acid site; b) ChLCuB-GKP (Acc. No. MZ540909) with >450 amino acid site; c) *Ca. P.trifolii16S* rRNA (Acc. No. MZ557805) >600 amino acid site; and d) *Ca.P.trifolii* Sec A gene (Acc. No. MZ620707) > 260 amino acid site. The maximum site ratio was dN/dS<1, indicating negative/purifying selection, while a few sites with dN/dS>1 imply positive/diversifying selection

**Table 10.** The number of negatively and positively selected sites for ChiLCV-A-GKP (Acc. No. MZ540908), six ORFs, and ChLCuB-GKP (Acc. No. MZ540909) with βC1 gene of identified ChiLCV.

| Virus Components | Negative/Purifying Selection |  |  |  | Positive/Diversifying Selection |  |  |  |
| --- | --- | --- | --- | --- | --- | --- | --- | --- |
|  | SLAC | FEL p≤0.1 | FUBER | Total | SLAC | FEL p≤0.1 | FUBER | Total |
| **DNA-A** | 77 | 141 | 5 | 141 | 2 | 27 | 87 | 87 |
| **CP** | 44 | 88 | 78 | 88 | 0 | 0 | 1 | 1 |
| **Pre CP** | 7 | 15 | 10 | 15 | 0 | 0 | 5 | 5 |
| **REP** | 40 | 103 | 99 | 103 | 1 | 2 | 4 | 4 |
| **TRAP** | 7 | 13 | 11 | 13 | 0 | 1 | 3 | 3 |
| **REN** | 9 | 13 | 9 | 13 | 0 | 2 | 3 | 3 |
| **C4** | 2 | 4 | 2 | 4 | 0 | 4 | 6 | 6 |
| **Betasatellite** | 18 | 32 | 11 | 32 | 15 | 36 | 30 | 36 |
| **βC1** | 23 | 34 | 26 | 34 | 0 | 1 | 0 | 1 |

Furthermore, both phytoplasma 16S rRNA and Sec-A genes were under purifying selection; otherwise, we detected a few sites under diversifying selection via FEL and FUBAR, but no significant ratio for dN/dS >1 (**Table 11 and Table 12**) (**Figure 9c and 9d**).

**Table 11.** Selection pressure analysis for identified *Ca.P. trifolii* 16s rRNA (Acc. No. MZ557805) and *Ca.P.trifolii* Sec A gene (Acc. No. MZ620707).

| Phytoplasma Components | dN | dS | dN/dS |
| --- | --- | --- | --- |
| 16S rRNA | 0.001±0.000 | 0.004±0.001 | 0.25 |
| SEC A | 0.004±0.001 | 0.007±0.001 | 0.57 |

**Table 12.** The number of negatively and positively selected sites for identified *Ca.P. trifolii* 16s rRNA (Acc. No. MZ557805) and *Ca.P.trifolii* Sec A gene (Acc. No. MZ620707).

| Phytoplasma Components | Negative/Purifying Selection |  |  |  | Positive/Diversifying Selection |  |  |  |
| --- | --- | --- | --- | --- | --- | --- | --- | --- |
|  | SLAC | FEL p≤0.1 | FUBER | Total | SLAC | FEL p≤0.1 | FUBER | Total |
| 16S rRNA | 0 | 8 | 0 | 8 | 0 | 5 | 9 | 9 |
| SEC A | 0 | 2 | 1 | 2 | 0 | 0 | 5 | 5 |

## Discussion

A variety of significant events influenced begomovirus population proliferation, resulting in the evolution of segmented DNA sequences in these viruses (Sohrab et al., 2016a; Rojas et al., 2005). The diverse host range of begomoviruses is a result of various farming methods and the vector’s polyphagous nature (whitefly). As a result of begomovirus’s evolution and the new strains emerging within them, the beginning of a devastating threat to the farming of chillies and major trouble for universal agriculture has been observed (Leke et. al., 2015; Varma et al., 2005). A variety of factors influence virus genetic diversity, the most important of which is a mutation (Rubio et al., 2020; Roossinck, 1997; Balol et al., 2010). A higher mutation rate facilitates the evolution of viral genotypes that are more adapted to living in demanding new environments (Pagan and Garcia-Arenal, 2018, Barton et al., 2010; Sanjua’n et al., 2010). Recombination is a second, even more, essential factor that contributes to the high variety of plant viruses’ diversity (Rubio et al., 2020; Martin et al., 2011).In addition to these two critical factors, the upsurge in nucleotide substitution is significant in evolution, which explains why begomoviruses have a high genetic level of variation and continue to evolve as swiftly as RNA viruses (Duffy and Holmes, 2009; Lefeuvre et al., 2011; Lima et al., 2017; Moya et al., 2004; Drake et al., 1999). For monopartite begomoviruses, betasatellite mostly causes the onset of characteristic symptoms signs (Gnanasekaran et al., 2019; Li et al., 2018). ChiLCV is one of the most destructive begomoviruses, significantly reducing yields in chilli farming around the globe (Mishra et al., 2022; Pandey et al., 2022). Numerous plants are susceptible to ChiLCV infection, including hibiscus, tomato, papaya, eggplant, and chilli (Malathi et al., 2017; Mishra et al., 2022; Srivastava et al., 2022). Similarly, phytoplasmas induce approximately 300 distinct plant diseases, including both wild and cultivated species of plants of various families, inflicting substantial harm to the economy globally (Kumari et al., 2019; Rao et al., 2011; Hoshi et al., 2007). The association of complex phytoplasma and viruses with a variety of plant diseases has previously been studied (Arocha et al., 2009). Thus, to establish symptoms epidemiology and the existence of both infection-producing pathogens, appropriate disease diagnostics were essential. Therefore, to trace and characterise the disease-causing agents in chilli plants, we performed collection of symptomatic chilli leaf, isolation, and identification of pathogens with detailed molecular computational analysis for begomovirus and phytoplasma isolates associated with mixed infection in chilli plants. In this study, two pathogens were used: ChiLCV and its associated ChLCuB from begomovirus and *Candidatus Phytoplasma trifolii* 16S rRNA, Sec-A gene. To the best of our awareness, this is the first time a closely integrated and detailed comparative *in silico* study of begomovirus (ChiLCV-A and ChLCuB) and phytoplasma (16S rRNA and Sec-A) has been performed, and the symptoms was validated by the full-length sequence of these identified components (ChiLCV along with ChLCuB isolate GKP and *Candidatus Phytoplasma trifolii* 16S rRNA and Sec-A gene).

Although recombination and mutation are the two main causes of genetic variability, other processes like genetic drift and natural selection also have a direct impact on the emergence and evolution of plant viruses (Rubio et al., 2020; Lima et al., 2017; Barton et al., 2010). This research also shows that, despite sequences exhibiting nucleotide identities above 90 %, their transcript levels of similarity were lower. In our phylogenetic investigation, we observed that, at the clustering bunches, all the isolates differed for the ChiLCV-GKP and its six ORFs, demonstrating the role of recombination in genetic variation at the gene level. Additionally, we found that the bias and transition substitution among ChiLCV-A ORFs varied, with AV1 having the highest value. Similarly, we observed differences in transition and transversion substitutions/bias for 16S rRNA and Sec A gene readings that were exceptionally significant when compared to ChiLCV isolates and their coding region. In addition to this, the present study established the importance of recombination events in genetic diversity via relevant target region breakpoints in ChiLCV-A and ChLCuB, which are responsible for the increased pace of evolution and host range extension. We noticed 21 breakpoints for our GKP isolate of ChiLCV-A and 11 for associated ChLCuB, along with maximum breakpoints in AV1/CP, which indicates its crucial role in recombination among this isolate. In contrast, the results for the Sec-A gene and the 16S rRNA of *Candidatus Phytoplasma trifolii* did not reveal any significant breakpoints.

Although RNA viruses have a considerably higher nucleotide substitution rate than dsDNA, in the case of begomovirus (ssDNA), the mean rate of nucleotide substitution was found to be comparable to RNA viruses, i.e., 10^−2^-10^−5^ nucleotide substitution /site/year (Duffy et al., 2008; Pagán and Garca-Arenal 2018; Sanjua’n et al., 2010). As a result, geminiviruses have remarkable genetic variability. In this investigation, we noticed that Rep had the maximum active participation in variability with 9.62×10^−4^ substitutions/site/year (within the 95% HPD interval of 5.64×10^−5^ to 2.28×10^−3^) but CP had the highest mutation rate among all ORFs at its wobble codon position, reinforcing the standpoint that its involvement in genetic mutations leads to genetic variability. However, in the case of phytoplasma, we noticed a very positive outcome for 16s rRNA of 1.639E-4 substitutions/site/year [6.2133E-5, 2.606E-4] and 7.0218E-4 substitutions/site/year [2.4295E-3, 1.14E-2], which was less than ChiLCV isolated but comparable to many ORFs, suggesting that phytoplasma is indeed moving towards high genetic variability. While examining both begomoviral and *Ca.P.trifolii* phytoplasmas components, we noticed that the nucleotide diversity **(π)** and the total number of mutations **(η)** for phytoplasma were in the range of Ren and AC4 for 16S rRNA but lower for Sec A (**Table 7 and Table 8**). Interestingly, the negative results of all neutrality tests indicate that our ChiLCV isolate and its associated betasatellite are under purifying selection, indicating that the gene’s nature appears conserved. Positive results indicate the sudden population contraction in the case of the CP gene. Ren, AC4 and betasatellite, which exhibited diversifying and positive selection, fared exceptionally well, whereas phytoplasma did not. According to the current study’s *in silico* results revealed, these two pathogens rapidly evolved with significant genetic variability to the chilli crop. The strength of plant virus evolution has led to numerous instances where phytoviruses not only infect plants but also spread and multiply in insect vectors and decrease the longevity and fecundity of insects (Diaz-Pendon et al., 2010; Ghanim et al., 1998; Bosco et al., 2004; Sylvester et al., 1980; Rubinstein et al., 1997). Additionally, plant viruses can interact with human cells as well, although conclusive evidence is still lacking (Balique et al., 2015; Rebolledo-Mendez et al., 2013).

## Conclusion

This is the first time phytoplasma (*Ca.P.trifoliĩ*) has been found in a association with ChiLCV species from the chilli crop, and a large computational evaluation of these two destructive pathogens has been undertaken. The overall study shows the impact of genetic variability in begomovirus and phytoplasma among economically important chilli cultivars. Our results indicate that the Gkp isolate for ChiLCV-A, as well as betasatellite segments of known begomoviruses and components of *Candidatus Phytoplasma trifolii* (16S rRNA and Sec A), exhibit diverse evolutionary patterns with significant variation and recombination events. As a result, regulatory regions critical for genome structure and functionality may be under significant selection pressure in ChiLCV-GKP isolate segments. Considering that each segment can function as a separate independent autonomous fragment and is the consequence of genetic material exchange between similar begomoviruses. This current study will pave the way for the development of a management strategy to protect crops from the enormous threat posed by several leaf curl viruses and phytoplasma. Studying these mixed infections in different hosts with classic mixed symptoms is crucial because it will address the epidemiological consequences of viral diversity as well as future therapies.

## Data Availability Statement

The original contributions presented in the study are included in the article/Supplementary Material; further inquiries can be directed to the corresponding authors.

## Author Contributions

VP and AS performed experiments and wrote the manuscript. RG and SM guided the design of the whole test scheme, and AA-S and MS critically reviewed the manuscript and aided in the evaluation of certain datasets and bioinformatics approaches. All authors contributed to the article and approved the submitted version.

## Acknowledgements

AA-S and MS are thankful to Sultan Qaboos University, Oman for financial support.

## References

Agricultural and Processed Food Products Export Development Authority (APEDA); 2021. https://agriexchange.apeda.gov.in/India%20Production/India_Productions.aspx?hscode=1098 [Accessed October 11, 2022].

Aljanabi S, Parmessur Y, Moutia Y, Saumtally S, Dookun A. Further evidence of the association of a phytoplasma and a virus with yellow leaf syndrome in sugarcane. Plant Pathol. 2008; 50:628–636. doi:10.1046/j.1365-3059.2001.00604.x

Altschul SF, Gish W, Miller W, Myers EW, Lipman DJ. Basic local alignment search tool. J. of Mol. Biol. 1990; 215:403–10. doi:10.1016/S0022-2836(05)80360-2.

Amaral-Mello APO, Bedendo IP, Kitajima EW, Ribeiro LF, Kobori R. Tomato big bud associated with a phytoplasma belonging to group 16Sr III in Brazil. Int. J. Pest Manage. 2006; 52:233–237. doi:10.1080/09670870600733766

Arocha Y, Pinol B, Acosta K, Almeida R, Devonshire J, et al. Detection of phytoplasma and potyvirus pathogens in papaya (Carica papaya L.) affected with “Bunchy top symptom” (BTS) in eastern Cuba. Crop Prot. 2009; 28:640–646. doi:10.1016/j.cropro.2009.03.020

Balique F, Hervé Lecoq, Raoult D, Colson P. Can Plant Viruses Cross the Kingdom Border and Be Pathogenic to Humans? Viruses 2015; 7:2074–2098. doi: 10.3390/v7042074

Barton NH. Mutation and the evolution of recombination. Phil. Trans. R. Soc. B. 2010;365: 1281–1294. doi:10.1098/rstb.2009.0320

Balol GB, Divya BL, Basavaraj S, Sundaresha S, Mahesh YS, et al. Sources of genetic variation in plant virus populations. J. Pure Appl Microbiol. 2010;4(2): 803–808.

Bosco D, Mason G, Accotto GP. TYLCSV DNA, but not infectivity, can be transovarially inherited by the progeny of the whitefly vector *Bemisia tabaci* (Gennadius). Virology 2004; 323:276–283. doi 10.1016/J.VIROL.2004.03.010

Briddon RW, Bull SE, Mansoor S, Amin I, Markham PG. Universal primers for the PCR-mediated amplification of DNA beta: a molecule associated with some monopartite begomoviruses. Mol. Biotech. 2002; 20:315–318. doi:10.1385/MB:20:3:315

Briddon RW, Martin DP, Roumagnac P, Navas-Castillo J, Fiallo-Olive E, et al. Alphasatellitidae: A New Family with Two Subfamilies for the Classification of Geminivirus-and Nanovirus-Associated Alphasatellites. Arch. of Virol. 2018. 163;2587–2600. doi:10.1007/s00705-018-3854-2.

Chaturvedi Y, Rao GP, Tewari AK, Duduk B, Bertaccini A. Phytoplasma in ornamentals: detection, diversity and management. Acta Phytopath. Entomol. Hung. 2010; 45:31–69. doi:10.1556/APhyt.45.2010.1.3

Deng S, Hiruki C. Amplification of 16S rRNA genes from culturable and nonculturable mollicutes. J. Microbiol. Meth. 1991;14: 53e61. doi:10.1016/0167-7012(91)90007-D

Drake JW, John J, Holland JJ. Mutation rates among RNA viruses. PNAS 1999;96 (24):13910–13913.doi: 10.1073/pnas.96.24.13910

Diaz-Pendon JA, Canizares MC, Moriones E, Bejarano ER, Czosnek H, et al. Tomato yellow leaf curl viruses: Menage trois between the virus complex, the plant and the whitefly vector. Mol. Plant Pathol. 2010;11, 441–450. doi: 10.1111/j.1364-3703.2010.00618.x

Duffy S, Holmes EC. Phylogenetic evidence for rapid rates of molecular evolution in the single-stranded DNA begomovirus Tomato yellow leaf curl virus J. Virol. 2008; 82:957–965. doi:10.1128/JVI.01929-07

Duffy S, Holmes EC. Validation of high rates of nucleotide substitution in geminiviruses: phylogenetic evidence from East African cassava mosaic viruses. J Gen Virol. 2009; 90,1539–1547. doi:10.1099/vir.0.009266-0

Ermacora P, Osler R. Symptoms of Phytoplasma Diseases eds R. Musetti and L. Pagliari (Springer:Humana Press), 2019;1875:53–67. doi: 10.1007/978-1-4939-8837-2_5.

Fiallo-Olive E, Jean-Michel L, Martin DP, Roumagnac P, Varsani A, et al. ICTV Virus Taxonomy Profile Geminiviridae. J. of Gen. Virol. 2021; 102:001696. doi: 10.1099/jgv.0.001696

Fiallo-Olive E, Tovar R, Navas-Castillo J. Deciphering the Biology of Deltasatellites from the New World: Maintenance by New World Begomoviruses and Whitefly Transmission. New Phytol. 2016;212, 680–692. doi:10.1111/nph.14071.

Food and Agriculture Organization (FAO). 2020; https://www.fao.org/faostat/en/#data/QCL [Accessed October 11, 2022].

Ganefianti DW, Hidayat SH, Syukur M. Susceptible Phase of Chili Pepper Due to Yellow Leaf Curl Begomovirus Infection. IJASEIT 2017;7(2):594. doi:10.18517/ijaseit.7.2.1872

Ghanim M, Morin S, Zeidan M, Czosneck H. Evidence for transovarial transmission of tomato yellow leaf curl virus by its vector, the whitefly Bemisia tabaci. Virology 1998; 240: 295–303. doi 10.1006/VIRO.1997.8937

Gibbs A, Ohshima K. Potyviruses and the digital revolution. Annu. Rev. Phytopathol. 2010; 48:205–223. doi: 10.1146/annurev-phyto-073009-114404

Girsova NV, Bottner-Parker KD, Bogoutdinov DZ, Kastalyeva TB, Meshkov_YI. et al. Diverse phytoplasmas associated with leguminous crops in Russia. Euro. J. of Plant Pathol., 2017;149(3):599–610. doi: 10.1007/s10658-017-1209-6

Gnanasekaran P, Kumar KR, Tacharyya DR, Kumar RV, Chakraborty S. Multifaceted role of geminivirus associated betasatellite in pathogenesis. Mol Plant Pathol 2019;20(7):1019–1033. doi:10.1111/mpp.12800

Goodman R M. Geminiviruses. J. Gen. Virol. 1981; 54:9–21. doi: 10.1186/1471-2148-9-112

Gundersen DE, Lee IM. Ultrasensitive detection of phytoplasmas by nested PCR assays using two universal primer pairs. Phytopath. Mediterr. 1996; 35:144e151.

Gutierrez C. Geminiviruses and the plant cell cycle. Plant Mol. Biol. 2000;43:763–772. doi: 10.1023/a:1006462028363

Hall TA. BioEdit: A User-Friendly Biological Sequence Alignment Editor and Analysis Program for Windows 95/98/NT. Nucleic Acids Symp. Series. 1999;41: 95–98.

Haible D, Kober S, Jeske H. Rolling circle amplification revolutionizes diagnosis and genomics of geminiviruses. J. of Virol. Meth. 2006;135:9–16. doi:10.1016/j.jviromet.2006.01.017.

Hodgetts J, Boonham N, Mumford R, Harrison N, Dickinson M. Phytoplasma phylogenetics based on analysis of secA and 23S rRNA gene sequences for improved resolution of candidate species of ‘Candidatus Phytoplasma’. Int. J. Syst. Evol. Microbiol. 2008;58:1826e1837. doi: 10.1099/ijs.0.65668-0

Hoshi A, Ishii Y, Kakizawa S, Oshima K, Namba S. Hostparasite interaction of phytoplasmas from a molecular biological perspective. Bull. Insectol. 2007;60, 105–107

Huson, D.H., Bryant, D. Application of Phylogenetic Networks in Evolutionary Studies. Mol. Bio. and Evol., 2006;23:254–267. doi:10.1093/molbev/msj030.

Kaminska M, Sliwa H, Rudzinska-Langwald A. Shoot proliferation of bleeding heart (Dicentra spectabilis) associated with phytoplasma and virus infection. Phytopathol. 2005; 37: 33–4

Khan ZA, Khan JA. Characterization of a new begomovirus and betasatellite associated with chilli leaf curl disease in India. Arch. Virol. 2007; 162:561–565. doi: 10.1007/s00705-016-3096-0

Kumar S, Stecher G, Li M, Knyaz C, Tamura K. MEGA X: molecular evolutionary genetics analysis across computing platforms. Mol. Biol. Evol. 2018;35(6): 1547–1549. doi:10.1093/molbev/msy096

Kumar RV, Singh AK, Singh AK, Yadav T, Basu S. Complexity of begomovirus and betasatellite populations associated with chilli leaf curl disease in India. J. Gen. Virol. 2015;96:3143–3158. doi:10.1099/jgv.0.000254.

Kumari S, Nagendran K, Rai AB, Singh B, Rao GP et al. Global Status of Phytoplasma Diseases in Vegetable Crops. Front. Microbiol. 2019:10:1349. doi: 10.3389/fmicb.2019.01349

Lavanya R, Arun V. Detection of Begomovirus in chilli and tomato plants using functionalized gold nanoparticles. Scient. Reports 2021;11, 14203 doi: s41598-021-93615-9

Lebsky V, Hernandez-Gonzalez J, Arguello-Astorga G, Cardenasconejo Y, Poghosyan A. Detection of phytoplasmas in mixed infection with begomoviruses: a case study of tomato and pepper in Mexico. Bull. Insectol. 2011;64: 55–56.

Lee I M, Martini M, Marcone C, Zhu S F. Classification of phytoplasma strains in the elm yellows group (16SrV) and proposal of ‘Candidatus Phytoplasma ulmi’ for the phytoplasma associated with elm yellows. In. J. of Sys. and Evol. Microbio. 2004;54(2):337–347. doi: 10.1099/ijs.0.02697-0

Lee IM, Zhao Y, Davis RE. Prospects to multiple gene-based systems for differentiation and classification of phytoplasmas. In: Weintraub, P.G., Jones, P. (Eds.), (Phytoplasmas Genomes, Plant Hosts and Vectors: CAB International, UK), 2010b;51–63. doi:10.1079/9781845935306.0051

Lefeuvre P, Harkins GW, Lett JM, Briddon RW, Chase MW, Moury B. Evolutionary time-scale of the begomoviruses: evidence from integrated sequences in the Nicotiana genome. PLoS ONE 2011;6(5):e19193. doi:10.1371/journal.pone.0019193

Leke WN, Mignouna DB, Brown JK, Kvarnheden A. Begomovirus disease complex: emerging threat to vegetable production systems of West and Central Africa. Agri. & Food Security 2015; 4:1. doi: 10.1186/s40066-014-0020-2

Letunic I, Bork P. Interactive Tree of Life (iTOL) v5: an online tool forphylogenetic tree display and annotation. Nucleic Acids Res. 2021; 49(1):293–296. doi:10.1093/nar/gkab301

Li F, Yang X, Bisaro DM., Zhou X. The beta C1 protein of geminivirus-betasatellite complexes: a target and repressor of host defenses. Mol Plant. 2018; 11:1424–1426. doi:10.1016/j.molp.2018.10.007

Lima A, Silva JC, Silva FN, Castillo-Urquiza GP, Silva F. et al. The diversification of begomovirus populations is predominantly driven by mutational dynamics. Virus Evol. 2017; 3(1),1–14. doi:1093/ve/vex005

Malathi VG, Renukadevi P, Chakraborty S, Biswas KK, Roy A. et al. “Begomoviruses and Their Satellites Occurring in India: Distribution, Diversity and Pathogenesis”, in a Century of Plant Virology in India, Eds. B. Mandal, G.P. Rao, V.K. Baranwal, R.K. Jain (Springer: Singapore), 2017; 75–177.

Mall S, Chaturvedi Y, Rao GP, Barnwal VK. Phytoplasma’s diversity in India. Bull. Insectol. 2011; 64:1721–8861.

Martin DP, Biagini P, Lefeuvre P, Golden M, Roumagnac P, et al. Recombination in eukaryotic single stranded DNA viruses. Viruses. 2011;3(9): 1699–1738. doi:10.3390/v3091699

Martin DP, Murrell B, Golden M, Khoosal A, Muhire B. RDP4: detection and analysis of recombination patterns in virus genomes. Virus Evol. 2015;1(1):vev003. doi: 10.1093/ve/vev003

Marwal A, Nehra C, Verma RK., Mishra M, Srivastava D. First report of papaya leaf curl virus and its associated papaya leaf curl betasatellite infecting *Catharanthus roseus* plants in India. The J. of Horticult. Sci. Biotech. 2021 doi:10.1080/14620316/2021/1912646

Mishra M, Verma RK, Pandey_V, Srivastava A, Sharma P. Role of Diversity and Recombination in the Emergence of Chilli Leaf Curl Virus. Pathogens. 2022;11(5):529. doi: 10.3390/pathogens11050529.

Mishra M, Verma RK, Marwal A, Sharma P, Gaur RK. Biology and Interaction of the Natural Occurrence of Distinct Monopartite Begomoviruses Associated with Satellites in *Capsicum annum* from India. Front. in microbiol. 2020;11:512957. doi: 10.3389/fmicb.2020.512957

Moya A, Holmes EC, González-Candelas F. The population genetics and evolutionary epidemiology of RNA viruses. Nat. Rev. Microbiol. 2004;2:279–288. doi: 10.1038/nrmicro863

Moffat AS. Geminiviruses emerge as serious crop threat. Science 1999; 286, 1835–1835. doi:10.1126/science.286.5446.1835

Mubin M, Ijaz S, Nahid N, Hassan M, Younus A. Journey of Begomovirus Betasatellite Molecules: From Satellites to Indispensable Partners. Virus Genes. 2020;56:16–26. doi:10.1007/s11262-019-01716-5.

Muhire BM, Varsani A, Martin DP. SDT: a virus classification tool based on pairwise sequence alignment and identity calculation. PLOS One. 2014;26:9:e108277. doi:10.1371/journal.pone.0108277.

Pagan I, Garcia-Arenal F. Population Genomics of Plant Viruses, in Population Genomics: Microorganisms, eds M. F. Polz and O. P. Rajora. (Cham: Springer International Publishing), 2018; 233–265. doi:10.1007/13836_2018_15

Pandey V, Srivastava A, and Gaur RK. Begomovirus: a curse for the agricultural crops. Arch. of Phytopathol.and Plant Prot‥ 2021;54(15-16):949–978. doi:10.1080/03235408.2020.1868909

Pandey V, Srivastava A, Mishra M, Gaur RK. Chilli leaf curl disease populations in India are highly recombinant, and rapidly segregated. 3 Biotech. 2022; 12(3):83. doi:10.1007/s13205-022-03139-w.

Patil BL, Fauquet CM. Differential Interaction between Cassava Mosaic Geminiviruses and Geminivirus Satellites. J. of Gen. Virol. 2010;91:1871–1882, doi:10.1099/vir.0.019513-0.

Pratap D, Ashwin RK, Mukherjee SK. Molecular characterization and infectivity of a Tomato leaf curl New Delhi virus variant associated with newly emerging yellow mosaic disease of eggplant in India. Virol J. 2011;8, 305. doi:10.1186/1743-422X-8-305

Rambaut A, Drummond AJ, Xie D, Baele G, Suchard MA. Posterior summarization in bayesian phylogenetics using tracer. Syst Biol. 2018;7(5): 901–904. doi:10.1093/sysbio/syy032

Rao G P, Madhupriya T V, Manimekalai R, Tiwari A K, Yadav A. A century progress of research on phytoplasma diseases in India. Phytopatho.Mollicutes 2017;7:1–38. doi:10.5958/2249-4677.2017.00001.9

Rao GP, Mall S, Raj SK, Sneha SK. Phytoplasma disease affecting various plant species in India. Acta Phytopathol. Entomol. Hung. 2011; 46:59–99. doi:10.1556/APhyt.46.2011.1.7

Rao GP, Rao A, Kumar M, Ranebennur H, Mitra S. Identification of phytoplasma in six fruit crops in India. Eur J Plant Pathol. 2021; 156:1197–1206 doi: 10.1007/s10658-020-01949-3

Rebolledo-Mendez JD, Vaishnav RA, Cooper NG, Friedland RP. Cross-kingdom sequence similarities between human micro-RNAs and plant viruses. Commun. Integr. Biol. 2013;6: e24951. doi: 10.4161/cib.24951

Rojas M R. Macedo MA, Maliano MR, Soto-Aguilar M, Souza JO, et al. World management of geminiviruses Annu. Rev. Phytopathol. 2018; 56, 637–677. doi:10.1146/annurev-phyto-080615-100327

Rojas M R, Hagen C, Lucas WJ, Gilbertson R L. Exploiting chinks in the plant’s armor: evolution and emergence of geminiviruses. Annual Review of Phytopathology 2005; 43:361–394. doi: 10.1146/annurev.phyto.43.040204.135939

Rojas MR, Gilbertson RL, Russell DR, Maxwell DP. Use of degenerate primers in the polymerase chain reaction to detect whitefly-transmitted geminiviruses. Plant Dis. 1993; 77:340–340. doi:10.1094/PD-77-0340

Roossinck MJ. Mechanisms of plant virus evolution. Annu Rev Phytopathol 1997; 35:1953–1965. doi:10.1146/annurev.phyto.35.1.19

Rozas J, Ferrer-Mata A, Sánchez-DelBarrio JC, Guirao-Rico S, Librado P. et al. DnaSP 6: DNA Sequence Polymorphism Analysis of Large Data Sets. Mol. Bio. and Evol. 2017; 34:3299–3302, doi:10.1093/molbev/msx248.

Rubio L, Galipienso L, Ferriol I. Detection of Plant Viruses and Disease Management: Relevance of Genetic Diversity and Evolution. Front. in Plant Sci. 2020;11:1–23. doi: 10.3389/fpls.2020.01092

Rubinstein G, Czosnek H. Long-term association of tomato yellow leaf curl virus with its whitefly vector Bemisia tabaci: Effect on the insect transmission capacity, longevity and fecundity. J. Gen. Virol. 1997:78,2683–2689. Doi 10.1099/0022-1317-78-10-2683

Sahu AK, Verma RK, Gaur RK, Sanan-Mishra N. Complexity and recombination analysis of novel begomovirus associated with Spinach yellow vein disease in India. Plant Gene. 2018;13:42–49. doi: 10.1016/j.plgene.2018.01.001

Sanjua’n R, Nebot MR, Chirico N, Louis M, Mansky LM, et al. Viral Mutation Rates. J. of Virol. 2010;84(19), 9733–9748. Doi: doi:10.1128/JVI.00694-10

Scholthof KBG, Adkins S, Czosnek H, Palukaitis_P, Jacquot,E, et al. Top 10 plant viruses in molecular plant pathology. Mol. Plant Pathol. 2011;12, 938–954. doi: 10.1111/j.1364-3703.2011.00752.x

Smart CD, Schneider B, Blomquist CL, Guerra LJ, Harrison NA, et al. Phytoplasma specific PCR primers based on sequences of 16S-23SrRNA spacer region. Appl. Env. Microbiol. 1996; 62:2988–2993. doi: 10.1128/aem.62.8.2988-2993.1996

Sohrab SS, Yasir M, El-Kafrawy SA, Al-Zahrani HSM, Mousa MAA, et al. Phylogenetic relationships, recombination analysis and genetic variability of Tomato yellow leaf curl virus infecting tomato in Jeddah, Saudi Arabia. Plant Omics J. 2016a;9(1):90–98.

Sohrab SS, Ayman T Abbas, Yasir M, Mousa MAA, El-Kafrawy SA, et al. Association of tomato leaf curl Sudan virus with leaf curl disease of tomato in Jeddah, Saudi Arabia. Virus Diseases 2016b;27(2):145–153. doi: 10.1007/s13337-016-0308-x

Sohrab SS, Yasir M, El-Kafrawy SA, Mousa MAA, Bakhashwain AA. First report of Begomovirus causing yellow mosaic disease of ridge gourd in Saudi Arabia. Spain 6th Euro Virol. Congress and Expo.5,1. Abstract retrieved from Virol-mycol. 2016c. doi: 10.4172/2161-0517.C1.009

Srivastava A, Pandey V, Sahu_AK, Yadav D, Al-Sadi AM, et al. Evolutionary Dynamics of Begomoviruses and Its Satellites Infecting Papaya in India. Front. Microbiol. 2022; 13:879413. doi: 10.3389/fmicb.2022.879413

Srivastava A, Pandey V, Verma R K, Marwal A, Mishra R, et al. First complete genome sequence of Tomato leaf curl virus (ToLCV) from *Salvia splendens* in India. J. of Phytopathol. 2022a;170: 479–491. doi:10.1111/jph.13099

Suchard MA, Lemey P, Baele G, Ayres DL, Drummond A J, et al. Bayesian Phylogenetic and Phylodynamic Data Integration Using BEAST 1.10. Virus Evol. 2018; 4(1): vey016. doi:10.1093/ve/vey016

Sylvester ES. Circulative and propagative virus transmission by aphids. Ann. Rev. Entomol. 1980; 25:257–286. doi:10.1146/annurev.en.25.010180.001353

Varma A,_Malathi VG. Emerging geminivirus problems: A serious threat to crop production. Annu. of App. Bio. 2005;142(2):145–164. doi: 10.1111/j.17447348.2003.tb00240.x

Weaver S, Shank SD, Spielman SJ, Li M, Muse SV, et al. Datamonkey 2.0: A Modern Web Application for Characterizing Selective and Other Evolutionary Processes. Mol. Bio. and Evol. 2018;35: 773–777, doi:10.1093/molbev/msx335

Wei W, Lee I-M, Davis RE, Suo X, Zhao Y. Automated RFLP pattern comparison and similarity coefficient calculation for rapid delineation of new and distinct phytoplasma 16Sr subgroup lineages. Int. J. Syst. Evol. Microbiol. 2008; 58:2368–2377. doi:10.1099/ijs.0.65868-0

Zhao Y, Wei W, Lee I-M, Shao J, Suo X, et al. Construction of an interactive online phytoplasma classification tool, iPhyClassifier, and its application in analysis of the peach X-disease phytoplasma group (16SrIII). Int. J. Syst. Evol. Microbiol. 2009; 59:2582–2593. doi:10.1099/ijs.0.010249-0

Zhou X. Advances in Understanding Begomovirus Satellites. Annual Review of Phytopathol. 2013;51, 357–381. doi:10.1146/annurev-phyto-082712-102234

