## Supplemental Table S11 for "Perilous coexistence: *Chilli Leaf Curl Virus* and *Candidatus Phytoplasma trifolii* infecting *Capsicum annuum*, India"

**Table. S11** Results of different neutrality tests analysis *Ca.P. trifolii* 16S rRNA (Acc. No. MZ557805) and *Ca.P.trifolii* Sec A gene (Acc. No. MZ620707).

| **Phytoplasma component** | **Neutrality tests** | | |
|---|---|---|---|
| | **Tajima's *D*** | **Fu & Li's *D*** | **Fu & Li's *F*** |
| **16S rRNA** | -2.31994 | -7.31755 | -6.30638 |
| **Sec-A** | -2.17993 | -2.74409 | -3.01383 |
