## Supplemental Table S1 for "Perilous coexistence: *Chilli Leaf Curl Virus* and *Candidatus Phytoplasma trifolii* infecting *Capsicum annuum*, India"

**Table. S1** ORFs details of ChiLCV-A-GKP (Acc. No. MZ540908) and ChLCuB-GKP (Acc. No. MZ540909) similarity score with other begomovirus geographical isolates.

| Virus isolate | Accession number | ORFs | Range | Maximum nucleotide % Identity with other isolate | Maximum Amino Acid % identity |
|---|---|---|---|---|---|
| **ChiLCV-A/GKP** | | | | | |
| | **MZ540908** | **AV1** | 267-1082 | **MK757213**_94.201 | **MK757213**_87.801 |
| | | **AV2** | 149-514 | | |
| | | **AC1** | 1531-2616 | | |
| | | **AC2** | 1242-1628 | | |
| | | **AC3** | 1079-1483 | | |
| | | **AC4** | 2166-2429 | | |
| **ChLCuB /GKP** | | | | | |
| | **MZ540909** | **βC1** | 220-582 | **MT385295**_97.801 | **DQ343289**_50.601 |
