## Supplemental Table S2 for "Perilous coexistence: *Chilli Leaf Curl Virus* and *Candidatus Phytoplasma trifolii* infecting *Capsicum annuum*, India"

**Table. S2** List of begomovirus isolates collected from GENBANK, NCBI, USA, based on BLAST similarity with our identified ChiLCV-A-GKP (Acc. No. MZ540908) isolate and organized in listed in order of similarity score.

| **Accession no.** | **Species** | **Acronym** | **Year** | **Country** | **Host** |
|---|---|---|---|---|---|
| MZ540908 | *Chilli leaf curl virus* | ChiLCV-[IN:GKP:Chi:21] | 2021 | India | *Capsicum annuum* |
| MK757213 | *Chilli leaf curl virus* | ChiLCV-[OM:Tom30-19:Tom:16] | 2016 | Oman | *Solanum lycopersicum* |
| HE806437 | *Chilli leaf curl virus* | ChiLCV-[OM:T-37:11] | 2011 | Oman | *Solanum lycopersicum* |
| MK757215 | *Chilli leaf curl virus* | ChiLCV-[OM:Tom32-20:16] | 2016 | Oman | *Solanum lycopersicum* |
| MK757214 | *Chilli leaf curl virus* | ChiLCV-[OM:Tom31-23:16] | 2016 | Oman | *Solanum lycopersicum* |
| MK757212 | *Chilli leaf curl virus* | ChiLCV-[OM:Tom26-12:16] | 2016 | Oman | *Solanum lycopersicum* |
| MK468696 | *Chilli leaf curl virus* | ChiLCV-[OM:Tob-13:15] | 2015 | Oman | *Nicotiana tabacum* |
| MK468694 | *Chilli leaf curl virus* | ChiLCV-[OM:Tob-11:15] | 2015 | Oman | *Nicotiana tabacum* |
| MN119490 | *Chilli leaf curl virus* | ChiLCV-[OM:P14:Pum:16] | 2016 | Oman | *Cucurbita maxima* |
| KX787939 | *Chilli leaf curl virus* | ChiLCV-[OM:Wat-122:15] | 2015 | Oman | *Citrullus lanatus* |
| KF229718 | *Chilli leaf curl virus* | ChiLCV-[OM:Tom-26:12] | 2012 | Oman | *Solanum lycopersicum* |
| MN119491 | *Chilli leaf curl virus* | ChiLCV-[OM:P5:Pum:16] | 2016 | Oman | *Cucurbita maxima* |
| HG969264 | *Chilli leaf curl virus* | ChiLCV-[OM:Rad-4:13] | 2013 | Oman | - |
| MN119492 | *Chilli leaf curl virus* | ChiLCV-[OM:P6:Pum:16] | 2016 | Oman | *Cucurbita maxima* |
| HG969265 | *Chilli leaf curl virus* | ChiLCV-[OM:C1:13] | 2013 | Oman | - |
| HG969263 | *Chilli leaf curl virus* | ChiLCV-[OM:Rad-3:13] | 2013 | Oman | - |
| HF968755 | *Chilli leaf curl virus* | ChiLCV-[OM:PT-2:12] | 2012 | Oman | *Petunia* |
| HF968756 | *Chilli leaf curl virus* | ChiLCV-[OM:PT-3:12] | 2012 | Oman | *Petunia* |
| LN680633 | *Chilli leaf curl virus* | ChiLCV-[OM:Tom112:14] | 2014 | Oman | *Solanum lycopersicum* |
| MF737343 | *Chilli leaf curl virus* | ChiLCV-[IN:MM1:17] | 2017 | India | *Capsicum annuum* |
| HG969257 | *Chilli leaf curl virus* | ChiLCV-[OM:HP-12:13] | 2013 | Oman | - |
| LN680625 | *Chilli leaf curl virus* | ChiLCV-[OM:Tom85:14] | 2014 | Oman | *Solanum lycopersicum* |
| LN680626 | *Chilli leaf curl virus* | ChiLCV-[OM:Tom88:14] | 2014 | Oman | *Solanum lycopersicum* |
| HG969197 | *Chilli leaf curl virus* | ChiLCV-[OM:Tom84:14] | 2014 | Oman | *Solanum lycopersicum* |
| LN680624 | *Chilli leaf curl virus* | ChiLCV-[OM:Tom87:14] | 2014 | Oman | *Solanum lycopersicum* |
| HG969255 | *Chilli leaf curl virus* | ChiLCV-[OM:HP-5:13] | 2013 | Oman | - |
| JN604495 | *Chilli leaf curl virus* | ChiLCV-[OM:Taq4:Tom11] | 2011 | Oman | *Solanum lycopersicum* |
| JN604498 | *Chilli leaf curl virus* | ChiLCV-[OM:Th3:Tom:11] | 2011 | Oman | *Solanum lycopersicum* |
| JN604494 | *Chilli leaf curl virus* | ChiLCV-[OM:Taq3:Tom:11] | 2011 | Oman | *Solanum lycopersicum* |
| HG941644 | *Chilli leaf curl virus* | ChiLCV-[OM:TB-23:Tob:13] | 2013 | Oman | *Nicotiana tabacum* |
| JN604499 | *Chilli leaf curl virus* | ChiLCV-[OM:Th4:Tom:11] | 2011 | Oman | *Solanum lycopersicum* |
| JN604493 | *Chilli leaf curl virus* | ChiLCV-[OM:Taq2:Tom:11] | 2011 | Oman | *Solanum lycopersicum* |

| | | | | | |
|---|---|---|---|---|---|
| JN604497 | *Chilli leaf curl virus* | ChiLCV-[OM:Th2:Tom:11] | 2011 | Oman | *Solanum lycopersicum* |
| HG941643 | *Chilli leaf curl virus* | ChiLCV-[OM:BZ-8:13] | 2013 | Oman | - |
| MH475359 | *Chilli leaf curl virus* | ChiLCV-[OM:Wed8-25:UI:16] | 2016 | Oman | *Urtica incisa* |
| JN604496 | *Chilli leaf curl virus* | ChiLCV-[OM:Th1:Tom:11] | 2011 | Oman | *Solanum lycopersicum* |
| MH475358 | *Chilli leaf curl virus* | ChiLCV-[OM:Wed8-24:UI:16] | 2016 | Oman | - |
| JN604500 | *Chilli leaf curl virus* | ChiLCV-[OM:KW1:Tom:11] | 2011 | Oman | *Solanum lycopersicum* |
| JN604491 | *Chilli leaf curl virus* | ChiLCV-[OM:Sh1:Pep:11] | 2011 | Oman | *Capsicum annuum* |
| JN604490 | *Chilli leaf curl virus* | ChiLCV-[OM:Sal2:Pep:11] | 2011 | Oman | *Capsicum annuum* |
| JN604489 | *Chilli leaf curl virus* | ChiLCV-[OM:Sal1:Pep:11] | 2011 | Oman | *Capsicum annuum* |
| EU939533 | *Chilli leaf curl virus* | ChiLCV-[IN:Nar:Pep:04] | 2004 | India | *Capsicum annuum* |
| AF336806 | *Chilli leaf curl virus* | ChiLCV-[Pak:Mul:Pep:98] | 1998 | Pakistan | *Capsicum annuum* |
| FM179613 | *Chilli leaf curl virus* | ChiLCV-[Pak:Pot1:Pot:08] | 2008 | Pakistan | *Solanum tuberosum* |
| MN839535 | *Chilli leaf curl virus* | ChiLCV-[Pak:WA19:Pap:18] | 2018 | Pakistan | *Carica papaya* |
| KY420138 | *Chilli leaf curl virus* | ChiLCV-[Pak:SZ-86:Cot:15] | 2015 | Pakistan | *Gossypium herbaceum* |
| KF471061 | *Chilli leaf curl virus* | ChiLCV-[IN:VIRO 385:Amar:15] | 2015 | India | *Amaranthus sp.* |
| EF190217 | *Pepper leaf curl virus* | PepLCV-[IN:Var:Chi:06] | 2006 | India | *Capsicum annuum* |
| MH538340 | *Chilli leaf curl virus* | ChiLCV-[Pak:p-622:Ins-Chi:09] | 2009 | Pakistan | *Insect on Capsicum annuum crop* |
| JN663861 | *Pepper leaf curl varansi virus* | PepLCVV-[IN:Jor:Chi:09] | 2009 | India | *Capsicum annuum* |
| KM023148 | *Chilli leaf curl virus* | ChiLCV-[Pak: CapAS 2:Chi:13] | 2013 | Pakistan | *Capsicum annuum* |
| HM140371 | *Chilli leaf curl virus* | ChiLCV-[IN:Noi:Pap:09] | 2009 | India | *Carica papaya* |
| JN663870 | *Pepper leaf curl virus* | PepLCV-[IN:Pal:Chi:09] | 2009 | India | *Capsicum annuum* |
| HM007104 | *Chilli leaf curl virus* | ChiLCV-[IN:pChJodB2:Chi:09] | 2009 | India | *Capsicum annuum* |
| KX533940 | *Chilli leaf curl virus* | ChiLCV-[IN:ND:chi:16] | 2016 | India | *Capsicum annuum* |
| MN417111 | *Chilli leaf curl virus* | ChiLCV-[IN:CHL45:chi:17] | 2017 | India | *Capsicum annuum* |
| KM023147 | *Chilli leaf curl virus* | ChiLCV-[IN:CapAS 1:chi:13] | 2013 | India | *Capsicum annuum* |
| JN663846 | *Chilli leaf curl virus* | ChiLCV-[IN:Ahm:Xba I-14:chi:09] | 2009 | India | *Capsicum annuum* |
| KF111685 | *Tomato leaf curl Liwa virus* | TLCLWV-[OM:Tom-43:12] | 2012 | Oman | *Solanum tuberosum* |
| KF111684 | *Tomato leaf curl Liwa virus* | TLCLWV-[OM:Tom-42:12] | 2012 | Oman | *Solanum tuberosum* |
| KF111686 | *Tomato leaf curl Liwa virus* | TLCLWV-[OM:Tom-44:12] | 2012 | Oman | *Solanum tuberosum* |
