## Supplemental Table S3 for "Perilous coexistence: *Chilli Leaf Curl Virus* and *Candidatus Phytoplasma trifolii* infecting *Capsicum annuum*, India"

**Table. S3** List of betsatellite collected from GENBANK, NCBI, USA, based on BLAST similarity with our identified ChLCuB-GKP (Acc. No. MZ540909) and organized in listed in order of similarity score

| **Accession No.** | **Species** | **Acronym** | **Year** | **Country** | **Host** |
|---|---|---|---|---|---|
| MZ540909 | *Chilli leaf curl betsatellite* | ChLCuB/[IN:Gkp:Chi:21] | 2021 | India | *Capsicum annuum* |
| MT385295 | *Chilli leaf curl betsatellite* | ChLCuB/[IN:KAN-05:sal spl:20] | 2020 | India | *Salvia splendens* |
| JX193616 | *Chilli leaf curl betsatellite* | ChLCuB/[IN: Meeut:Chi:11] | 2011 | India | *Capsicum annuum* |
| MT385294 | *Chilli leaf curl betsatellite* | ChLCuB/[IN:KAN-03:sal spl:20] | 2020 | India | *Salvia splendens* |
| HM143904 | *Chilli leaf curl betsatellite* | ChLCuB/[IN:Pan4:Pap:08] | 2008 | India | *Carica papaya* |
| HM143901 | *Tomato leaf curl betsatellite* | ToLCB/[IN:Pan1:Pap:08] | 2008 | India | *Carica papaya* |
| HM143905 | *Tomato leaf curl betsatellite* | ToLCB/[IN:Pan5:Pap:08] | 2008 | India | *Carica papaya* |
| KM880104 | *Tomato leaf curl Bangladesh betasatellite* | ToLCBDB/[IN:Ahm:Chi:14] | 2014 | India | *Capsicum annuum* |
| HM143911 | *Tomato leaf curl betsatellite* | ToLCB/[IN:Naj 2:Pap:08] | 2008 | India | *Carica papaya* |
| HM143910 | *Tomato leaf curl betsatellite* | ToLCB/[IN:DU:Pap:09] | 2009 | India | *Carica papaya* |
| MH577024 | *Tomato leaf curl Bangladesh betasatellite* | ToLCBDB/[Ban:pTAJa30:Tom:16] | 2016 | Bangladesh | *Solanum lycopersicum* |
| MT316408 | *Chilli leaf curl betsatellite* | ChLCuB/[Ban:Bagerhat:Chi:19] | 2019 | Bangladesh | *Capsicum annuum L.* |
| MF155644 | *Tomato leaf curl betsatellite* | ToLCB/[IN:ND:Chi:17] | 2017 | India | *Capsicum annuum* |
| MH577022 | *Tomato leaf curl Bangladesh betasatellite* | ToLCBDB/[IN:pTAM1:Tom:09] | 2009 | India | *Solanum lycopersicum* |
| MK087125 | *Tomato leaf curl Bangladesh betasatellite* | ToLCBDB/[IN:FB1:FB:08] | 2008 | India | *Phaseolus vulgaris* |
| HM007105 | *Tomato leaf curl Bangladesh betasatellite* | ToLCBDB/[IN:pChJodBK7:Chi:09] | 2009 | India | *Capsicum annuum* |
| JN663876 | *Tomato leaf curl Bangladesh betasatellite* | ToLCBDB/[IN:Chi:08] | 2008 | India | *Capsicum annuum* |
| JN663875 | *Tomato leaf curl Bangladesh betasatellite* | ToLCBDB/[IN:KpnI-6:FB:Chi:08] | 2008 | India | *Capsicum annuum* |
| KJ868822 | *Tomato leaf curl Bangladesh betasatellite* | ToLCBDB/[IN:Gonda:Chi:13] | 2013 | India | *Capsicum annuum* |
| JN663847 | *Tomato leaf curl Bangladesh betasatellite* | ToLCBDB/[IN:KpnI-5:Chi:09] | 2009 | India | *Capsicum annuum* |
| JN663860 | *Tomato leaf curl Bangladesh betasatellite* | ToLCBDB/[IN:KpnI-1:Chi:08] | 2008 | India | *Capsicum annuum* |
| DQ343289 | *Chilli leaf curl betsatellite* | ChLCuB/[IN:Lko:05] | 2005 | India | --------------- |
| MH577023 | *Tomato leaf curl Bangladesh betasatellite* | ToLCBDB/[IN:pBamP25:Tom:16] | 2016 | India | *Solanum lycopersicum* |
| MT316407 | *Chilli leaf curl betsatellite* | ChLCuB/[Ban:Khulna:Chi:19] | 2019 | Bangladesh | *Capsicum annuum* |
| JF706231 | *Chilli leaf curl betsatellite* | ChLCuB/[IN:Jod:Chi:04] | 2004 | India | *Capsicum annuum* |
| JN663849 | *Papaya leaf curl betsatellite* | PaLCB/[IN: KpnI-3:Chi:08] | 2008 | India | *Capsicum annuum* |
| MH355642 | *Chilli leaf curl betsatellite* | ChLCuB/[IN:CDB1:AD:18] | 2018 | India | *African daisy* |
| JN663854 | *Tomato leaf curl Bangladesh betasatellite* | ToLCBDB/[IN:BamH I-2:Chi:10] | 2010 | India | *Capsicum annuum* |
| JN663856 | *Tomato leaf curl Bangladesh betasatellite* | ToLCBDB/[IN:KpnI 7:Chi:08] | 2008 | India | *Capsicum annuum* |

| | | | | | |
|---|---|---|---|---|---|
| MH577019 | *Tomato leaf curl Bangladesh betasatellite* | ToLCBDB/[IN:pTAsi21:Tom:16] | 2016 | India | *Solanum lycopersicum* |
| HM143902 | *Tomato leaf curl betasatellite* | ToLCB/[IN:Pan2:Pap:08] | 2008 | India | *Carica papaya* |
| MH577021 | *Tomato leaf curl Bangladesh betasatellite* | ToLCBDB/[IN:pBamU13:Tom:16] | 2016 | India | *Solanum lycopersicum* |
| JN663855 | *Tomato leaf curl Bangladesh betasatellite* | ToLCBDB/[IN:KpnI-6:Chi:10] | 2010 | India | *Capsicum annuum* |
| MT385299 | *Chilli leaf curl betasatellite* | ChLCuB/[IN: LKO-08:Sal spl:20] | 2020 | India | *Salvia splendens* |
| JN663869 | *Papaya leaf curl betasatellite* | PaLCB/[IN:KpnI-4:Chi:10] | 2010 | India | *Capsicum annuum* |
| JN663868 | *Papaya leaf curl betasatellite* | PaLCB/[IN:KpnI-3:Chi:10] | 2010 | India | *Capsicum annuum* |
| JQ654464 | *Tomato leaf curl Bangladesh betasatellite* | ToLCBDB/[IN:HJP01:MB:11] | 2011 | India | *Vigna radiata* |
| HM007118 | *Tomato leaf curl Bangladesh betasatellite* | ToLCBDB/[IN:pChPatnBK19:Chi:08] | 2008 | India | *Capsicum annuum* |
| LT827057 | *Tomato leaf curl Bangladesh betasatellite* | ToLCBDB/[IN:IS-12:Weed:16] | 2016 | India | *Weed* |
| KR957354 | *Tomato leaf curl Bangladesh betasatellite* | ToLCBDB/[IN:VIRO 765:Chi:09] | 2009 | India | *Capsicum annuum* |
| MT861129 | *Tomato leaf curl Bangladesh betasatellite* | ToLCBDB/[IN:SPUR1:BP:15] | 2015 | India | *Capsicum annuum* |
| EU582020 | *Chilli leaf curl betasatellite* | ChLCuB/[IN:Pataudi:Chi:08] | 2008 | India | *Capsicum annuum* |
| KJ700655 | *Chilli leaf curl betasatellite* | ChLCuB/[IN:RKB2:Pet:14] | 2014 | India | *Petunia* |
