## Supplemental Table S4 for "Perilous coexistence: *Chilli Leaf Curl Virus* and *Candidatus Phytoplasma trifolii* infecting *Capsicum annuum*, India"

**Table. S4** Amino acid pairwise % identities between ChiLCV-A-GKP (Acc. No. MZ540908) and all six gene components of identified begomoviruses with other similar isolates by use Sequence Demarcation Tool (SDT v1.2) (Muhire et al., 2014).

| Accession number | Begomovirus | Genome | Gene (percentage amino acid sequence identity) | | | | | |
|---|---|---|---|---|---|---|---|---|
| | | | Pre-CP | CP | REP | TrAP | REn | AC4 |
| MZ540908 | ChiLCV-[IN:GKP:Chi:21] | 100.00 | 100.00 | 100.00 | 100.00 | 100.00 | 100.00 | 100.00 |
| MK757213 | ChiLCV-[OM:Tom30-19:16] | 87.801 | 93.901 | 93.001 | 95.001 | 95.301 | 96.301 | 88.701 |
| MK757215 | ChiLCV-[OM:Tom32-20:16] | 87.801 | 93.801 | 92.601 | 94.701 | 95.301 | 96.301 | 88.701 |
| MK757214 | ChiLCV-[OM:Tom31-23:16] | 87.801 | 93.801 | 93.001 | 95.001 | 95.301 | 96.301 | 88.701 |
| MK757212 | ChiLCV-[OM:Tom26-12:16] | 87.701 | 93.801 | 93.001 | 95.001 | 95.301 | 96.301 | 88.701 |
| MN119491 | ChiLCV-[OM:P5:Pum:16] | 87.601 | 93.701 | 92.201 | 95.001 | 94.501 | 96.301 | 88.701 |
| HG969265 | ChiLCV-[OM:C1:13] | 87.601 | 90.201 | 92.201 | 94.701 | 95.301 | 94.801 | 88.701 |
| MK468696 | ChiLCV-[OM:Tob-13:15] | 87.601 | 93.901 | 93.001 | 95.001 | 94.501 | 94.801 | 88.701 |
| HG969264 | ChiLCV-[OM:Rad-4:13] | 87.601 | 93.701 | 92.601 | 95.001 | 94.501 | 95.501 | 88.701 |
| MN119492 | ChiLCV-[OM:P6:Pum:16] | 87.601 | 93.901 | 93.001 | 95.001 | 94.501 | 95.501 | 88.701 |
| HE806437 | ChiLCV-[OM:T-37:11] | 87.601 | 93.901 | 93.001 | 95.301 | 95.301 | 94.801 | 88.701 |
| MK468694 | ChiLCV-[OM:Tob-11:15] | 87.601 | 93.901 | 93.001 | 95.001 | 95.301 | 94.801 | 88.701 |
| KF229718 | ChiLCV-[OM:Tom-26:12] | 87.501 | 93.901 | 93.001 | 95.301 | 94.501 | 95.501 | 88.701 |
| HG969263 | ChiLCV-[OM:Rad-3:13] | 87.401 | 93.801 | 91.801 | 95.001 | 95.301 | 95.501 | 88.701 |

| | | | | | | | | |
|---|---|---|---|---|---|---|---|---|
| HF968755 | ChiLCV-[OM:PT-2:12] | 87.301 | 93.801 | 92.601 | 94.701 | 95.301 | 95.501 | 88.701 |
| LN680633 | ChiLCV-[OM:Tom112:14] | 86.701 | 93.801 | 92.201 | 95.001 | 94.501 | 94.801 | 88.701 |
| HF968756 | ChiLCV-[OM:PT-3:12] | 86.301 | 93.801 | 92.601 | 95.401 | 91.401 | 91.801 | 88.701 |
| MF737343 | ChiLCV-[IN:MM1:17] | 86.301 | 95.201 | 94.201 | 92.201 | 95.301 | 91.801 | 75.301 |
| HG969197 | ChiLCV-[OM:Tom84:14] | 84.101 | 90.101 | 92.601 | 94.701 | 95.301 | 94.801 | 88.701 |
| LN680625 | ChiLCV-[OM:Tom85:14] | 84.001 | 90.201 | 92.601 | 95.001 | 95.301 | 94.801 | 88.701 |
| HG969257 | ChiLCV-[OM:HP-12:13] | 84.001 | 90.201 | 92.601 | 95.001 | 95.301 | 94.001 | 88.701 |
| LN680626 | ChiLCV-[OM:Tom88:14] | 83.901 | 90.201 | 92.601 | 95.001 | 95.301 | 94.001 | 88.701 |
| LN680624 | ChiLCV-[OM:Tom87:14] | 83.901 | 92.901 | 92.601 | 95.001 | 95.301 | 94.001 | 88.701 |
| HG969255 | ChiLCV-[OM:HP-5:13] | 83.801 | 90.201 | 92.601 | 95.001 | 91.401 | 91.801 | 88.701 |
| JN604491 | ChiLCV-[OM:Sh1:Pep:11] | 81.101 | 83.601 | 83.601 | 95.801 | 96.101 | 94.001 | 88.701 |
| JN604493 | ChiLCV-[OM:Taq2:Tom:11] | 80.801 | 84.401 | 84.401 | 94.701 | 95.301 | 20.701 | 87.601 |
| AF336806 | ChiLCV-[Pak:Mul:Pep:98] | 77.001 | 84.801 | 84.401 | 95.701 | 96.101 | 89.601 | 85.601 |
| MN839535 | ChiLCV-[Pak:WA19:Pap:18] | 76.801 | 84.801 | 84.801 | 95.101 | 96.901 | 87.301 | 84.501 |
| FM179613 | ChiLCV-[Pak:Pot1:Pot:08] | 76.301 | 84.801 | 84.801 | 95.701 | 96.101 | 89.601 | 85.601 |
| KF111685 | TLCLWV-[OM:Tom-43:12] | 75.301 | 93.401 | 93.001 | 77.301 | 94.501 | 94.801 | 21.801 |
| KF111684 | TLCLWV-[OM:Tom-42:12] | 74.901 | 93.701 | 92.601 | 77.301 | 94.501 | 94.001 | 38.801 |
| KF111686 | TLCLWV-[OM:Tom-44:12] | 74.101 | 93.801 | 92.601 | 76.501 | 88.301 | 88.101 | 35.301 |
| KM023147 | ChiLCV-[IN:CapAS 1:chi:13] | 74.201 | 84.801 | 84.801 | 95.001 | 100.001 | 84.301 | 93.801 |
| HM140371 | ChiLCV-[IN:Noi:Pap:09] | 73.901 | 84.801 | 84.801 | 95.001 | 100.001 | 84.301 | 93.801 |
| MN417111 | ChiLCV-[IN:CHL45:chi:17] | 73.201 | 84.401 | 84.401 | 95.601 | 97.701 | 84.301 | 87.601 |

| | | | | | | | | |
|---|---|---|---|---|---|---|---|---|
| JN663846 | ChiLCV-[IN:Ahm:Xba I-14:chi:09] | 72.701 | 85.201 | 85.201 | 94.201 | 96.101 | 83.601 | 91.801 |
| MN119490 | ChiLCV-[OM:P14:Pum:16] | 25.101 | 93.901 | 93.001 | 95.001 | 95.301 | 96.301 | 88.701 |
| KX787939 | ChiLCV-[OM:Wat-122:15] | 25.301 | 93.901 | 93.001 | 94.701 | 95.301 | 96.301 | 80.401 |
| JN663861 | PepLCVV-[IN:Jor:Chi:09] | 25.201 | 20.801 | 19.601 | 94.701 | 93.801 | 87.301 | 87.601 |
| EF190217 | PepLCV-[IN:Var:Chi:06] | 24.601 | 84.801 | 84.801 | 94.701 | 93.801 | 84.301 | 87.601 |
| MH538340 | ChiLCV-[Pak:p-622:Ins-Chi:09] | 24.601 | 84.001 | 84.001 | 95.001 | 97.701 | 84.301 | 82.501 |
| JN663870 | PepLCV-[IN:Pal:Chi:09] | 23.901 | 84.001 | 84.001 | 94.201 | 93.801 | 84.301 | 84.501 |
| KF471061 | ChiLCV-[IN:VIRO 385:Amar:15] | 22.901 | 18.901 | 18.901 | 94.701 | 93.001 | 84.301 | 85.601 |
| HM007104 | ChiLCV-[IN:pChJodB2:Chi:09] | 22.701 | 85.201 | 85.201 | 95.001 | 97.701 | 84.301 | 91.801 |
| KX533940 | ChiLCV-[IN:ND:chi:16] | 22.101 | 85.201 | 85.201 | 94.701 | 97.701 | 85.101 | 90.701 |
| KM023148 | ChiLCV-[Pak: CapAS 2:Chi:13] | 21.801 | 84.801 | 84.801 | 95.001 | 100.001 | 84.301 | 93.801 |
| KY420138 | ChiLCV-[Pak:SZ-86:Cot:15] | 21.901 | 82.201 | 78.801 | 94.501 | 96.101 | 84.301 | 82.501 |
| EU939533 | ChiLCV-[IN:Nar:Pep:04] | 21.601 | 84.401 | 84.401 | 95.601 | 96.901 | 85.801 | 82.501 |
| JN604490 | ChiLCV-[OM:Sal2:Pep:11] | 20.101 | 83.701 | 83.701 | 95.301 | 95.301 | 94.801 | 88.701 |
| JN604500 | ChiLCV-[OM:KW1:Tom:11] | 18.726 | 82.401 | 82.401 | 95.301 | 95.301 | 94.801 | 88.701 |
| JN604494 | ChiLCV_OM_Taq3_Tom_11 | 19.901 | 84.401 | 84.401 | 95.301 | 94.501 | 94.001 | 87.601 |
| JN604496 | ChiLCV-[OM:Th1:Tom:11] | 19.901 | 84.401 | 84.401 | 94.701 | 95.301 | 94.801 | 87.601 |
| MH475359 | ChiLCV-[OM:Wed8-25:UI:16] | 19.801 | 84.401 | 84.401 | 95.601 | 95.301 | 93.301 | 87.601 |
| JN604495 | ChiLCV-[OM:Taq4:Tom11] | 19.801 | 84.401 | 84.401 | 95.301 | 94.501 | 94.001 | 88.701 |

| | | | | | | | | |
|---|---|---|---|---|---|---|---|---|
| HG941644 | ChiLCV-[OM:TB-23:Tob:13] | 19.601 | 84.401 | 84.401 | 95.001 | 94.501 | 93.301 | 88.701 |
| HG941643 | ChiLCV-[OM:BZ-8:13] | 19.501 | 84.401 | 84.401 | 94.501 | 94.501 | 94.001 | 88.701 |
| JN604498 | ChiLCV-[OM:Th3:Tom:11] | 19.401 | 84.401 | 84.401 | 95.001 | 95.301 | 94.801 | 88.701 |
| MH475358 | ChiLCV-[OM:Wed8-24:UI:16] | 19.401 | 84.401 | 84.401 | 95.401 | 94.501 | 92.501 | 87.601 |
| JN604497 | ChiLCV-[OM:Th2:Tom:11] | 19.401 | 84.401 | 84.401 | 94.501 | 95.301 | 20.701 | 88.701 |
| JN604489 | ChiLCV-[OM:Sal1:Pep:11] | 19.001 | 83.301 | 83.301 | 94.701 | 94.501 | 94.001 | 87.601 |
| JN604499 | ChiLCV-[OM:Th4:Tom:11] | 18.901 | 84.401 | 84.401 | 95.001 | 95.301 | 94.801 | 88.701 |
