## Supplemental Table S5 for "Perilous coexistence: *Chilli Leaf Curl Virus* and *Candidatus Phytoplasma trifolii* infecting *Capsicum annuum*, India"

**Table. S5** Amino acid pairwise percentage identities between ChLCuB-GKP (Acc. No. MZ540909) and βC1 gene components of identified begomoviruses with other similar isolates by use Sequence Demarcation Tool (SDT v1.2) (Muhire et al., 2014).

| **Accession Number** | **Begomovirus** | **Betasatellite** | **Gene (percentage amino acid sequence identity)** |
|---|---|---|---|
| | | | **βC1** |
| MZ540909 | ChLCuB/[IN:Gkp:Chi:21] | 100.001 | 100.001 |
| DQ343289 | ChLCuB/[IN:Lko:05] | 50.601 | 85.801 |
| JF706231 | ChLCuB/[IN:Jod:Chi:04] | 48.001 | 88.301 |
| HM007105 | ToLCBDB/[IN:pChJodBK7:Chi:09] | 47.801 | 20.501 |
| MT316408 | ChLCuB/[Ban:Bagerhat:Chi:19] | 47.201 | 87.501 |
| MT385299 | ChLCuB/[IN: LKO-08:Sal spl:20] | 47.201 | 88.301 |
| JN663854 | ToLCBDB/[IN:BamH I-2:Chi:10] | 46.401 | 87.501 |
| MH577021 | ToLCBDB/[IN:pBamU13:Tom:16] | 45.901 | 85.801 |
| KJ700655 | ChLCuB/[IN:RKB2:Pet:14] | 45.701 | 85.801 |
| KJ868822 | ToLCBDB/[IN:Gonda:Chi:13] | 44.601 | 86.701 |
| MT316407 | ChLCuB/[Ban:Khulna:Chi:19] | 44.501 | 87.501 |
| EU582020 | ChLCuB/[IN:Pataudi:Chi:08] | 44.401 | 87.501 |
| JQ654464 | ToLCBDB/[IN:HJP01:MB:11] | 43.201 | 89.201 |
| MH577019 | ToLCBDB/[IN:pTAsi21:Tom:16] | 42.301 | 85.001 |
| JN663855 | ToLCBDB/[IN:KpnI-6:Chi:10] | 42.101 | 88.301 |
| MF155644 | ToLCB/[IN:ND:Chi:17] | 40.201 | 85.601 |
| KR957354 | ToLCBDB/[IN:VIRO 765:Chi:09] | 38.401 | 85.001 |
| KM880104 | ToLCBDB/[IN:Ahm:Chi:14] | 37.301 | 90.801 |
| MK087125 | ToLCBDB/[IN:FB1:FB:08] | 36.301 | 91.701 |
| MH577022 | ToLCBDB/[IN:pTAM1:Tom:09] | 34.801 | 92.501 |
| HM143905 | ToLCB/[IN:Pan5:Pap:08] | 32.901 | 89.801 |
| JX193616 | ChLCuB/[IN: Meeut:Chi:11] | 32.701 | 95.801 |
| MT385294 | ChLCuB/[IN:KAN-03:sal spl:20] | 31.901 | 94.601 |
| JN663847 | ToLCBDB/[IN:KpnI-5:Chi:09] | 31.601 | 86.701 |
| MH577023 | ToLCBDB/[IN:pBamP25:Tom:16] | 31.101 | 90.801 |
| MT385295 | ChLCuB/[IN:KAN-05:sal spl:20] | 30.401 | 100.001 |

| | | | |
|---|---|---|---|
| HM143910 | ChLCuB/[IN:KAN-05:sal spl:20] | 30.301 | 84.701 |
| HM143911 | ToLCB/[IN:Naj 2:Pap:08] | 30.201 | 85.601 |
| JN663875 | ToLCBDB/[IN:KpnI-6:FB:Chi:08] | 30.001 | 87.501 |
| JN663869 | PaLCB/[IN:KpnI-4:Chi:10] | 27.801 | 82.501 |
| JN663868 | PaLCB/[IN:KpnI-3:Chi:10] | 27.501 | 82.501 |
| MH577024 | ToLCBDB/[Ban:pTAJa30:Tom:16] | 27.301 | 89.801 |
| MH355642 | ChLCuB/[IN:CDB1:AD:18] | 25.201 | 88.301 |
| JN663856 | ToLCBDB/[IN:KpnI 7:Chi:08] | 24.001 | 84.201 |
| HM143902 | ToLCB/[IN:Pan2:Pap:08] | 23.101 | 83.901 |
| JN663860 | ToLCBDB/[IN:KpnI-1:Chi:08] | 23.301 | 20.501 |
| JN663849 | PaLCB/[IN: KpnI-3:Chi:08] | 22.701 | 86.701 |
| HM143901 | ToLCB/[IN:Pan1:Pap:08] | 21.801 | 91.501 |
| HM007118 | ToLCBDB/[IN:pChPatnBK19:Chi:08] | 21.601 | 85.801 |
| MT861129 | ToLCBDB/[IN:SPUR1:BP:15] | 21.501 | 89.201 |
| JN663876 | ToLCBDB/[IN:Chi:08] | 20.901 | 88.301 |
| HM143904 | ChLCuB/[IN:Pan4:Pap:08] | 20.201 | 87.301 |
| LT827057 | ToLCBDB/[IN:IS-12:Weed:16] | 18.201 | 86.701 |
