## Supplemental Table S6 for "Perilous coexistence: *Chilli Leaf Curl Virus* and *Candidatus Phytoplasma trifolii* infecting *Capsicum annuum*, India"

**Table. S6** List of 16S rRNA isolates collected from GENBANK, NCBI, USA, and USA based on BLAST similarity and organised in listed in order of similarity score.

| Accession number | Species | ACRONYMS | Year | Country | Host |
|---|---|---|---|---|---|
| MZ557805 | *Candidatus Phytoplasma trifolii* | Ca.P.trifolii/16VI-A:21 | 2021 | India | *Capsicum annuum* |
| MN861370 | *Candidatus Phytoplasma trifolii* | Ca.P.trifolii/HSD2/16VI-A:19 | 2019 | India | *Helichrysum sp.* |
| MN861357 | *Candidatus Phytoplasma trifolii* | Ca.P.trifolii/HSD1/16VI-A:19 | 2019 | India | *Helichrysum sp* |
| KX284698 | *Brinjal little leaf phytoplasma* | Brinjal little leaf/LLB_MKS2/16VI-D:14 | 2014 | India | *Solanum melongena L* |
| KX671554 | *Lectuca sativa' flattened stem phytoplasma* | 'Lectuca sativa' flattened stem/NDL:14 | 2014 | India | *Lectuca sativa* |
| KP027531 | *Catharanthus phyllody phytoplasma* | Catharanthus phyllody /KA-02:14 | 2014 | India | *Catharanthus roseus* |
| KP027517 | *Brinjal little leaf phytoplasma* | Brinjal little leaf/TN-34/16VI-D:14 | 2014 | India | *Solanum melongena* |
| KP027513 | *Brinjal little leaf phytoplasma* | Brinjal little leaf/AP-28/16VI-D :14 | 2014 | India | *Solanum melongena* |
| KP027512 | *Brinjal little leaf phytoplasma* | Brinjal little leaf/AP-26/16VI-D:14 | 2014 | India | *Solanum melongena* |
| KP027510 | *Brinjal little leaf phytoplasma* | Brinjal little leaf/AP-24/16VI-D:14 | 2014 | India | *Solanum melongena* |
| KP027508 | *Brinjal little leaf phytoplasma* | Brinjal little leaf/AP-22/16VI-D:14 | 2014 | India | *Solanum melongena* |
| KP027505 | *Brinjal little leaf phytoplasma* | Brinjal little leaf/AP-19/16VI-D:14 | 2014 | India | *Solanum melongena* |
| KP027504 | *Brinjal little leaf phytoplasma* | Brinjal little leaf/AP-18/16VI-D:14 | 2014 | India | *Solanum melongena* |
| KP027503 | *Brinjal little leaf phytoplasma* | Brinjal little leaf/AP-17/16VI-D:14 | 2014 | India | *Solanum melongena* |
| KP027501 | *Brinjal little leaf phytoplasma* | Brinjal little leaf/AP-14/16VI-D:14 | 2014 | India | *Solanum melongena* |
| KP027500 | *Brinjal little leaf phytoplasma* | Brinjal little leaf/AP-13/16VI-D:14 | 2014 | India | *Solanum melongena* |
| MZ773477 | *Candidatus Phytoplasma trifolii* | Ca.P trifolii/THS1/16VI-A:21 | 2021 | India | *Roselle* |
| MZ425930 | *Brinjal little leaf phytoplasma* | Brinjal little leaf/T-M2/16VI-D :21 | 2021 | India | *Solanum melongena* |
| MZ425929 | *Brinjal little leaf phytoplasma* | Brinjal little leaf/T-M1/16VI-D:21 | 2021 | India | *Solanum melongena* |
| JX104336 | *Candidatus Phytoplasma trifolii* | Ca.P trifolii/BRGL-2/16VI-A:12 | 2012 | India | *Solanum melongena* |
| MW648598 | *Hishimonus phycitis' leafhopper phytoplasma* | Hishimonus phycitis leafhopper/HpLH-LKO2:21 | 2021 | India | *Hishimonus phycitis* |
| MW648597 | *Hishimonus phycitis' leafhopper phytoplasma* | Hishimonus phycitis leafhopper/HpLH-LKO1:21 | 2021 | India | *Hishimonus phycitis* |
| MW648596 | *Catharanthus roseus phytoplasma* | Catharanthus roseus phytoplasma/Cr-LKO2:21 | 2021 | India | *Catharanthus roseus* |
| MW648595 | *Catharanthus roseus phytoplasma* | Catharanthus roseus /Cr-LK /O1:21 | 2021 | India | *Catharanthus roseus* |
| MW648594 | *Chrysanthemum coronarium' stunting phytoplasma* | Chrysanthemum coronarium/ChS-LKO2:21 | 2021 | India | *Glebionis coronaria* |
| MW648593 | *Chrysanthemum coronarium' stunting phytoplasma* | Chrysanthemum coronarium/ChS-LKO1:21 | 2021 | India | *Glebionis coronaria* |
| KC178679 | *Brinjal little leaf phytoplasma* | Brinjal little leaf/B-NDL/16VI-D:12 | 2012 | India | *Solanum melongena* |
| MW261866 | *Candidatus Phytoplasma trifolii* | Ca.P. trifolii/Nagaon2/16VI-A:20 | 2020 | India | *Solanum melongena* |
| MW261863 | *Candidatus Phytoplasma trifolii* | Ca.P. trifolii/Nagaon1//16VI-A:20 | 2020 | India | *Solanum violaceum* |
| MT739373 | *Brinjal little leaf phytoplasma* | Brinjal little leaf/BLL//16VI-D:20 | 2020 | India | *Solanum melongena* |
| EF186820 | *Brinjal little leaf phytoplasma* | Brinjal little leaf/BLL//16VI-D:06 | 2006 | U.S.A | ---------------------- |
| KX284703 | *Brinjal little leaf phytoplasma* | Brinjal little leaf/LLB_MKS//16VI-D:16 | 2016 | India | *Solanum melongena L.* |

| | | | | | |
|---|---|---|---|---|---|
| KX284699 | *Brinjal little leaf phytoplasma* | Brinjal little leaf/LLB_MKS3/16VI-D:16 | 2016 | India | *Solanum melongena L.* |
| KP027521 | *Brinjal little leaf phytoplasma* | Brinjal little leaf/TN-38//16VI-D:14 | 2014 | India | *Solanum melongena* |
| MF996472 | *Candidatus Phytoplasma trifolii* | Ca.P. trifolii/NeWB1/16VI-A:17 | 2017 | India | *neem* |
| KX284702 | *Brinjal little leaf phytoplasma* | Brinjal little leaf/LLB_MKS6/16VI-A:16 | 2016 | India | *Solanum melongena L.* |
| KX284701 | *Brinjal little leaf phytoplasma* | Brinjal little leaf/LLB_MKS5/16VI-A:16 | 2016 | India | *Solanum melongena L.* |
| KX284697 | *Brinjal little leaf phytoplasma* | Brinjal little leaf/LLB_MKS1/16VI-A:16 | 2016 | India | *Solanum melongena L.* |
| KP027495 | *Brinjal little leaf phytoplasma* | Brinjal little leaf/KA-04/16VI-A:14 | 2014 | India | *Solanum melongena* |
| AF228053 | *Periwinkle little leaf phytoplasma* | Periwinkle little leaf/16S:00 | 2000 | Bangladesh | *Catharanthus roseus* |
| AF228052 | *Brinjal little leaf phytoplasma* | Brinjal little leaf/16VI-A:00 | 2000 | Bangladesh | *Solanum melongena* |
| MW261864 | *Candidatus Phytoplasma trifolii* | Ca.P trifolii/Sonitpur1/16VI-A:14 | 2020 | India | *Datura sp.* |
| KP027506 | *Brinjal little leaf phytoplasma* | Brinjal little leaf/AP-20/16VI-A:16 | 2014 | India | *Solanum melongena* |
| KX284700 | *Brinjal little leaf phytoplasma* | Brinjal little leaf/LLB_MKS4/16VI-A:16 | 2016 | India | *Solanum melongena L.* |
| X83431 | *Mollicutes sp.* | Mollicutes sp:14 | 1994 | Germeny | *Solanum melongena* |
| KP027524 | *Brinjal little leaf phytoplasma* | Brinjal little leaf/TN-42/16VI-A:14 | 2014 | India | *Solanum melongena* |
| KP027522 | *Brinjal little leaf phytoplasma* | Brinjal little leaf/TN-39/16VI-A:14 | 2014 | India | *Solanum melongena* |
| KP027514 | *Brinjal little leaf phytoplasma* | Brinjal little leaf/AP-31/16VI-A:07 | 2014 | India | *Solanum melongena* |
| EF534205 | *Iranian safflower phyllody phytoplasma* | Iranian safflower phyllody:07 | 2007 | Iran | ---------------------- |
| MW847614 | *Candidatus Phytoplasma trifolii* | Ca.P trifolii/SV1/16VI-A:21 | 2021 | India | *Solanum violaceum* |
| HQ609491 | *Potato witches'-broom phytoplasma* | Potato witches'-broom/16VI-A/AK-DN1L:10 | 2010 | U.S.A | *Solanum tuberosum* |
| HQ609489 | *Potato witches'-broom phytoplasma* | Potato witches'-broom /CN-99:10 | 2010 | China | *Solanum tuberosum* |
| GU004368 | *Potato purple top phytoplasma PPT-AK4* | Potato purple top/PPT-AK4 strain AKpot4:09 | 2009 | U.S.A | *Solanum tuberosum* |
| EU346380 | *Candidatus Phytoplasma trifolii* | Ca.P. trifolii/AK-1/16VI-A:16 | 2016 | India | *Solanum melongena L.* |
| AY390261 | *Candidatus Phytoplasma trifolii* | Ca.P. trifolii//16VI-A:03 | 2003 | Canada | *Trifolium hybridum* |
| KR072667 | *Alaska potato witches'-broom phytoplasma* | Alaska potato witches'-broom/AK-5:15 | 2015 | U.S.A | *Solanum tuberosum* |
| GU004367 | *Potato purple top phytoplasma PPT-AK4* | Potato purple top/PPT-AK2:09 | 2009 | U.S.A | *Solanum tuberosum* |
| GU004366 | *Potato purple top phytoplasma PPT-AK4* | Potato purple top/PPT-AK1:09 | 2009 | U.S.A | *Solanum tuberosum* |
| EU543441 | *Candidatus Phytoplasma trifolii* | Ca.P. trifolii/16VI-A:08 | 2008 | Czech Republic | *hybrid Rhododendron cultivar Don Juan* |
| EU543440 | *Candidatus Phytoplasma trifolii* | Ca.P. trifolii/16VI-A:08 | 2008 | Czech Republic | *hybrid Rhododendron cultivar Don Juan* |
| EU143330 | *'Vaccinium myrtillus' phytoplasma* | 'Vaccinium myrtillus':07 | 2007 | Austria | *Vaccinium myrtillus* |
| AY270156 | *Centaurea solstitialis virescence phytoplasma* | Centaurea solstitialis virescence:03 | 2003 | Italy | *Centaurea solstitialis* |
| EF186821 | *Lucerne virescence phytoplasma* | Lucerne virescence:06 | 2006 | France | ---------------------- |
| MF385584 | *Candidatus Phytoplasma trifolii* | Ca.P.trifolii/Delaware/16VI-A:17 | 2017 | U.S.A | *Ulmus americana* |
| HQ589189 | *Candidatus Phytoplasma trifolii* | Ca.P.trifolii/CP-1/16VI-A:10 | 2010 | Canada | *Periwinkle* |
| GU004369 | *Potato purple top phytoplasma PPT-AK5* | Potato purple top/PPT-AK5:09 | 2009 | U.S.A | *Solanum tuberosum* |
| AB279597 | *Candidatus Phytoplasma trifolii* | Ca.P.trifolii/16VI-A:06 | 2006 | South Korea | *Lespedeza sp.* |
| AY500818 | *Potato witches'-broom phytoplasma* | Potato witches'-broom:03 | 2003 | Canada | ---------------------- |
| AF409069 | *Candidatus Phytoplasma trifolii* | Ca.P.trifolii/16VI-A:01 | 2001 | U.S.A | ---------------------- |

| | | | | | |
|---|---|---|---|---|---|
| DQ256089 | *Potato witches'-broom phytoplasma* | Potato witches'-broom:05 | 2005 | Canada | *Solanum tuberosum* |
| KX641022 | *Saponaria sp.' stunting and witches' broom phytoplasma* | 'Saponaria sp.'stunting and witches' broom/Pune 4:16 | 2016 | India | *Saponaria sp.* |
| AF409070 | *Candidatus Phytoplasma trifolii* | Ca.P.trifolii/16VI-A:01 | 2001 | U.S.A | ---------------------- |
| AY500130 | *Candidatus Phytoplasma trifolii* | Ca.P. trifolii/16VI-A:03 | 2003 | Canada | ---------------------- |
| AB076404 | *Periwinkle little leaf phytoplasma* | Potato witches'-broom:01 | 2001 | South Korea | ---------------------- |
| MZ413119 | *'Acacia arabica' phytoplasma* | 'Acacia arabica' /BaSD2:21 | 2021 | India | *Acacia arabica* |
| MZ413118 | *'Acacia arabica' phytoplasma* | 'Acacia arabica' /BaSD1:21 | 2021 | India | *Acacia arabica* |
| MF509775 | *'Cannabis sativa' phytoplasma* | 'Cannabis sativa'/NDL :17 | 2017 | India | *Cannabis sativa* |
| MK693146 | *Candidatus Phytoplasma trifolii* | Ca.P.trifolii/IARI-2/16VI-A:19 | 2019 | India | *Gladiolus grandiflorus* |
| MK369688 | *Candidatus Phytoplasma trifolii* | Ca.P. trifolii/IARI-1/16VI-A:19 | 2019 | India | *Gladiolus grandiflorus* |
| KY856746 | *Hishimonus phycitis' 16SrVI phytoplasma* | 'Hishimonas phycitis'/NDL/16SrVI:21 | 2021 | India | *Acacia arabica* |
| KY398725 | *Candidatus Phytoplasma trifolii* | Ca.P. trifolii/PW32/16VI-A:21 | 2021 | India | *Acacia arabica* |
| KX689245 | *Brinjal little leaf phytoplasma* | Brinjal little leaf/CG-A/16VI-D:17 | 2017 | India | *Cannabis sativa* |
| KX689234 | *Brinjal little leaf phytoplasma* | Brinjal little leaf/IARI-A/16VI-D:19 | 2019 | India | *Gladiolus grandiflorus* |
| KR072666 | *Columbia Basin potato purple top phytoplasma* | Columbia Basin potato purple top:19 | 2019 | India | *Gladiolus grandiflorus* |
| OM654559 | *Sesamum indicum' phyllody phytoplasma* | 'Sesamum indicum' /Torogh-Mashhad:22 | 2022 | Iran | *Sesamum indicum* |
| KC478606 | *Brinjal little leaf phytoplasma* | Brinjal little leaf/BLL-VNS-1/16VI-D:13 | 2013 | India | *Solanum melongena* |
| KF178706 | *Candidatus Phytoplasma trifolii* | Ca.P.trifolii/16VI-A:13 | 2013 | USA | ---------------------- |
| JF508517 | *Cucumber phyllody phytoplasma* | Cucumber phyllody/Cuph2:11 | 2011 | Iran | *Cucumis sativus* |
| JF508514 | *Sesame phyllody phytoplasma* | Sesame phyllody/Seph2:11 | 2011 | Iran | *Sesamum indicum* |
| HQ436488 | *Chile pepper phytoplasma* | Chile pepper/NM:10 | 2010 | USA | *Capsicum annuum* |
| AY496005 | *Washington potato purple top phytoplasma* | Washington Potato Purple Top/PPT-Wa8:03 | 2003 | USA | *Solanum tuberosum* |
| MH547069 | *Periwinkle little leaf phytoplasma* | Periwinkle little leaf/7 16S:18 | 2018 | India | *Solanum melongena* |
| EU346381 | *Candidatus Phytoplasma trifolii* | Ca.P.trifolii/AK-2/16VI-A:07 | 2007 | USA | ---------------------- |
| JQ409541 | *Brinjal little leaf phytoplasma* | Brinjal little leaf/16VI-D:12 | 2012 | India | ---------------------- |
| EF592606 | *Iranian cabbage yellows phytoplasma* | Iranian cabbage yellows:07 | 2007 | Iran | ---------------------- |
| MK660149 | *Brassica juncea' phyllody phytoplasma* | Brassica juncea' phyllody /OM2:19 | 2019 | China | *Brassica juncea* |
| MK660148 | *Brassica juncea' phyllody phytoplasma* | Brassica juncea' phyllody/OM1:19 | 2019 | China | *Brassica juncea* |
| KP119494 | *'Brassica juncea' phyllody phytoplasma* | Cucurbita pepo' /Neyshabur:16 | 2016 | Iran | *Cucurbita pepo* |
| MT981209 | *Candidatus Phytoplasma trifolii* | Ca.P. trifolii/AP/16VI-A:20 | 2020 | Iran | *Malus sp.* |
| FJ525437 | *Chile pepper phytoplasma* | Chile pepper/New Mexico:08 | 2008 | USA | ---------------------- |
