## Supplemental Table S7 for "Perilous coexistence: *Chilli Leaf Curl Virus* and *Candidatus Phytoplasma trifolii* infecting *Capsicum annuum*, India"

**Table. S7** List of Sec-A isolates collected from GENBANK, NCBI, USA, and USA based on BLAST similarity and organised in listed in order of similarity score

| Accession Number | Species | Acronyms | Year | Country | Host |
|---|---|---|---|---|---|
| MZ620707 | *Candidatus Phytoplasma trifolii* | Ca.P.trifolii/secA:21 | 2021 | India | *Capsicum annuum* |
| KR906730 | *Brinjal little leaf phytoplasma* | Brinjal little leaf/secA:15 | 2015 | India | *Solanum melongena* |
| KR906727 | *Brinjal little leaf phytoplasma* | Brinjal little leaf/secA:15 | 2015 | India | *Solanum melongena* |
| KR906724 | *Brinjal little leaf phytoplasma* | Brinjal little leaf/IARI-4/secA:15 | 2015 | India | *Solanum melongena* |
| KR906716 | *Brinjal little leaf phytoplasma* | Brinjal little leaf/NOIDA-1/secA:15 | 2015 | India | *Solanum melongena* |
| KR906729 | *Cannabis sativa' phytoplasma* | Cannabis sativa/NOIDA/secA:15 | 2015 | India | *Cannabis sativa* |
| KX784498 | *Candidatus Phytoplasma trifolii* | Ca.P.trifolii/VP-03/SecA:15 | 2015 | India | *Cannabis sativa* |
| KX894792 | *Candidatus Phytoplasma trifolii* | Ca.P.trifolii/VP 05/SecA:15 | 2016 | India | ------------- |
| KY064175 | *Candidatus Phytoplasma trifolii* | Ca.P.trifolii/MKS 04/Sec A:16 | 2016 | India | ------------- |
| KX857660 | *Candidatus Phytoplasma trifolii* | Ca.P.trifolii/VP 04/SecA:16 | 2016 | India | ------------- |
| KR906718 | *Brinjal little leaf phytoplasma* | Brinjal little leaf/Odisha-2/secA:16 | 2015 | India | *Solanum melongena* |
| KY228385 | *Candidatus Phytoplasma trifolii* | Ca.P.trifolii/MKS 06:15 | 2016 | India | *Solanum melongena* |
| KY073129 | *Candidatus Phytoplasma trifolii* | Ca.P.trifolii/MKS 07/SecA:16 | 2016 | India | ------------- |
| EU168743 | *Brinjal little leaf phytoplasma* | Brinjal little leaf/secA:16 | 2007 | UK | ------------- |
| KX622584 | *Candidatus Phytoplasma trifolii* | Ca.P.trifolii/VP-02/SecA:07 | 2016 | India | *Solanum melongena* |
| KR906721 | *Brinjal little leaf phytoplasma* | Brinjal little leaf/IARI-1/secA:15 | 2015 | India | *Solanum melongena* |
| KX610808 | *Candidatus Phytoplasma trifolii* | Ca.P.trifolii/MKS 01/SecA:16 | 2016 | India | *Solanum melongena* |
| KR906717 | *Brinjal little leaf phytoplasma* | Brinjal little leaf/Odisha-1/secA:15 | 2015 | India | *Solanum melongena* |
| KU297161 | *Brinjal little leaf phytoplasma* | Brinjal little leaf/secA:15 | 2015 | India | *Solanum melongena* |
| EU168742 | *Potato witches'-broom phytoplasma* | Potato witches broom/SecA:07 | 2007 | UK | ------------- |
| MK392370 | *Candidatus Phytoplasma trifolii* | Ca.P.trifolii/TR-Lah19/SecA:19 | 2019 | Turkiye | *Brassica oleracea* |
| EU168744 | *Catharanthus phyllody phytoplasma* | Ca.phyllody phytoplasma/ SecA:07 | 2007 | UK | ------------- |
| KR906722 | *Brinjal little leaf phytoplasma* | Brinjal little leaf/IARI-2/secA:15 | 2015 | India | *Solanum melongena* |
| KR906720 | *Brinjal little leaf phytoplasma* | Brinjal little leaf/Haryana-2/secA:15 | 2015 | India | *Solanum melongena* |
| KR906719 | *Brinjal little leaf phytoplasma* | Brinjal little leaf/Haryana-1/secA:15 | 2015 | India | *Solanum melongena* |
| MW654220 | *Hishimonus phycitis' 16SrVI phytoplasma* | Hishimonus phycitis/HpLH-LKO2/secA:21 | 2021 | India | *Hishimonus phycitis* |
| MW654219 | *Hishimonus phycitis' 16SrVI phytoplasma* | Hishimonus phycitis/HpLH-LKO1/secA:21 | 2021 | India | *Hishimonus phycitis* |
| MW654218 | *Catharanthus roseus' phytoplasma* | Catharanthus roseus/Cr-LKO2/secA:21 | 2021 | India | *Catharanthus roseus* |
| MW654217 | *Catharanthus roseus' phytoplasma* | Catharanthus roseus/Cr-LKO1:21 | 2021 | India | *Catharanthus roseus* |
| MW654216 | *Chrysanthemum coronarium' stunting phytoplasma* | Chrysanthemum coronarium stunting/ChS-LKO2/secA:21 | 2021 | India | *Glebionis coronaria* |
| MW654215 | *Chrysanthemum coronarium' stunting phytoplasma* | Chrysanthemum coronarium stunting/ChS-LKO1/secA:21 | 2021 | India | *Glebionis coronaria* |
| KR906725 | *Brinjal little leaf phytoplasma* | Brinjal little leaf/Assam-1 protein/secA:17 | 2017 | India | *Rauvolfia serpentina* |

| | | | | | |
|---|---|---|---|---|---|
| MG721533 | *Candidatus Phytoplasma trifolii* | Ca.P.trifolii/secA:15 | 2015 | India | *Solanum melongena* |
| KR906726 | *Brinjal little leaf phytoplasma* | Brinjal little leaf/CG-1/secA:15 | 2015 | India | *Solanum melongena* |
| KR906728 | *'Hishimonus phycitis' phytoplasma IARI* | Hishimonus phycitis/IARI/secA:15 | 2015 | India | *Hishimonus phycitis* |
| KR906723 | *Brinjal little leaf phytoplasma* | Brinjal little leaf/IARI-3/secA:15 | 2015 | India | *Solanum melongena* |
| MG566065 | *Candidatus Phytoplasma trifolii* | Ca.P.trifolii/Odisha/secA:17 | 2017 | India | *Solanum tuberosum* |
| KY815101 | *Candidatus Phytoplasma trifolii* | Ca.P.trifolii/Orissa/secA:17 | 2017 | India | *Solanum tuberosum* |
| KJ462044 | *Potato witches'-broom phytoplasma* | Potato witches broom/PWB_ex-TC/SecA:14 | 2014 | USA | *Solanum tuberosum* |
| MG821486 | *Pouzolzia zeylanica' phytoplasma* | Pouzolzia zeylanica /Kamrup/SecA :18 | 2018 | India | *Pouzolzia zeylanica* |
| KC347008 | *Brinjal little leaf phytoplasma* | Brinjal little leaf/Sm-NDL/SecA:12 | 2012 | India | *Solanum melongena* |
| KT335271 | *Brinjal little leaf phytoplasma* | Brinjal little leaf/SecA:15 | 2015 | India | ------------------- |
| MW885174 | *Candidatus Phytoplasma trifolii* | Ca.P.trifolii/SV/SecA:21 | 2021 | India | *Solanum violaceum* |
