## Supplemental Table S8 for "Perilous coexistence: *Chilli Leaf Curl Virus* and *Candidatus Phytoplasma trifolii* infecting *Capsicum annuum*, India"

**Table. S8** Amino acid pairwise percentage identities between identified *Ca. P. trifolii* 16S rRNA (Acc. No. MZ557805) with other similar isolates by use Sequence Demarcation Tool (SDT v1.2) (Muhire et al., 2014).

| Accession number | ACRONYMS | 16S rRNA (percentage amino acid sequence identity) |
|---|---|---|
| MZ557805 | Ca.P.trifolii/16VI-A:21 | 100.00 |
| MN861370 | Ca.P.trifolii/HSD2/16VI-A:19 | 97.401 |
| MN861357 | Ca.P.trifolii/HSD1/16VI-A:19 | 97.401 |
| KX284698 | Brinjal little leaf/LLB_MKS2/16VI-D:14 | 97.401 |
| KX671554 | 'Lectuca sativa' flattened stem/NDL:14 | 97.401 |
| KP027510 | Brinjal little leaf/AP-24/16VI-D:14 | 97.401 |
| KP027517 | Brinjal little leaf/TN-34/16VI-D:14 | 97.401 |
| KP027503 | Brinjal little leaf/AP-17/16VI-D:14 | 97.401 |
| MZ773477 | Ca.P trifolii/THS1/16VI-A:21 | 97.401 |
| JX104336 | Ca.P trifolii/BRGL-2/16VI-A:12 | 97.401 |
| MW648598 | Hishimonus phycitis leafhopper/HpLH-LKO2:21 | 97.401 |
| MW648597 | Hishimonus phycitis leafhopper/HpLH-LKO1:21 | 97.401 |
| MW648596 | Catharanthus roseus phytoplasma/Cr-LKO2:21 | 97.401 |
| MW648595 | Catharanthus roseus /Cr-LK /O1:21 | 97.401 |
| MW648594 | Chrysanthemum coronarium/ChS-LKO2:21 | 97.401 |
| MW648593 | Chrysanthemum coronarium/ChS-LKO1:21 | 97.401 |
| MH547069 | Periwinkle little leaf/7 16S:18 | 97.401 |
| MZ413119 | Acacia arabica/BaSD2:21 | 97.401 |
| MZ413118 | Acacia arabica/BaSD1:21 | 97.401 |
| KP027495 | Brinjal little leaf/KA-04/16VI-A:14 | 97.201 |
| KX284700 | Brinjal little leaf/LLB_MKS4/16VI-A:16 | 95.601 |
| MK660149 | Brassica juncea phyllody /OM2:19 | 95.401 |
| MK660148 | Brassica juncea phyllody/OM1:19 | 95.401 |
| KP119494 | Cucurbita pepo /Neyshabur:16 | 95.401 |
| EF592606 | Iranian cabbage yellows:07 | 95.301 |
| AB279597 | Ca.P.trifolii/16VI-A:06 | 95.101 |
| AF409069 | Ca.P.trifolii/16VI-A:01 | 95.101 |
| AY390261 | Ca.P. trifolii//16VI-A:03 | 95.101 |
| HQ609489 | Potato witches'-broom /CN-99:10 | 94.801 |
| EU346380 | Ca.P. trifolii/AK-1/16VI-A:16 | 94.801 |
| AY500818 | Potato witches'-broom:03 | 94.801 |
| AY500130 | Ca.P. trifolii/16VI-A:03 | 94.801 |
| KY398725 | Ca.P. trifolii/PW32/16VI-A:21 | 94.801 |
| KF178706 | Ca.P.trifolii/16VI-A:13 | 94.801 |
| EU543440 | Ca.P. trifolii/16VI-A:08 | 94.601 |

| | | |
|---|---|---|
| AF409070 | Ca.P.trifolii/16VI-A:01 | 94.601 |
| DQ256089 | Potato witches'-broom:05 | 94.601 |
| HQ589189 | Ca.P.trifolii/CP-1/16VI-A:10 | 94.301 |
| KX284702 | Brinjal little leaf/LLB_MKS6/16VI-A:16 | 82.801 |
| EU346381 | Ca.P.trifolii/AK-2/16VI-A:07 | 82.201 |
| KX284697 | Brinjal little leaf/LLB_MKS1/16VI-A:16 | 80.701 |
| KX284699 | Brinjal little leaf/LLB_MKS3/16VI-D:16 | 78.401 |
| KP027521 | Brinjal little leaf/TN-38//16VI-D:14 | 76.301 |
| X83431 | Mollicutes sp:14 | 74.001 |
| EF534205 | Iranian safflower phyllody:07 | 70.201 |
| FJ525437 | Chile pepper/New Mexico:08 | 62.201 |
| KP027514 | Brinjal little leaf/AP-31/16VI-A:07 | 53.101 |
| EF186821 | Lucerne virescence:06 | 52.201 |
| KP027506 | Brinjal little leaf/AP-20/16VI-A:16 | 49.101 |
| KP027524 | Brinjal little leaf/TN-42/16VI-A:14 | 39.801 |
| HQ436488 | Chile pepper/NM:10 | 25.801 |
| AY496005 | Washington Potato Purple Top/PPT-Wa8:03 | 25.001 |
| KR072666 | Columbia Basin potato purple top:19 | 25.501 |
| OM654559 | 'Sesamum indicum' /Torogh-Mashhad:22 | 24.601 |
| JF508514 | Sesame phyllody/Seph2:11 | 24.801 |
| KX284703 | Brinjal little leaf/LLB_MKS//16VI-D:16 | 24.301 |
| JF508517 | Cucumber phyllody/Cuph2:11 | 23.501 |
| AY270156 | Centaurea solstitialis virescence:03 | 23.901 |
| GU004366 | Potato purple top/PPT-AK1:09 | 23.401 |
| EU543441 | Ca.P. trifolii/16VI-A:08 | 23.301 |
| MF996472 | Ca.P. trifolii/NeWB1/16VI-A:17 | 22.601 |
| AF228053 | Periwinkle little leaf/16S:00 | 22.601 |
| MZ425930 | Brinjal little leaf/T-M2/16VI-D :21 | 22.801 |
| MZ425929 | Brinjal little leaf/T-M1/16VI-D:21 | 22.801 |
| KC178679 | Brinjal little leaf/B-NDL/16VI-D:12 | 22.801 |
| MF996472 | Ca.P. trifolii/NeWB1/16VI-A:17 | 22.601 |
| AF228052 | Brinjal little leaf/16VI-A:00 | 22.701 |
| KC478606 | Brinjal little leaf/BLL-VNS-1/16VI-D:13 | 22.201 |
| EU143330 | 'Vaccinium myrtillus':07 | 22.201 |
| KR072667 | Alaska potato witches'-broom/AK-5:15 | 21.701 |
| GU004367 | Potato purple top/PPT-AK2:09 | 21.201 |
| HQ609491 | Potato witches'-broom/16VI-A/AK-DN1L:10 | 21.401 |
| MF385584 | Ca.P.trifolii/Delaware/16VI-A:17 | 21.301 |
| MK693146 | Ca.P.trifolii/IARI-2/16VI-A:19 | 21.101 |

| | | | |
|---|---|---|---|
| MK369688 | Ca.P. trifolii/IARI-1/16VI-A:19 | | 21.101 |
| KY856746 | 'Hishimonas phycitis'/NDL/16SrVI:21 | | 21.101 |
| KX689245 | Brinjal little leaf/CG-A/16VI-D:17 | | 21.101 |
| KX689234 | Brinjal little leaf/IARI-A/16VI-D:19 | | 21.101 |
| KX641022 | 'Saponaria sp.'stunting and witches' broom/Pune 4:16 | | 20.901 |
| JQ409541 | Brinjal little leaf/16VI-D:12 | | 20.901 |
| MT981209 | Ca.P. trifolii/AP/16VI-A:20 | | 20.201 |
| MF509775 | 'Cannabis sativa'/NDL :17 | | 20.801 |
| AB076404 | Potato witches'-broom:01 | | 20.701 |
| KP027504 | Brinjal little leaf/AP-18/16VI-D:14 | | 20.601 |
| MT739373 | Brinjal little leaf/BLL//16VI-D:20 | | 20.501 |
| MW261864 | Ca.P trifolii/Sonitpur1/16VI-A:14 | | 20.301 |
| MW847614 | Ca.P trifolii/SV1/16VI-A:21 | | 20.201 |
| GU004369 | Potato purple top/PPT-AK5:09 | | 20.101 |
| MW261866 | Ca.P. trifolii/Nagaon2/16VI-A:20 | | 19.801 |
| MW261863 | Ca.P. trifolii/Nagaon1//16VI-A:20 | | 19.801 |
| EF186820 | Brinjal little leaf/BLL//16VI-D:06 | | 19.801 |
| KP027501 | Brinjal little leaf/AP-14/16VI-D:14 | | 19.601 |
| KP027508 | Brinjal little leaf/AP-22/16VI-D:14 | | 19.501 |
| KP027512 | Brinjal little leaf/AP-26/16VI-D:14 | | 19.501 |
| KP027531 | Catharanthus phyllody /KA-02:14 | | 19.301 |
| KP027513 | Brinjal little leaf/AP-28/16VI-D :14 | | 19.301 |
| KP027500 | Brinjal little leaf/AP-13/16VI-D:14 | | 19.301 |
| KP027505 | Brinjal little leaf/AP-19/16VI-D:14 | | 18.801 |
| GU004368 | Potato purple top/PPT-AK4 strain AKpot4:09 | | 17.901 |
| KP027522 | Brinjal little leaf/TN-39/16VI-A:14 | | 17.801 |
