## Supplemental Table S9 for "Perilous coexistence: *Chilli Leaf Curl Virus* and *Candidatus Phytoplasma trifolii* infecting *Capsicum annuum*, India"

**Table. S9** Amino acid pairwise percentage identities between identified with *Ca.P.trifolii* Sec A gene (Acc. No. MZ620707) other similar isolates by use Sequence Demarcation Tool (SDT v1.2) (Muhire et al., 2014).

.

| Accession Number | Acronyms | Sec A (percentage amino acid sequence identity) |
|---|---|---|
| MZ620707 | Ca.P.trifolii/secA:21 | 100.00 |
| KT335271 | Brinjal little leaf/SecA:15 | 96.901 |
| KR906730 | Brinjal little leaf/secA:15 | 96.201 |
| KR906727 | Brinjal little leaf/secA:15 | 96.201 |
| KR906724 | Brinjal little leaf/IARI-4/secA:15 | 96.201 |
| KR906716 | Brinjal little leaf/NOIDA-1/secA:15 | 96.201 |
| KR906729 | Cannabis sativa/NOIDA/secA:15 | 96.201 |
| KR906721 | Brinjal little leaf/IARI-1/secA:15 | 96.101 |
| KR906718 | Brinjal little leaf/Odisha-2/secA:16 | 96.101 |
| KY228385 | Ca.P.trifolii/MKS 06:15 | 96.101 |
| KX610808 | Ca.P.trifolii/MKS 01/SecA:16 | 96.001 |
| KR906717 | Brinjal little leaf/Odisha-1/secA:15 | 96.001 |
| KU297161 | Brinjal little leaf/secA:15 | 96.001 |
| KR906720 | Brinjal little leaf/Haryana-2/secA:15 | 95.701 |
| KR906719 | Brinjal little leaf/Haryana-1/secA:15 | 95.701 |
| MW654220 | Hishimonus phycitis/HpLH-LKO2/secA:21 | 95.701 |
| MW654219 | Hishimonus phycitis/HpLH-LKO1/secA:21 | 95.701 |
| MW654218 | Catharanthus roseus/Cr-LKO2/secA:21 | 95.701 |
| MW654217 | Catharanthus roseus/Cr-LKO1:21 | 95.701 |
| MW654216 | Chrysanthemum coronarium stunting/ChS-LKO2/secA:21 | 95.701 |
| MW654215 | Chrysanthemum coronarium stunting/ChS-LKO1/secA:21 | 95.701 |
| KR906725 | Brinjal little leaf/Assam-1 protein/secA:17 | 95.701 |
| KR906726 | Brinjal little leaf/CG-1/secA:15 | 95.701 |
| KX784498 | Ca.P.trifolii/VP-03/SecA:15 | 95.501 |
| EU168744 | Ca.phyllody phytoplasma/ SecA:07 | 94.901 |
| EU168743 | Brinjal little leaf/secA:16 | 94.901 |
| KJ462044 | Potato witches broom/PWB_ex-TC/SecA:14 | 94.401 |
| EU168742 | Potato witches broom/SecA:07 | 92.901 |
| MG821486 | Pouzolzia zeylanica /Kamrup/SecA :18 | 84.701 |
| KC347008 | Brinjal little leaf/Sm-NDL/SecA:12 | 84.701 |
| MG566065 | Ca.P.trifolii/Odisha/secA:17 | 27.601 |
| KY815101 | Ca.P.trifolii/Orissa/secA:17 | 27.601 |
| MG721533 | Ca.P.trifolii/secA:15 | 23.001 |
| KX622584 | Ca.P.trifolii/VP-02/SecA:07 | 22.101 |

| | | |
|---|---|---|
| KY064175 | Ca.P.trifolii/MKS 04/Sec A:16 | 21.501 |
| KX857660 | Ca.P.trifolii/VP 04/SecA:16 | 21.501 |
| KY073129 | Ca.P.trifolii/MKS 07/SecA:16 | 21.501 |
| KR906728 | Hishimonus phycitis/IARI/secA:15 | 20.001 |
| KR906723 | Brinjal little leaf/IARI-3/secA:15 | 20.001 |
| KR906722 | Brinjal little leaf/IARI-2/secA:15 | 18.501 |
| MK392370 | Ca.P.trifolii/TR-Lah19/SecA:19 | 17.101 |
| KX894792 | Ca.P.trifolii/VP 05/SecA:15 | 17.301 |
