## Supplemental Table S10 for "Perilous coexistence: *Chilli Leaf Curl Virus* and *Candidatus Phytoplasma trifolii* infecting *Capsicum annuum*, India"

**Table. S10** Results of different neutrality tests analysis for ChiLCV-A-GKP (Acc. No. MZ540908), six ORFs, and ChLCuB-GKP (Acc. No. MZ540909) with βC1 gene of identified ChiLCV.

| **Virus component** | **Neutrality tests** | | |
|---|---|---|---|
| | **Tajima's *D*** | **Fu & Li's *D*** | **Fu & Li's *F*** |
| **DNA-A** | -0.35905 | -0.48063 | -0.51826 |
| **CP** | 1.72762 | 0.35934 | 1.08044 |
| **Pre-CP** | -0.46411 | -2.12229 | -1.77351 |
| **Rep** | -0.93360 | 0.28882 | -0.23731 |
| **TrAP** | -1.41601 | -1.54705 | -1.79185 |
| **REn** | -1.42102 | -2.01244 | -2.13846 |
| **C4** | -1.37974 | 0.87840 | 0.02623 |
| **Betasatellite** | -1.27507 | -1.60744 | -1.77381 |
| **βC1** | -0.91271 | -1.16114 | -1.27747 |
