## Supplemental Figure S2 for "Perilous coexistence: *Chilli Leaf Curl Virus* and *Candidatus Phytoplasma trifolii* infecting *Capsicum annuum*, India"

a)

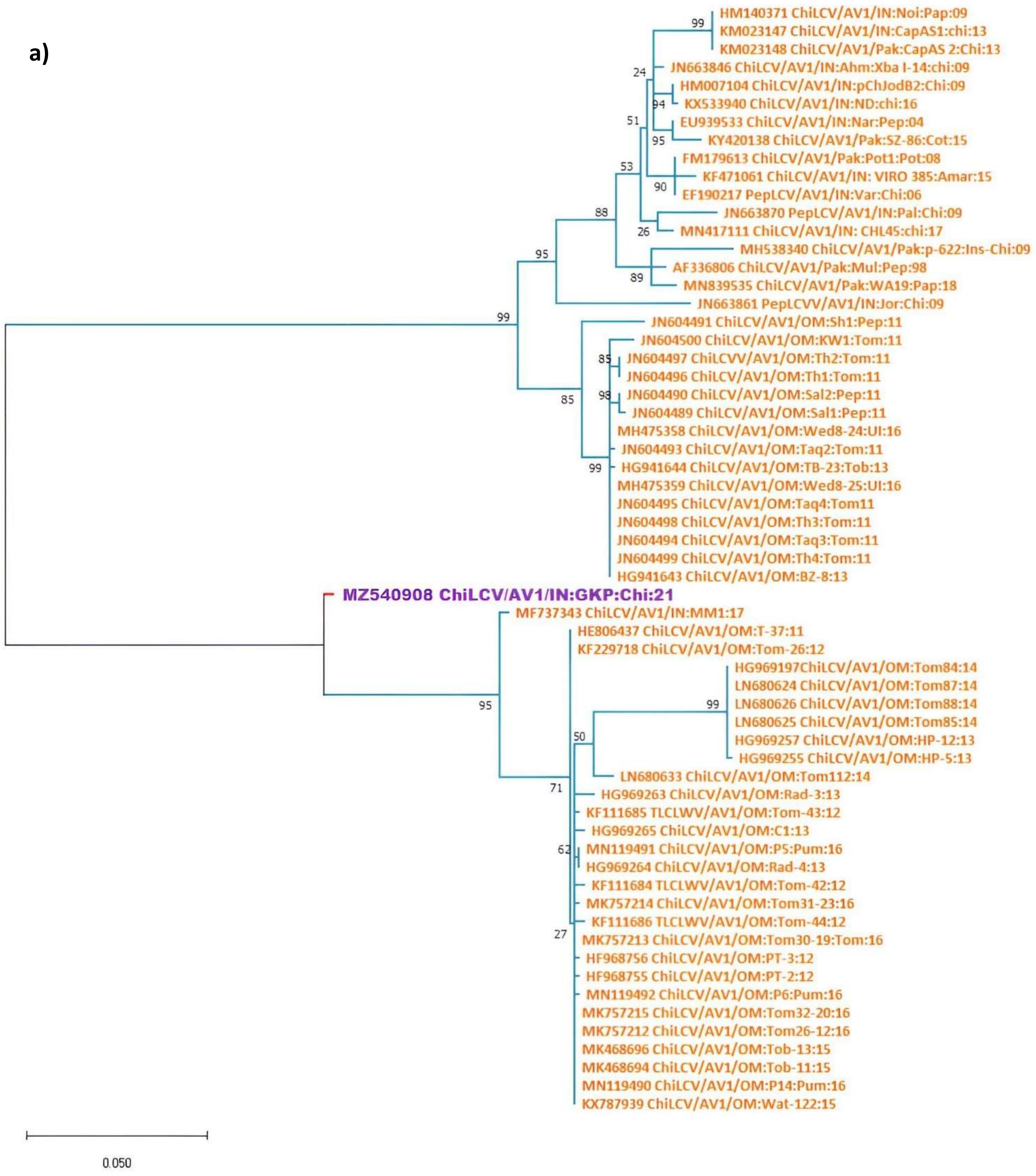
HM140371 ChiLCV/AV1/IN:Noi:Pap:09
KM023147 ChiLCV/AV1/IN:CapAS1:chi:13
KM023148 ChiLCV/AV1/Pak:CapAS 2:Chi:13
JN663846 ChiLCV/AV1/IN:Ahm:Xba I-14:chi:09
HM007104 ChiLCV/AV1/IN:pChJodB2:Chi:09
KX533940 ChiLCV/AV1/IN:ND:chi:16
EU939533 ChiLCV/AV1/IN:Nar:Pep:04
KY420138 ChiLCV/AV1/Pak:SZ-86:Cot:15
FM179613 ChiLCV/AV1/Pak:Pot1:Pot:08
KF471061 ChiLCV/AV1/IN: VIRO 385:Amar:15
EF190217 PepLCV/AV1/IN:Var:Chi:06
JN663870 PepLCV/AV1/IN:Pal:Chi:09
MN417111 ChiLCV/AV1/IN: CHL45:chi:17
MH538340 ChiLCV/AV1/Pak:p-622:Ins-Chi:09
AF336806 ChiLCV/AV1/Pak:Mul:Pep:98
MN839535 ChiLCV/AV1/Pak:WA19:Pap:18
JN663861 PepLCVV/AV1/IN:Jor:Chi:09
JN604491 ChiLCV/AV1/OM:Sh1:Pep:11
JN604500 ChiLCV/AV1/OM:KW1:Tom:11
JN604497 ChiLCVV/AV1/OM:Th2:Tom:11
JN604496 ChiLCV/AV1/OM:Th1:Tom:11
JN604490 ChiLCV/AV1/OM:Sal2:Pep:11
JN604489 ChiLCV/AV1/OM:Sal1:Pep:11
MH475358 ChiLCV/AV1/OM:Wed8-24:Ul:16
JN604493 ChiLCV/AV1/OM:Taq2:Tom:11
HG941644 ChiLCV/AV1/OM:TB-23:Tob:13
MH475359 ChiLCV/AV1/OM:Wed8-25:Ul:16
JN604495 ChiLCV/AV1/OM:Taq4:Tom11
JN604498 ChiLCV/AV1/OM:Th3:Tom:11
JN604494 ChiLCV/AV1/OM:Taq3:Tom:11
JN604499 ChiLCV/AV1/OM:Th4:Tom:11
HG941643 ChiLCV/AV1/OM:BZ-8:13
MZ540908 ChiLCV/AV1/IN:GKP:Chi:21
MF737343 ChiLCV/AV1/IN:MM1:17
HE806437 ChiLCV/AV1/OM:T-37:11
KF229718 ChiLCV/AV1/OM:Tom-26:12
HG969197ChiLCV/AV1/OM:Tom84:14
LN680624 ChiLCV/AV1/OM:Tom87:14
LN680626 ChiLCV/AV1/OM:Tom88:14
LN680625 ChiLCV/AV1/OM:Tom85:14
HG969257 ChiLCV/AV1/OM:HP-12:13
HG969255 ChiLCV/AV1/OM:HP-5:13
LN680633 ChiLCV/AV1/OM:Tom112:14
HG969263 ChiLCV/AV1/OM:Rad-3:13
KF111685 TLCLWV/AV1/OM:Tom-43:12
HG969265 ChiLCV/AV1/OM:C1:13
MN119491 ChiLCV/AV1/OM:P5:Pum:16
HG969264 ChiLCV/AV1/OM:Rad-4:13
KF111684 TLCLWV/AV1/OM:Tom-42:12
MK757214 ChiLCV/AV1/OM:Tom31-23:16
KF111686 TLCLWV/AV1/OM:Tom-44:12
MK757213 ChiLCV/AV1/OM:Tom30-19:Tom:16
HF968756 ChiLCV/AV1/OM:PT-3:12
HF968755 ChiLCV/AV1/OM:PT-2:12
MN119492 ChiLCV/AV1/OM:P6:Pum:16
MK757215 ChiLCV/AV1/OM:Tom32-20:16
MK757212 ChiLCV/AV1/OM:Tom26-12:16
MK468696 ChiLCV/AV1/OM:Tob-13:15
MK468694 ChiLCV/AV1/OM:Tob-11:15
MN119490 ChiLCV/AV1/OM:P14:Pum:16
KX787939 ChiLCV/AV1/OM:Wat-122:15
0.050

b)

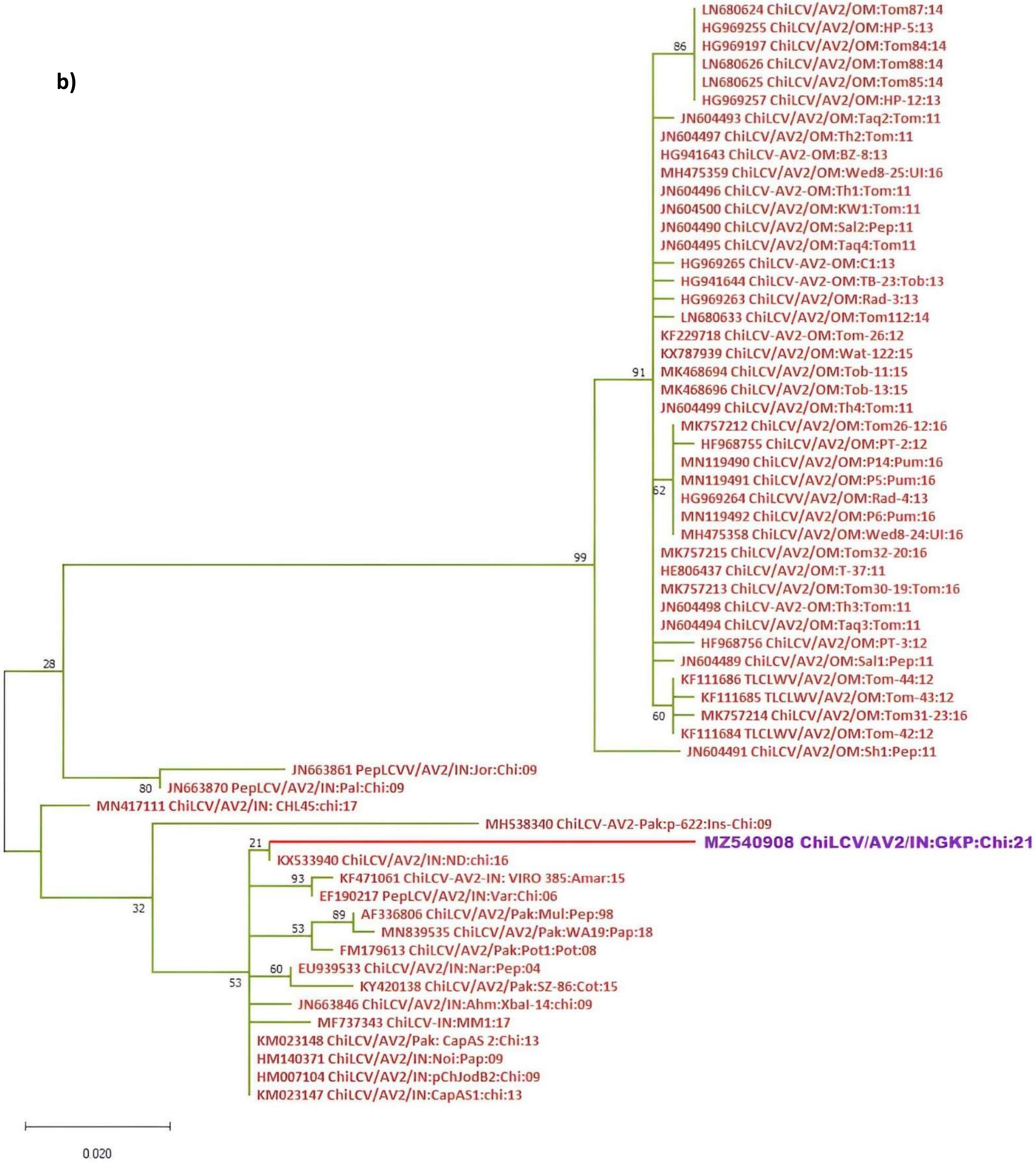
LN680624 ChiLCV/AV2/OM:Tom87:14
HG969255 ChiLCV/AV2/OM:HP-5:13
HG969197 ChiLCV/AV2/OM:Tom84:14
LN680626 ChiLCV/AV2/OM:Tom88:14
LN680625 ChiLCV/AV2/OM:Tom85:14
HG969257 ChiLCV/AV2/OM:HP-12:13
JN604493 ChiLCV/AV2/OM:Taq2:Tom:11
JN604497 ChiLCV/AV2/OM:Th2:Tom:11
HG941643 ChiLCV-AV2-OM:BZ-8:13
MH475359 ChiLCV/AV2/OM:Wed8-25:UI:16
JN604496 ChiLCV-AV2-OM:Th1:Tom:11
JN604500 ChiLCV/AV2/OM:KW1:Tom:11
JN604490 ChiLCV/AV2/OM:Sal2:Pep:11
JN604495 ChiLCV/AV2/OM:Taq4:Tom11
HG969265 ChiLCV-AV2-OM:C1:13
HG941644 ChiLCV-AV2-OM:TB-23:Tob:13
HG969263 ChiLCV/AV2/OM:Rad-3:13
LN680633 ChiLCV/AV2/OM:Tom112:14
KF229718 ChiLCV-AV2-OM:Tom-26:12
KX787939 ChiLCV/AV2/OM:Wat-122:15
MK468694 ChiLCV/AV2/OM:Tob-11:15
MK468696 ChiLCV/AV2/OM:Tob-13:15
JN604499 ChiLCV/AV2/OM:Th4:Tom:11
MK757212 ChiLCV/AV2/OM:Tom26-12:16
HF968755 ChiLCV/AV2/OM:PT-2:12
MN119490 ChiLCV/AV2/OM:P14:Pum:16
MN119491 ChiLCV/AV2/OM:P5:Pum:16
HG969264 ChiLCVV/AV2/OM:Rad-4:13
MN119492 ChiLCV/AV2/OM:P6:Pum:16
MH475358 ChiLCV/AV2/OM:Wed8-24:UI:16
MK757215 ChiLCV/AV2/OM:Tom32-20:16
HE806437 ChiLCV/AV2/OM:T-37:11
MK757213 ChiLCV/AV2/OM:Tom30-19:Tom:16
JN604498 ChiLCV-AV2-OM:Th3:Tom:11
JN604494 ChiLCV/AV2/OM:Taq3:Tom:11
HF968756 ChiLCV/AV2/OM:PT-3:12
JN604489 ChiLCV/AV2/OM:Sal1:Pep:11
KF111686 TLCLWV/AV2/OM:Tom-44:12
KF111685 TLCLWV/AV2/OM:Tom-43:12
MK757214 ChiLCV/AV2/OM:Tom31-23:16
KF111684 TLCLWV/AV2/OM:Tom-42:12
JN604491 ChiLCV/AV2/OM:Sh1:Pep:11
JN663861 PepLCVV/AV2/IN:Jor:Chi:09
JN663870 PepLCV/AV2/IN:Pal:Chi:09
MN417111 ChiLCV/AV2/IN: CHL45:chi:17
MH538340 ChiLCV-AV2-Pak:p-622:Ins-Chi:09
MZ540908 ChiLCV/AV2/IN:GKP:Chi:21
KX533940 ChiLCV/AV2/IN:ND:chi:16
KF471061 ChiLCV-AV2-IN: VIRO 385:Amar:15
EF190217 PepLCV/AV2/IN:Var:Chi:06
AF336806 ChiLCV/AV2/Pak:Mul:Pep:98
MN839535 ChiLCV/AV2/Pak:WA19:Pap:18
FM179613 ChiLCV/AV2/Pak:Pot1:Pot:08
EU939533 ChiLCV/AV2/IN:Nar:Pep:04
KY420138 ChiLCV/AV2/Pak:SZ-86:Cot:15
JN663846 ChiLCV/AV2/IN:Ahm:XbaI-14:chi:09
MF737343 ChiLCV-IN:MM1:17
KM023148 ChiLCV/AV2/Pak: CapAS 2:Chi:13
HM140371 ChiLCV/AV2/IN:Noi:Pap:09
HM007104 ChiLCV/AV2/IN:pChJodB2:Chi:09
KM023147 ChiLCV/AV2/IN:CapAS1:chi:13
86
91
62
60
99
28
80
21
93
89
53
32
60
53
0.020

c)

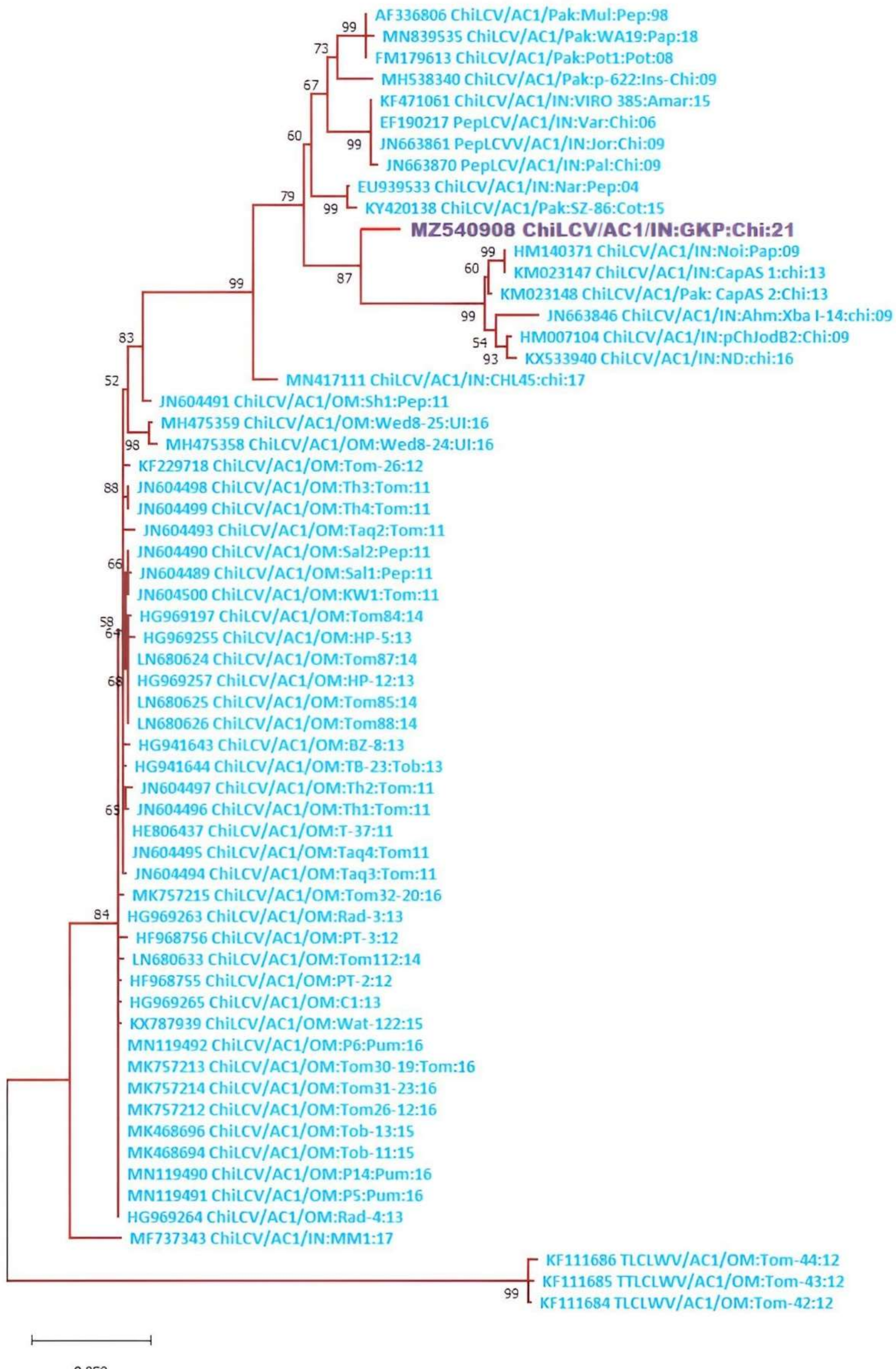
AF336806 ChiLCV/AC1/Pak:Mul:Pep:98
MN839535 ChiLCV/AC1/Pak:WA19:Pap:18
FM179613 ChiLCV/AC1/Pak:Pot1:Pot:08
MH538340 ChiLCV/AC1/Pak:p-622:Ins-Chi:09
KF471061 ChiLCV/AC1/IN:VIRO 385:Amar:15
EF190217 PepLCV/AC1/IN:Var:Chi:06
JN663861 PepLCVV/AC1/IN:Jor:Chi:09
JN663870 PepLCV/AC1/IN:Pal:Chi:09
EU939533 ChiLCV/AC1/IN:Nar:Pep:04
KY420138 ChiLCV/AC1/Pak:SZ-86:Cot:15
MZ540908 ChiLCV/AC1/IN:GKP:Chi:21
HM140371 ChiLCV/AC1/IN:Noi:Pap:09
KM023147 ChiLCV/AC1/IN:CapAS 1:chi:13
KM023148 ChiLCV/AC1/Pak: CapAS 2:Chi:13
JN663846 ChiLCV/AC1/IN:Ahm:Xba I-14:chi:09
HM007104 ChiLCV/AC1/IN:pChJodB2:Chi:09
KX533940 ChiLCV/AC1/IN:ND:chi:16
MN417111 ChiLCV/AC1/IN:CHL45:chi:17
JN604491 ChiLCV/AC1/OM:Sh1:Pep:11
MH475359 ChiLCV/AC1/OM:Wed8-25:Ul:16
MH475358 ChiLCV/AC1/OM:Wed8-24:Ul:16
KF229718 ChiLCV/AC1/OM:Tom-26:12
JN604498 ChiLCV/AC1/OM:Th3:Tom:11
JN604499 ChiLCV/AC1/OM:Th4:Tom:11
JN604493 ChiLCV/AC1/OM:Taq2:Tom:11
JN604490 ChiLCV/AC1/OM:Sal2:Pep:11
JN604489 ChiLCV/AC1/OM:Sal1:Pep:11
JN604500 ChiLCV/AC1/OM:KW1:Tom:11
HG969197 ChiLCV/AC1/OM:Tom84:14
HG969255 ChiLCV/AC1/OM:HP-5:13
LN680624 ChiLCV/AC1/OM:Tom87:14
HG969257 ChiLCV/AC1/OM:HP-12:13
LN680625 ChiLCV/AC1/OM:Tom85:14
LN680626 ChiLCV/AC1/OM:Tom88:14
HG941643 ChiLCV/AC1/OM:BZ-8:13
HG941644 ChiLCV/AC1/OM:TB-23:Tob:13
JN604497 ChiLCV/AC1/OM:Th2:Tom:11
JN604496 ChiLCV/AC1/OM:Th1:Tom:11
HE806437 ChiLCV/AC1/OM:T-37:11
JN604495 ChiLCV/AC1/OM:Taq4:Tom11
JN604494 ChiLCV/AC1/OM:Taq3:Tom:11
MK757215 ChiLCV/AC1/OM:Tom32-20:16
HG969263 ChiLCV/AC1/OM:Rad-3:13
HF968756 ChiLCV/AC1/OM:PT-3:12
LN680633 ChiLCV/AC1/OM:Tom112:14
HF968755 ChiLCV/AC1/OM:PT-2:12
HG969265 ChiLCV/AC1/OM:C1:13
KX787939 ChiLCV/AC1/OM:Wat-122:15
MN119492 ChiLCV/AC1/OM:P6:Pum:16
MK757213 ChiLCV/AC1/OM:Tom30-19:Tom:16
MK757214 ChiLCV/AC1/OM:Tom31-23:16
MK757212 ChiLCV/AC1/OM:Tom26-12:16
MK468696 ChiLCV/AC1/OM:Tob-13:15
MK468694 ChiLCV/AC1/OM:Tob-11:15
MN119490 ChiLCV/AC1/OM:P14:Pum:16
MN119491 ChiLCV/AC1/OM:P5:Pum:16
HG969264 ChiLCV/AC1/OM:Rad-4:13
MF737343 ChiLCV/AC1/IN:MM1:17
KF111686 TLCLWV/AC1/OM:Tom-44:12
KF111685 TTLCLWV/AC1/OM:Tom-43:12
KF111684 TLCLWV/AC1/OM:Tom-42:12

d)

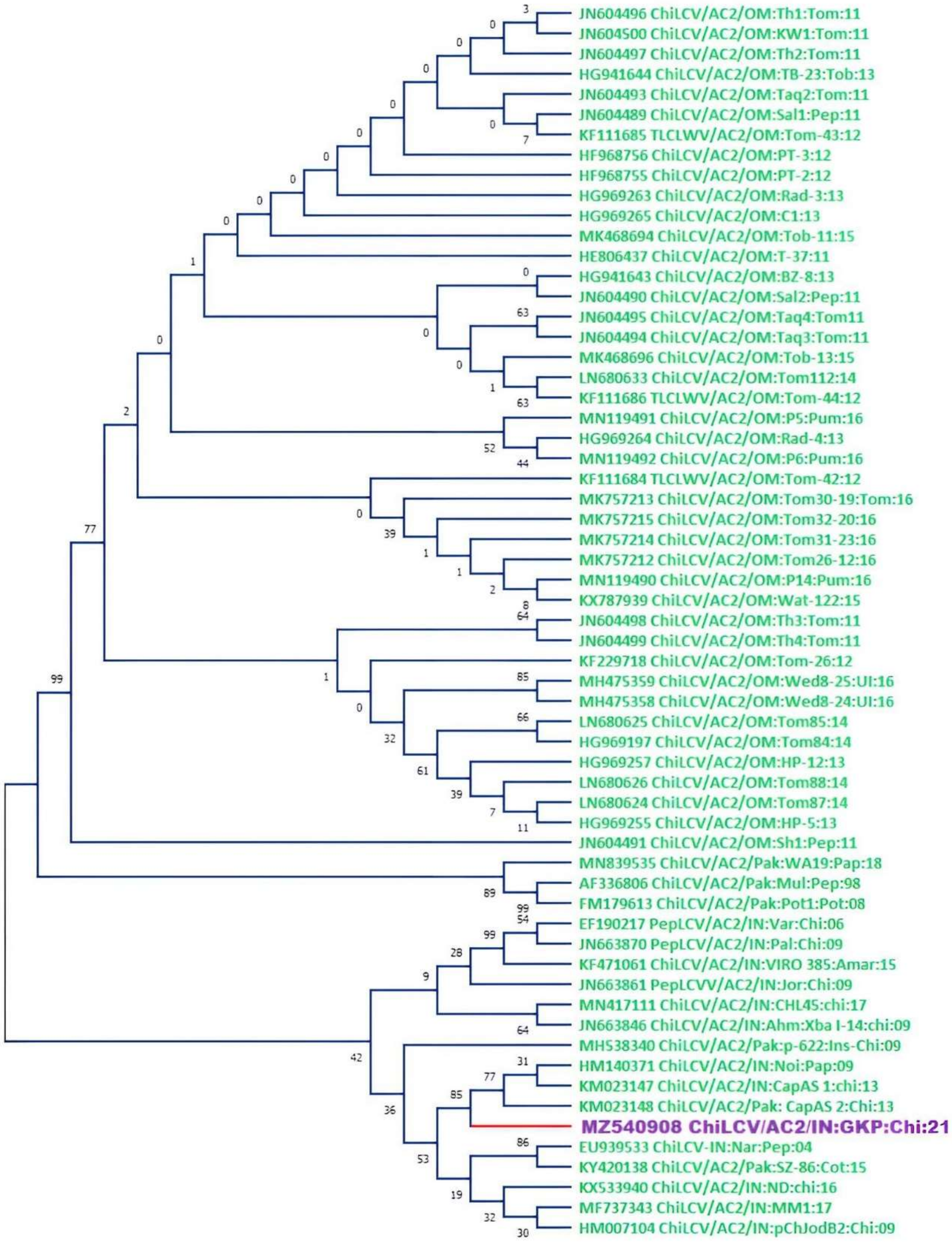
JN604496 ChiLCV/AC2/OM:Th1:Tom:11
JN604500 ChiLCV/AC2/OM:KW1:Tom:11
JN604497 ChiLCV/AC2/OM:Th2:Tom:11
HG941644 ChiLCV/AC2/OM:TB-23:Tob:13
JN604493 ChiLCV/AC2/OM:Taq2:Tom:11
JN604489 ChiLCV/AC2/OM:Sal1:Pep:11
KF111685 TLCLWV/AC2/OM:Tom-43:12
HF968756 ChiLCV/AC2/OM:PT-3:12
HF968755 ChiLCV/AC2/OM:PT-2:12
HG969263 ChiLCV/AC2/OM:Rad-3:13
HG969265 ChiLCV/AC2/OM:C1:13
MK468694 ChiLCV/AC2/OM:Tob-11:15
HE806437 ChiLCV/AC2/OM:T-37:11
HG941643 ChiLCV/AC2/OM:BZ-8:13
JN604490 ChiLCV/AC2/OM:Sal2:Pep:11
JN604495 ChiLCV/AC2/OM:Taq4:Tom11
JN604494 ChiLCV/AC2/OM:Taq3:Tom:11
MK468696 ChiLCV/AC2/OM:Tob-13:15
LN680633 ChiLCV/AC2/OM:Tom112:14
KF111686 TLCLWV/AC2/OM:Tom-44:12
MN119491 ChiLCV/AC2/OM:P5:Pum:16
HG969264 ChiLCV/AC2/OM:Rad-4:13
MN119492 ChiLCV/AC2/OM:P6:Pum:16
KF111684 TLCLWV/AC2/OM:Tom-42:12
MK757213 ChiLCV/AC2/OM:Tom30-19:Tom:16
MK757215 ChiLCV/AC2/OM:Tom32-20:16
MK757214 ChiLCV/AC2/OM:Tom31-23:16
MK757212 ChiLCV/AC2/OM:Tom26-12:16
MN119490 ChiLCV/AC2/OM:P14:Pum:16
KX787939 ChiLCV/AC2/OM:Wat-122:15
JN604498 ChiLCV/AC2/OM:Th3:Tom:11
JN604499 ChiLCV/AC2/OM:Th4:Tom:11
KF229718 ChiLCV/AC2/OM:Tom-26:12
MH475359 ChiLCV/AC2/OM:Wed8-25:UI:16
MH475358 ChiLCV/AC2/OM:Wed8-24:UI:16
LN680625 ChiLCV/AC2/OM:Tom85:14
HG969197 ChiLCV/AC2/OM:Tom84:14
HG969257 ChiLCV/AC2/OM:HP-12:13
LN680626 ChiLCV/AC2/OM:Tom88:14
LN680624 ChiLCV/AC2/OM:Tom87:14
HG969255 ChiLCV/AC2/OM:HP-5:13
JN604491 ChiLCV/AC2/OM:Sh1:Pep:11
MN839535 ChiLCV/AC2/Pak:WA19:Pap:18
AF336806 ChiLCV/AC2/Pak:Mul:Pep:98
FM179613 ChiLCV/AC2/Pak:Pot1:Pot:08
EF190217 PepLCV/AC2/IN:Var:Chi:06
JN663870 PepLCV/AC2/IN:Pal:Chi:09
KF471061 ChiLCV/AC2/IN:VIRO 385:Amar:15
JN663861 PepLCVV/AC2/IN:Jor:Chi:09
MN417111 ChiLCV/AC2/IN:CHL45:chi:17
JN663846 ChiLCV/AC2/IN:Ahm:Xba I-14:chi:09
MH538340 ChiLCV/AC2/Pak:p-622:Ins-Chi:09
HM140371 ChiLCV/AC2/IN:Noi:Pap:09
KM023147 ChiLCV/AC2/IN:CapAS 1:chi:13
KM023148 ChiLCV/AC2/Pak: CapAS 2:Chi:13
MZ540908 ChiLCV/AC2/IN:GKP:Chi:21
EU939533 ChiLCV-IN:Nar:Pep:04
KY420138 ChiLCV/AC2/Pak:SZ-86:Cot:15
KX533940 ChiLCV/AC2/IN:ND:chi:16
MF737343 ChiLCV/AC2/IN:MM1:17
HM007104 ChiLCV/AC2/IN:pChJodB2:Chi:09

e)

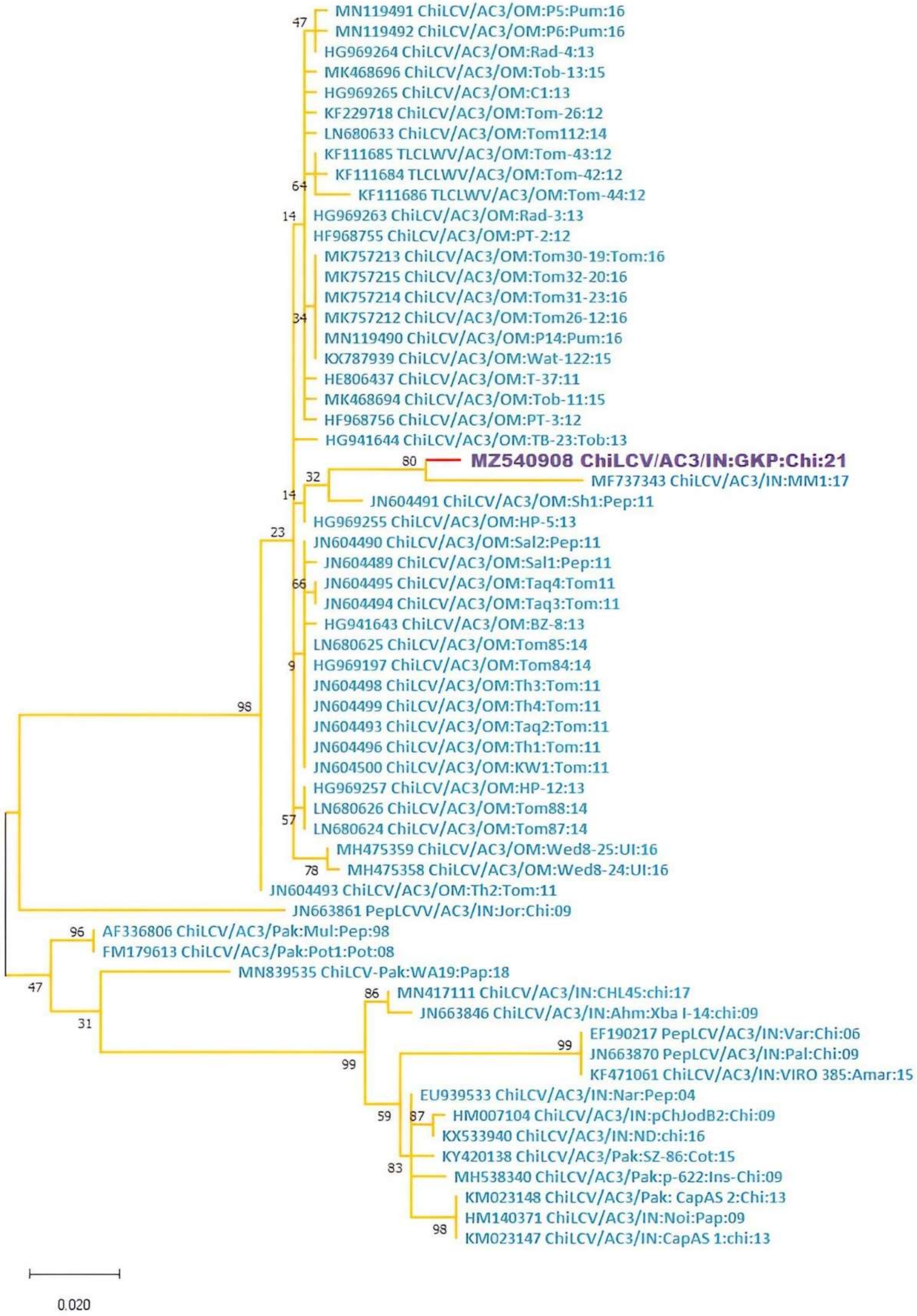
MN119491 ChiLCV/AC3/OM:P5:Pum:16
MN119492 ChiLCV/AC3/OM:P6:Pum:16
HG969264 ChiLCV/AC3/OM:Rad-4:13
MK468696 ChiLCV/AC3/OM:Tob-13:15
HG969265 ChiLCV/AC3/OM:C1:13
KF229718 ChiLCV/AC3/OM:Tom-26:12
LN680633 ChiLCV/AC3/OM:Tom112:14
KF111685 TLCLWV/AC3/OM:Tom-43:12
KF111684 TLCLWV/AC3/OM:Tom-42:12
KF111686 TLCLWV/AC3/OM:Tom-44:12
HG969263 ChiLCV/AC3/OM:Rad-3:13
HF968755 ChiLCV/AC3/OM:PT-2:12
MK757213 ChiLCV/AC3/OM:Tom30-19:Tom:16
MK757215 ChiLCV/AC3/OM:Tom32-20:16
MK757214 ChiLCV/AC3/OM:Tom31-23:16
MK757212 ChiLCV/AC3/OM:Tom26-12:16
MN119490 ChiLCV/AC3/OM:P14:Pum:16
KX787939 ChiLCV/AC3/OM:Wat-122:15
HE806437 ChiLCV/AC3/OM:T-37:11
MK468694 ChiLCV/AC3/OM:Tob-11:15
HF968756 ChiLCV/AC3/OM:PT-3:12
HG941644 ChiLCV/AC3/OM:TB-23:Tob:13
MZ540908 ChiLCV/AC3/IN:GKP:Chi:21
MF737343 ChiLCV/AC3/IN:MM1:17
JN604491 ChiLCV/AC3/OM:Sh1:Pep:11
HG969255 ChiLCV/AC3/OM:HP-5:13
JN604490 ChiLCV/AC3/OM:Sal2:Pep:11
JN604489 ChiLCV/AC3/OM:Sal1:Pep:11
JN604495 ChiLCV/AC3/OM:Taq4:Tom11
JN604494 ChiLCV/AC3/OM:Taq3:Tom:11
HG941643 ChiLCV/AC3/OM:BZ-8:13
LN680625 ChiLCV/AC3/OM:Tom85:14
HG969197 ChiLCV/AC3/OM:Tom84:14
JN604498 ChiLCV/AC3/OM:Th3:Tom:11
JN604499 ChiLCV/AC3/OM:Th4:Tom:11
JN604493 ChiLCV/AC3/OM:Taq2:Tom:11
JN604496 ChiLCV/AC3/OM:Th1:Tom:11
JN604500 ChiLCV/AC3/OM:KW1:Tom:11
HG969257 ChiLCV/AC3/OM:HP-12:13
LN680626 ChiLCV/AC3/OM:Tom88:14
LN680624 ChiLCV/AC3/OM:Tom87:14
MH475359 ChiLCV/AC3/OM:Wed8-25:UI:16
MH475358 ChiLCV/AC3/OM:Wed8-24:UI:16
JN604493 ChiLCV/AC3/OM:Th2:Tom:11
JN663861 PepLCVV/AC3/IN:Jor:Chi:09
AF336806 ChiLCV/AC3/Pak:Mul:Pep:98
FM179613 ChiLCV/AC3/Pak:Pot1:Pot:08
MN839535 ChiLCV-Pak:WA19:Pap:18
MN417111 ChiLCV/AC3/IN:CHL45:chi:17
JN663846 ChiLCV/AC3/IN:Ahm:Xba I-14:chi:09
EF190217 PepLCV/AC3/IN:Var:Chi:06
JN663870 PepLCV/AC3/IN:Pal:Chi:09
KF471061 ChiLCV/AC3/IN:VIRO 385:Amar:15
EU939533 ChiLCV/AC3/IN:Nar:Pep:04
HM007104 ChiLCV/AC3/IN:pChJodB2:Chi:09
KX533940 ChiLCV/AC3/IN:ND:chi:16
KY420138 ChiLCV/AC3/Pak:SZ-86:Cot:15
MH538340 ChiLCV/AC3/Pak:p-622:Ins-Chi:09
KM023148 ChiLCV/AC3/Pak: CapAS 2:Chi:13
HM140371 ChiLCV/AC3/IN:Noi:Pap:09
KM023147 ChiLCV/AC3/IN:CapAS 1:chi:13
47
64
14
34
80
32
14
23
66
9
98
57
78
96
47
31
86
99
99
59
87
83
98
0.020

f)

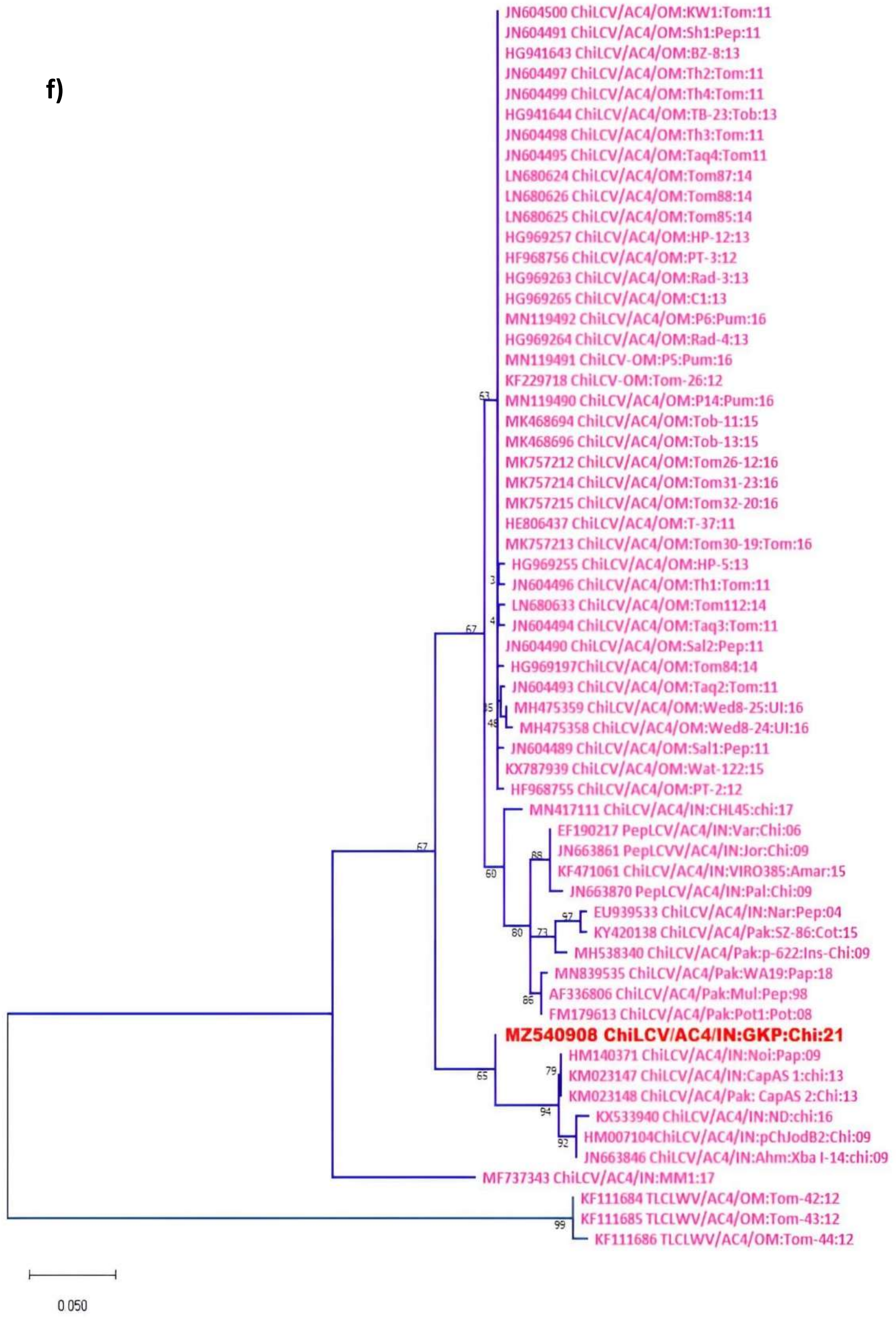


**Figure S2** Maximum-likelihood tree technique was used to create a phylogenetic dendrogram with the best nucleotide model for all 6 ORFs of ChiLCV-A-GKP (Acc. No. MZ540908); a) AV1 (Coat protein) model T92+G+I, b) AV2 (Pre-Coat protein) model JC+G, c) AC1 (Replication initiator protein) model T92+G, d) AC2 (Transcription activator protein) model K2+G, e) AC3 (Replication enhancer protein) model JC+G and f) AC4 K2+G; In MEGA v.10, each sequence was aligned by applying CLUSTAL W with bootstrap values of 1000 replicates (at nodes) (Kumar et al., 2018).

**Figure S2** Maximum-likelihood tree technique was used to create a phylogenetic dendrogram with the best nucleotide model for all 6 ORFs of ChiLCV-A-GKP (Acc. No. MZ540908); a) AV1 (Coat protein) model T92+G+I, b) AV2 (Pre-Coat protein) model JC+G, c) AC1 (Replication initiator protein) model T92+G, d) AC2 (Transcription activator protein) model K2+G, e) AC3 (Replication enhancer protein) model JC+G and f) AC4 K2+G; In MEGA v.10, each sequence was aligned by applying CLUSTAL W with bootstrap values of 1000 replicates (at nodes) (Kumar et al., 2018).
