## Supplemental Figure S1 for "Perilous coexistence: *Chilli Leaf Curl Virus* and *Candidatus Phytoplasma trifolii* infecting *Capsicum annuum*, India"

a)

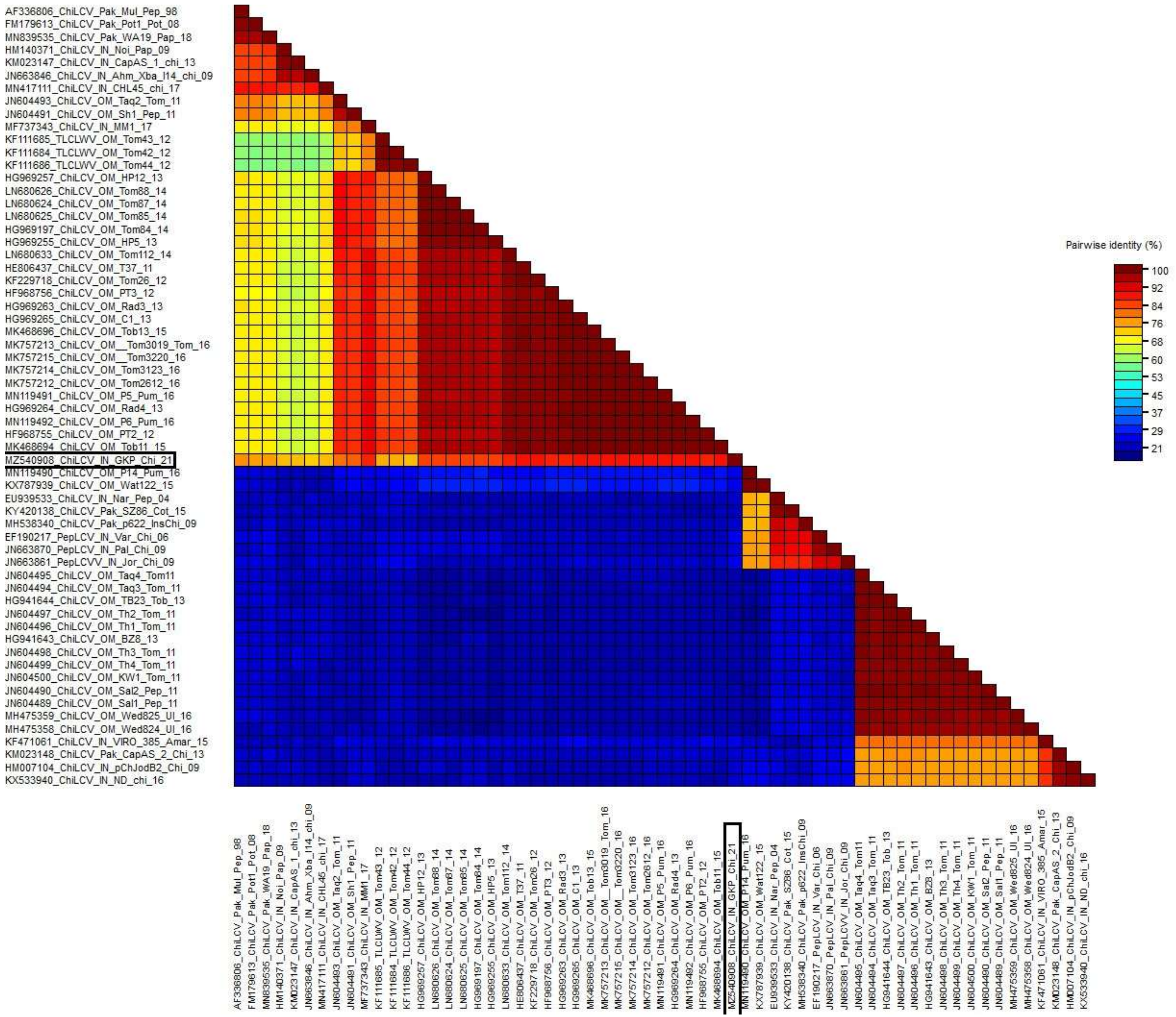
AF336806_ChiLCV_Pak_Mul_Pep_98
FM179613_ChiLCV_Pak_Pot1_Pot_08
MN839535_ChiLCV_Pak_WA19_Pap_18
HM140371_ChiLCV_IN_Noi_Pap_09
KM023147_ChiLCV_IN_CapAS_1_chi_13
JN663846_ChiLCV_IN_Ahm_Xba_I14_chi_09
MN417111_ChiLCV_IN_CHL45_chi_17
JN604493_ChiLCV_OM_Taq2_Tom_11
JN604491_ChiLCV_OM_Sh1_Pep_11
MF737343_ChiLCV_IN_MM1_17
KF111685_TLCLWV_OM_Tom43_12
KF111684_TLCLWV_OM_Tom42_12
KF111686_TLCLWV_OM_Tom44_12
HG969257_ChiLCV_OM_HP12_13
LN680626_ChiLCV_OM_Tom88_14
LN680624_ChiLCV_OM_Tom87_14
LN680625_ChiLCV_OM_Tom85_14
HG969197_ChiLCV_OM_Tom84_14
HG969255_ChiLCV_OM_HP5_13
LN680633_ChiLCV_OM_Tom112_14
HE806437_ChiLCV_OM_T37_11
KF229718_ChiLCV_OM_Tom26_12
HF968756_ChiLCV_OM_PT3_12
HG969263_ChiLCV_OM_Rad3_13
HG969265_ChiLCV_OM_C1_13
MK468696_ChiLCV_OM_Tob13_15
MK757213_ChiLCV_OM__Tom3019_Tom_16
MK757215_ChiLCV_OM__Tom3220_16
MK757214_ChiLCV_OM_Tom3123_16
MK757212_ChiLCV_OM_Tom2612_16
MN119491_ChiLCV_OM_P5_Pum_16
HG969264_ChiLCV_OM_Rad4_13
MN119492_ChiLCV_OM_P6_Pum_16
HF968755_ChiLCV_OM_PT2_12
MK468694_ChiLCV_OM_Tob11_15
MZ540908_ChiLCV_IN_GKP_Chi_21
MN119490_ChiLCV_OM_P14_Pum_16
KX787939_ChiLCV_OM_Wat122_15
EU939533_ChiLCV_IN_Nar_Pep_04
KY420138_ChiLCV_Pak_SZ86_Cot_15
MH538340_ChiLCV_Pak_p622_InsChi_09
EF190217_PepLCV_IN_Var_Chi_06
JN663870_PepLCV_IN_Pal_Chi_09
JN663861_PepLCVV_IN_Jor_Chi_09
JN604495_ChiLCV_OM_Taq4_Tom11
JN604494_ChiLCV_OM_Taq3_Tom_11
HG941644_ChiLCV_OM_TB23_Tob_13
JN604497_ChiLCV_OM_Th2_Tom_11
JN604496_ChiLCV_OM_Th1_Tom_11
HG941643_ChiLCV_OM_BZ8_13
JN604498_ChiLCV_OM_Th3_Tom_11
JN604499_ChiLCV_OM_Th4_Tom_11
JN604500_ChiLCV_OM_KW1_Tom_11
JN604490_ChiLCV_OM_Sal2_Pep_11
JN604489_ChiLCV_OM_Sal1_Pep_11
MH475359_ChiLCV_OM_Wed825_Ul_16
MH475358_ChiLCV_OM_Wed824_Ul_16
KF471061_ChiLCV_IN_VIRO_385_Amar_15
KM023148_ChiLCV_Pak_CapAS_2_Chi_13
HM007104_ChiLCV_IN_pChJodB2_Chi_09
KX533940_ChiLCV_IN_ND_chi_16
Pairwise identity (%)
100
92
84
76
68
60
53
45
37
29
21

b)

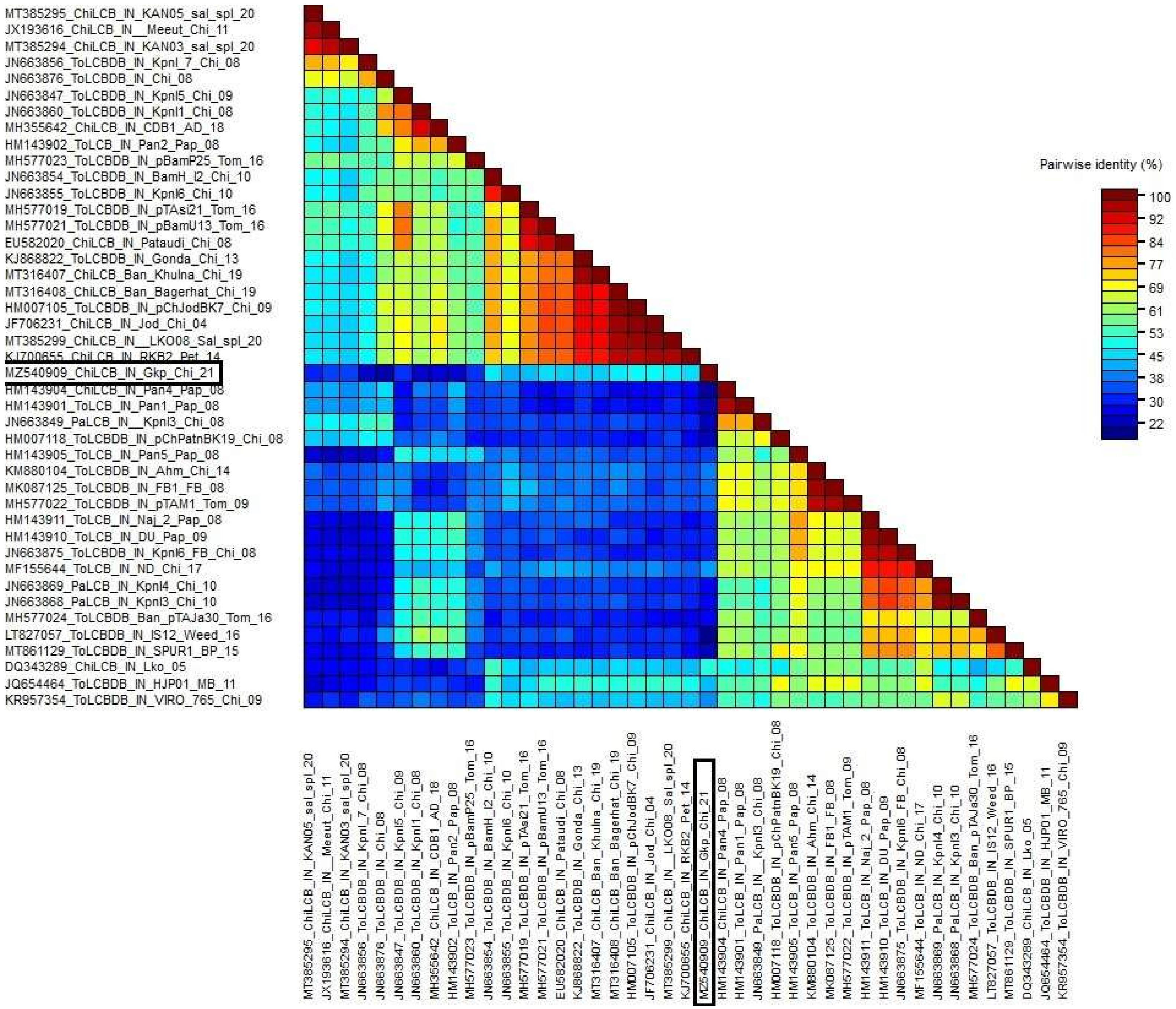
MT385295_ChiLCB_IN_KAN05_sal_spl_20
JX193616_ChiLCB_IN__Meeut_Chi_11
MT385294_ChiLCB_IN_KAN03_sal_spl_20
JN663856_ToLCBDB_IN_Kpnl_7_Chi_08
JN663876_ToLCBDB_IN_Chi_08
JN663847_ToLCBDB_IN_Kpnl5_Chi_09
JN663860_ToLCBDB_IN_Kpnl1_Chi_08
MH355642_ChiLCB_IN_CDB1_AD_18
HM143902_ToLCB_IN_Pan2_Pap_08
MH577023_ToLCBDB_IN_pBamP25_Tom_16
JN663854_ToLCBDB_IN_BamH_I2_Chi_10
JN663855_ToLCBDB_IN_Kpnl6_Chi_10
MH577019_ToLCBDB_IN_pTAsi21_Tom_16
MH577021_ToLCBDB_IN_pBamU13_Tom_16
EU582020_ChiLCB_IN_Pataudi_Chi_08
KJ868822_ToLCBDB_IN_Gonda_Chi_13
MT316407_ChiLCB_Ban_Khulna_Chi_19
MT316408_ChiLCB_Ban_Bagerhat_Chi_19
HM007105_ToLCBDB_IN_pChJodBK7_Chi_09
JF706231_ChiLCB_IN_Jod_Chi_04
MT385299_ChiLCB_IN__LKO08_Sal_spl_20
KJ700655_ChiLCB_IN_RKB2_Pet_14
MZ540909_ChiLCB_IN_Gkp_Chi_21
HM143904_ChiLCB_IN_Pan4_Pap_08
HM143901_ToLCB_IN_Pan1_Pap_08
JN663849_PaLCB_IN__Kpnl3_Chi_08
HM007118_ToLCBDB_IN_pChPatnBK19_Chi_08
HM143905_ToLCB_IN_Pan5_Pap_08
KM880104_ToLCBDB_IN_Ahm_Chi_14
MK087125_ToLCBDB_IN_FB1_FB_08
MH577022_ToLCBDB_IN_pTAM1_Tom_09
HM143911_ToLCB_IN_Naj_2_Pap_08
HM143910_ToLCB_IN_DU_Pap_09
JN663875_ToLCBDB_IN_Kpnl6_FB_Chi_08
MF155644_ToLCB_IN_ND_Chi_17
JN663869_PaLCB_IN_Kpnl4_Chi_10
JN663868_PaLCB_IN_Kpnl3_Chi_10
MH577024_ToLCBDB_Ban_pTAJa30_Tom_16
LT827057_ToLCBDB_IN_IS12_Weed_16
MT861129_ToLCBDB_IN_SPUR1_BP_15
DQ343289_ChiLCB_IN_Lko_05
JQ654464_ToLCBDB_IN_HJP01_MB_11
KR957354_ToLCBDB_IN_VIRO_765_Chi_09
Pairwise identity (%)
100
92
84
77
69
61
53
45
38
30
22

c)

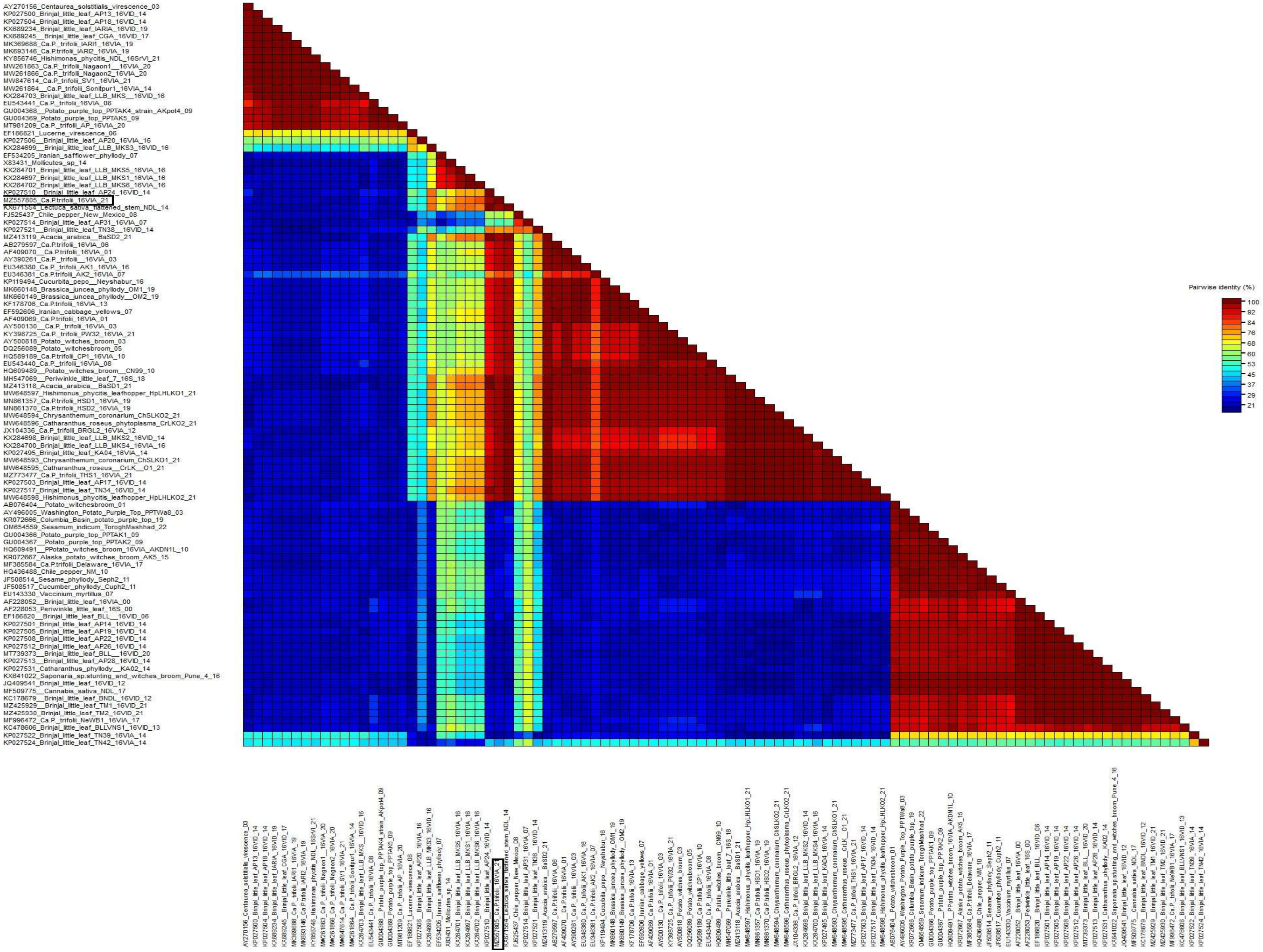
Pairwise identity (%)
100
92
84
76
68
60
53
45
37
29
21
AY270156_Centaurea_solstitialis_virescence_03
KP027500_Brinjal_little_leaf_AP13_16VID_14
KP027504_Brinjal_little_leaf_AP18_16VID_14
KX689234_Brinjal_little_leaf_IARIA_16VID_19
KX689245__Brinjal_little_leaf_CGA_16VID_17
MK369688_Ca.P._trifolii_IARI1_16VIA_19
MK693146_Ca.P.trifolii_IARI2_16VIA_19
KY856746_Hishimonas_phycitis_NDL_16SrVI_21
MW261863_Ca.P._trifolii_Nagaon1__16VIA_20
MW261866_Ca.P._trifolii_Nagaon2_16VIA_20
MW847614_Ca.P._trifolii_SV1_16VIA_21
MW261864__Ca.P._trifolii_Sonitpur1_16VIA_14
KX284703_Brinjal_little_leaf_LLB_MKS__16VID_16
EU543441_Ca.P._trifolii_16VIA_08
GU004368__Potato_purple_top_PPTAK4_strain_AKpot4_09
GU004369_Potato_purple_top_PPTAK5_09
MT981209_Ca.P._trifolii_AP_16VIA_20
EF186821_Lucerne_virescence_06
KP027506__Brinjal_little_leaf_AP20_16VIA_16
KX284699__Brinjal_little_leaf_LLB_MKS3_16VID_16
EF534205_Iranian_safflower_phyllody_07
X83431_Mollicutes_sp_14
KX284701_Brinjal_little_leaf_LLB_MKS5_16VIA_16
KX284697_Brinjal_little_leaf_LLB_MKS1_16VIA_16
KX284702_Brinjal_little_leaf_LLB_MKS6_16VIA_16
KP027510__Brinjal_little_leaf_AP24_16VID_14
MZ557805_Ca.P.trifolii_16VIA_21
KX671554_Lectuca_sativa_flattened_stem_NDL_14
FJ525437_Chile_pepper_New_Mexico_08
KP027514_Brinjal_little_leaf_AP31_16VIA_07
KP027521__Brinjal_little_leaf_TN38__16VID_14
MZ413119_Acacia_arabica__BaSD2_21
AB279597_Ca.P.trifolii_16VIA_06
AF409070__Ca.P.trifolii_16VIA_01
AY390261_Ca.P._trifolii__16VIA_03
EU346380_Ca.P._trifolii_AK1_16VIA_16
EU346381_Ca.P.trifolii_AK2_16VIA_07
KP119494_Cucurbita_pepo__Neyshabur_16
MK660148_Brassica_juncea_phyllody_OM1_19
MK660149_Brassica_juncea_phyllody__OM2_19
KF178706_Ca.P.trifolii_16VIA_13
EF592606_Iranian_cabbage_yellows_07
AF409069_Ca.P.trifolii_16VIA_01
AY500130__Ca.P._trifolii_16VIA_03
KY398725_Ca.P._trifolii_PW32_16VIA_21
AY500818_Potato_witches_broom_03
DQ256089_Potato_witchesbroom_05
HQ589189_Ca.P.trifolii_CP1_16VIA_10
EU543440_Ca.P._trifolii_16VIA_08
HQ609489__Potato_witches_broom__CN99_10
MH547069__Periwinkle_little_leaf_7_16S_18
MZ413118_Acacia_arabica__BaSD1_21
MW648597_Hishimonus_phycitis_leafhopper_HpLHLKO1_21
MN861357_Ca.P.trifolii_HSD1_16VIA_19
MN861370_Ca.P.trifolii_HSD2_16VIA_19
MW648594_Chrysanthemum_coronarium_ChSLKO2_21
MW648596_Catharanthus_roseus_phytoplasma_CrLKO2_21
JX104336_Ca.P._trifolii_BRGL2_16VIA_12
KX284698_Brinjal_little_leaf_LLB_MKS2_16VID_14
KX284700_Brinjal_little_leaf_LLB_MKS4_16VIA_16
KP027495_Brinjal_little_leaf_KA04_16VIA_14
MW648593_Chrysanthemum_coronarium_ChSLKO1_21
MW648595_Catharanthus_roseus__CrLK__O1_21
MZ773477_Ca.P._trifolii_THS1_16VIA_21
KP027503_Brinjal_little_leaf_AP17_16VID_14
KP027517_Brinjal_little_leaf_TN34_16VID_14
MW648598_Hishimonus_phycitis_leafhopper_HpLHLKO2_21
AB076404__Potato_witchesbroom_01
AY496005_Washington_Potato_Purple_Top_PPTWa8_03
KR072666_Columbia_Basin_potato_purple_top_19
OM654559_Sesamum_indicum_ToroghMashhad_22
GU004366_Potato_purple_top_PPTAK1_09
GU004367__Potato_purple_top_PPTAK2_09
HQ609491__PPotato_witches_broom_16VIA_AKDN1L_10
KR072667_Alaska_potato_witches_broom_AK5_15
MF385584_Ca.P.trifolii_Delaware_16VIA_17
HQ436488_Chile_pepper_NM_10
JF508514_Sesame_phyllody_Seph2_11
JF508517_Cucumber_phyllody_Cuph2_11
EU143330_Vaccinium_myrtillus_07
AF228052__Brinjal_little_leaf_16VIA_00
AF228053_Periwinkle_little_leaf_16S_00
EF186820__Brinjal_little_leaf_BLL__16VID_06
KP027501_Brinjal_little_leaf_AP14_16VID_14
KP027505_Brinjal_little_leaf_AP19_16VID_14
KP027508_Brinjal_little_leaf_AP22_16VID_14
KP027512_Brinjal_little_leaf_AP26_16VID_14
MT739373__Brinjal_little_leaf_BLL__16VID_20
KP027513__Brinjal_little_leaf_AP28_16VID_14
KP027531_Catharanthus_phyllody__KA02_14
KX641022_Saponaria_sp.stunting_and_witches_broom_Pune_4_16
JQ409541_Brinjal_little_leaf_16VID_12
MF509775__Cannabis_sativa_NDL_17
KC178679__Brinjal_little_leaf_BNDL_16VID_12
MZ425929__Brinjal_little_leaf_TM1_16VID_21
MZ425930_Brinjal_little_leaf_TM2_16VID_21
MF996472_Ca.P._trifolii_NeWB1_16VIA_17
KC478606_Brinjal_little_leaf_BLLVNS1_16VID_13
KP027522_Brinjal_little_leaf_TN39_16VIA_14
KP027524_Brinjal_little_leaf_TN42_16VIA_14

**d)**

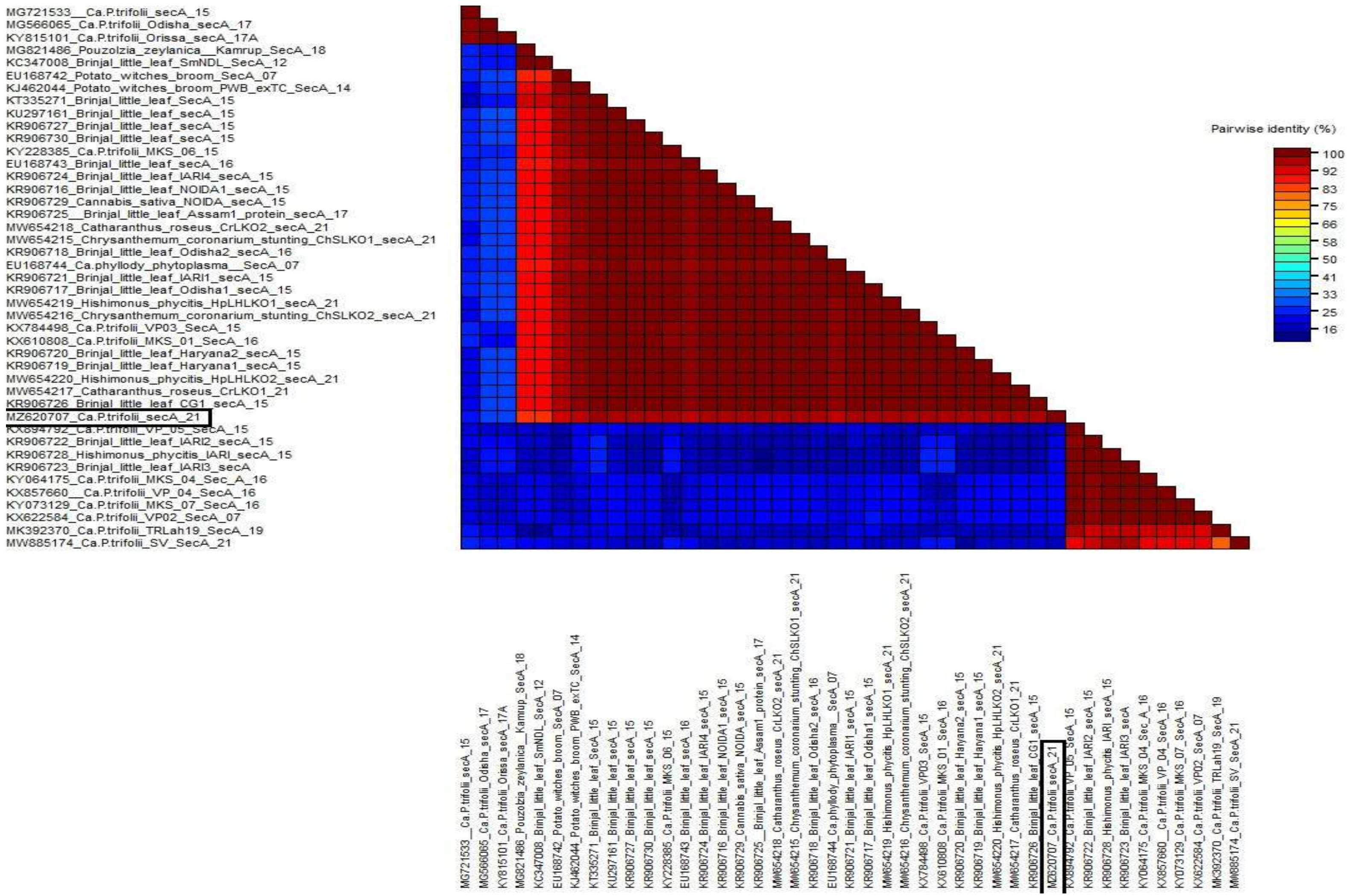


**Figure S1** Using the Sequence Demarcation Tool (SDT v1.2) (Muhire et al., 2014) and the MUSCLE approach, the amino acid sequence identity of the subsequent samples was determined (all of this has been highlighted in red): a) ChiLCV-A-GKP (Acc. No. MZ540908); b) ChLCuB-GKP (Acc. No. MZ540909); c) Ca. P.trifolii16S rRNA (Acc. No. MZ557805); and d) Ca.P.trifolii Sec A gene (Acc. No. MZ620707).
